# Evaluating Temporal Genomic Data for Detecting and Characterizing Balancing Selection

**DOI:** 10.64898/2026.11.12.766181

**Authors:** Ozgur Taskent, Nureda Kocaaydın, Gülşah Merve Kılınç, Matteo Fumagalli, Michael DeGiorgio, Mehmet Somel

## Abstract

Balancing selection encompasses a diverse set of evolutionary mechanisms that maintain genetic polymorphism within populations. However, both detecting balancing selection and distinguishing among its underlying mechanisms remain challenging. Recent advances in the generation of temporal genetic data and the application of machine learning offer new opportunities to address these challenges. In this study, we tested whether temporal genetic data improve the detection of balancing selection and the discrimination between two major balancing selection mechanisms, overdominance and negative frequency-dependent selection. Using forward simulations, we summarized temporal genetic data with features derived from a diverse set of population genetic summary statistics and trained support vector machine and random forest classifiers. Temporal data substantially improved the discrimination of balancing selection from neutral evolution, particularly when summarized by the first four moments of each statistic across time, increasing accuracy by up to 10%. Among individual summary statistics, simple measures of genetic diversity unexpectedly outperformed several more sophisticated statistics. In contrast, summary statistic-based approaches remained insufficient for effectively distinguishing overdominance from negative frequency-dependent selection, even when temporal information was incorporated. These results demonstrate the value of temporal genetic data for detecting balancing selection while highlighting the limitations of summary statistic-based inference for resolving alternative balancing selection mechanisms.

## Introduction

Balancing selection is a mode of natural selection that maintains genetic diversity within populations by favoring the persistence of multiple alleles at a locus. Under balancing selection, the time to the most recent common ancestor (TMRCA) of allelic lineages tends to increase, which results in an accumulation of polymorphisms and a greater proportion of intermediate-frequency alleles in genomic regions linked to the balanced locus (Bitarello et al. 2023). In cases where the TMRCA of the balanced alleles predates species divergence, trans-species polymorphisms may also arise (Leffler et al. 2013). On the other hand, balancing selection can be recent or transient, in which case the genomic signature is often similar to that of an incomplete selective sweep driven by positive selection (Sellis et al. 2011; Isildak et al. 2021).

Historically, the role of balancing selection in shaping overall genetic variation within populations has been contentious. In early work, the prevalence of balancing selection was often debated until the emergence of the neutral theory in the 1960s, which proposed that most polymorphisms are neutral or nearly neutral (Kimura 1968; Ohta 2013). Over time, consensus shifted toward viewing balancing selection as a relatively minor force in maintaining polymorphism. Meanwhile, early genetic scans identified only a limited number of robust cases of balancing selection (Kreitman and Hudson 1991; Andrés et al. 2009; Asthana et al. 2005; Hughes and Nei 1988; Charlesworth 2006). Notable examples include the maintenance of the sickle-cell hemoglobin variant HBB-S in sub-Saharan African human populations (Livingstone 1964), self-incompatibility allelic systems in flowering plants (Charlesworth 2006), chromosomal inversion polymorphisms in *Drosophila* (Hoffmann and Rieseberg 2008), plant disease resistance (*R*) genes (Tian et al. 2002; Bergelson et al. 2001), and the extraordinary diversity of genes in the major histocompatibility complex (MHC) region in vertebrates (Spurgin and Richardson 2010).

Following the publication of the human draft genome in 2001, researchers started genome-wide scans for positive as well as balancing selection, which revealed a limited set of candidate loci showing signatures of balancing selection (e.g., Andrés et al. 2009). Evidence from these scans and locus-specific studies has frequently implicated genes involved in immunity and host-pathogen interactions, including human leukocyte antigen (HLA) genes from the MHC region (Prugnolle et al. 2005; Andrés et al. 2009; Hill et al. 1991), *KIR3DL1/S1* (Norman et al. 2007), the cis-regulatory region of *CCR5* (Bamshad et al. 2002), *DEFB1* (Cagliani et al. 2008), interleukin genes (Fumagalli, Pozzoli, et al. 2009), *ERAP1/2* (Cagliani et al. 2010; Andrés et al. 2010), *ZC3HAV1* (Cagliani et al. 2012), *MEFV* (Fumagalli et al. 2009), *IL10* (Wilson et al. 2006), *LCE3B-C* (Pajic et al. 2016), and *G6PD (*Ruwende et al. 1995; Tishkoff et al. 2001; Verrelli et al. 2002). The ABO blood-group locus also presents signatures of balancing selection, likely due to its key role in mediating pathogen entry into host cells (Fumagalli et al. 2009; Leffler et al. 2013; Ségurel et al. 2012). Likewise, genes involved in cellular adhesion and extracellular components were detected as under balancing selection (e.g., *LAD1*; Teixeira et al. 2015). In addition to immune-related genes, those taking part in reproduction (sperm-expressed genes), such as *ZAN* (Gasper and Swanson 2006), *PKDREJ*, *ZNF473* (Andrés et al. 2009; de Filippo et al. 2016), and *OAZ3* (Christensen et al. 2006), were found as carrying signatures of balancing selection.

Many of these early scans relied on classical population genetic statistics such as Tajima’s *D* (Tajima 1989) and the Hudson-Kreitman-Aguadé (HKA) test (Hudson et al. 1987). Tajima’s *D* detects departures in the site frequency spectrum, specifically, an excess of intermediate-frequency alleles, from the expectation under a constant-size, neutrally-evolving population, whereas the HKA test compares polymorphism and divergence at a locus to neutral or reference expectations, enabling detection of regions with elevated polymorphism given a certain divergence rate. These complementary approaches exploit two distinct signatures of balancing selection. Notwithstanding their utility, Tajima’s *D* is strongly affected by demographic processes (Simonsen et al. 1995), and the HKA test alone does not directly exploit the skew in allele frequencies toward intermediate values (Innan 2006; Nielsen 2005). Recent simulation work also demonstrates that measures depending on skews in the site frequency spectrum (SFS), such as Tajima’s *D*, have power primarily to detect signatures of ancient balancing selection (selection acting for >25 *N_e_* generations), whereas haplotype-based statistics derived from extended haplotype homozygosity, such as iHS (Voight et al. 2006), are better suited to detecting balancing selection that emerged only recently but lose power after approximately 0.01*N*_e_ generations (Soni and Jensen 2024).

To overcome these limitations, powerful methods have been recently developed, the majority of which simultaneously leverage divergence data, frequency spectrum anomalies, and linkage disequilibrium patterns. These include coalescent-based *T*-statistics (DeGiorgio et al. 2014; Cheng and DeGiorgio 2019), the *B*-statistics (Cheng and DeGiorgio 2020, 2022), non-central deviation (NCD) statistics (Bitarello et al. 2018), *β* statistics (Siewert and Voight 2017, 2020), and tests based on extensions of the McDonald-Kreitman framework (Soni et al. 2022). These methods were used to identify many additional loci potentially under long-term balancing selection in humans, such as hundreds to thousands of candidate loci reported by Bitarello et al. (2018) and Soni et al. (2022). Aqil et al. (2023) further demonstrated that many large genomic deletions, which are expected to be deleterious and purged especially when they intersect coding regions, have nonetheless been maintained at moderate to high frequencies since the divergence of modern humans and their extinct relatives (i.e., Neanderthals and Denisovans), suggestive of balancing selection.

A variety of evolutionary mechanisms can lead to balancing selection. These include overdominance, negative frequency-dependent selection (NFDS), spatial and temporal variation in selection, antagonistic pleiotropy or evolutionary trade-offs, and segregation distortion balanced by deleterious fitness effects (DeGiorgio et al. 2014). In all these cases, selection preserves polymorphism in the population. However, the conditions required to reach and maintain a true equilibrium state differ across mechanisms. For instance, early theoretical work found that achieving equilibrium under antagonistic selection (which involves trade-offs) can be much more restrictive compared to other mechanisms of balancing selection (Smith and Hoekstra 1980; Connallon and Clark 2012). In contrast, more recent work using Fisher’s geometric model showed that conditions leading to equilibrium are far more permissive, particularly when dominance reversals are present (Connallon and Clark 2014; Ruzicka et al. 2025). Beyond these equilibrium considerations, theoretical work using Fisher’s geometric model suggests that overdominance might also arise transiently during adaptation (Sellis et al. 2011). Supporting this conclusion, a recent study also found that under the adaptive tracking model of evolution, many mutations under positive selection are maintained at intermediate allele frequencies due to continuously shifting environments and thus selection pressures, creating signatures of balancing selection (Song et al. 2025).

Among the specific mechanisms of balancing selection, overdominance is well-characterised, with examples such as sickle-cell anemia in humans (Livingstone 1964). Another mechanism of particular interest is NFDS, in which the fitness of an allele increases as it becomes rarer. NFDS has long been recognised in theoretical models (e.g., Ayala and Campbell 1974; Clarke and Allendorf 1979) and has documented empirical support in humans: for example, HLA genes show allele frequency patterns consistent with NFDS and the ABO locus has been suggested to evolve under NFDS due to interactions with pathogen-mediated selection (Leffler et al. 2013).

Despite these theoretical advances and the increasing body of empirical observations, we may be underpowered to distinguish balancing selection from neutrality using the currently available tools. For example, Brandt et al. (2026) recently showed that even under favorable parameter combinations, with relatively strong selection, long time-scales of maintenance of the balanced allele under overdominance, and intermediate equilibrium frequencies, the state-of-the-art balancing selection tests have power to detect only a small fraction of loci under balancing selection. Furthermore, distinguishing among different mechanisms of balancing selection is even more challenging. This difficulty arises because different balancing selection mechanisms (e.g., overdominance vs. NFDS) can generate similar allele frequency distributions and spatial polymorphism patterns, particularly when selection persists over long timescales (>4*N*_e_ generations; Fijarczyk and Babik 2015).

One possible way to overcome these difficulties is to use datasets that incorporate temporal sampling (Dehasque et al. 2020). With the advances in ancient DNA research, such datasets have become widely available for human populations over the last decade, enabling a finer resolution of allele-frequency trajectories than is possible with present-day data alone. Several studies focused on positive selection demonstrate the advantages of this increased temporal resolution. For instance, Mathieson et al. (2015) used genomic data from 230 ancient individuals spanning the last 8,000 years and showed that selection on the *LCT* gene, encoding the lactase enzyme responsible for milk-digestion, began around 5,000 years ago, placing its onset in the Bronze Age rather than the Neolithic. Using dense temporal sampling, Burger et al. (2020) subsequently provided evidence for continued strong selection on lactase persistence over the last 3,000 years. Simulation studies offer complementary support for the value of temporal genetic data in selection inference. For example, methods employing simulations demonstrate that time-series data can improve the detection of balancing selection (Korfmann et al. 2023; Soni 2026). Alongside advances in aDNA, new powerful methods that integrate time-transect data now enable estimation of both selection strength and timing of selection (Kerner et al. 2023; Laval et al. 2024; Mathieson 2020; Mathieson and Terhorst 2022; He et al. 2023).

Parallel to these methodological advances, machine learning (ML) has become increasingly central in evolutionary inference (DeGiorgio et al. 2026). ML approaches can integrate multiple summary statistics, capture nonlinear relationships, and generalize across complex parameter spaces, making them ideal for distinguishing overlapping evolutionary signals (Korfmann et al. 2023). Approaches such as support vector machines (SVMs), random forests (RFs), and deep learning architectures have been used to classify genomic regions according to selection type or demographic history (Schrider and Kern 2016; Kern and Schrider 2018; Sheehan and Song 2016; Torada et al. 2019; Flagel et al. 2019; Mughal and DeGiorgio 2019; Isildak et al. 2021; Whitehouse and Schrider 2023; Xue et al. 2021; Mahmoudi et al. 2022; Arnab et al. 2025; Quelin et al. 2025; Amin et al. 2026; van den Belt and Alachiotis 2025; Arnab et al. 2026).

In this study, we assess the performance of ML approaches applied to various population genetic summary statistics in detecting balancing selection as well as discriminating between different modes of balancing selection by using time series data through extensive simulations. Using population genetic models with parameters commonly used to describe human history, we consider evolutionary scenarios incorporating both overdominance and NFDS under human-like demographic and recombination parameters. Using a time-transect sampling scheme, analogous to that available in aDNA datasets, we assess whether ML classifiers can distinguish (1) balancing selection from neutrality and (2) overdominance from NFDS. This simulation-based framework enables systematic evaluation of inference power and the resolution of contemporary selection detection methods under realistic, temporally structured evolutionary models.

## Methods

### Simulations

To test whether population genetic summary statistics can discriminate between different modes of balancing selection and between balancing selection and neutral evolution, we performed forward-time population genetic simulations using SLiM v4.0.1 (Haller and Messer 2023). Simulations included scenarios of overdominance, NFDS, and neutral evolution.

We simulated a single 50 kilobase (kb) genomic region with mutation and recombination rates both set to 10⁻⁸ per site per generation. A single population of 10,000 diploid individuals was initialized and allowed to evolve for 100,000 generations as burn-in (Fig. 1). We then introduced a single target mutation at the center of the genomic element (position 25,000). The initial frequency of the target mutation was set to 5 × 10^-5^ (1/20,000) for balancing selection scenarios and to 0.01 for neutral evolution. That is, the target mutation was introduced on 200 different chromosomes (i.e., haplotypes) simultaneously in each neutral evolution simulation. The higher initial frequency in neutral simulations was chosen to reduce the probability of early loss due to genetic drift. Following the introduction of the target mutation, populations were allowed to evolve for an additional 40,480 generations.

**Figure 1.**
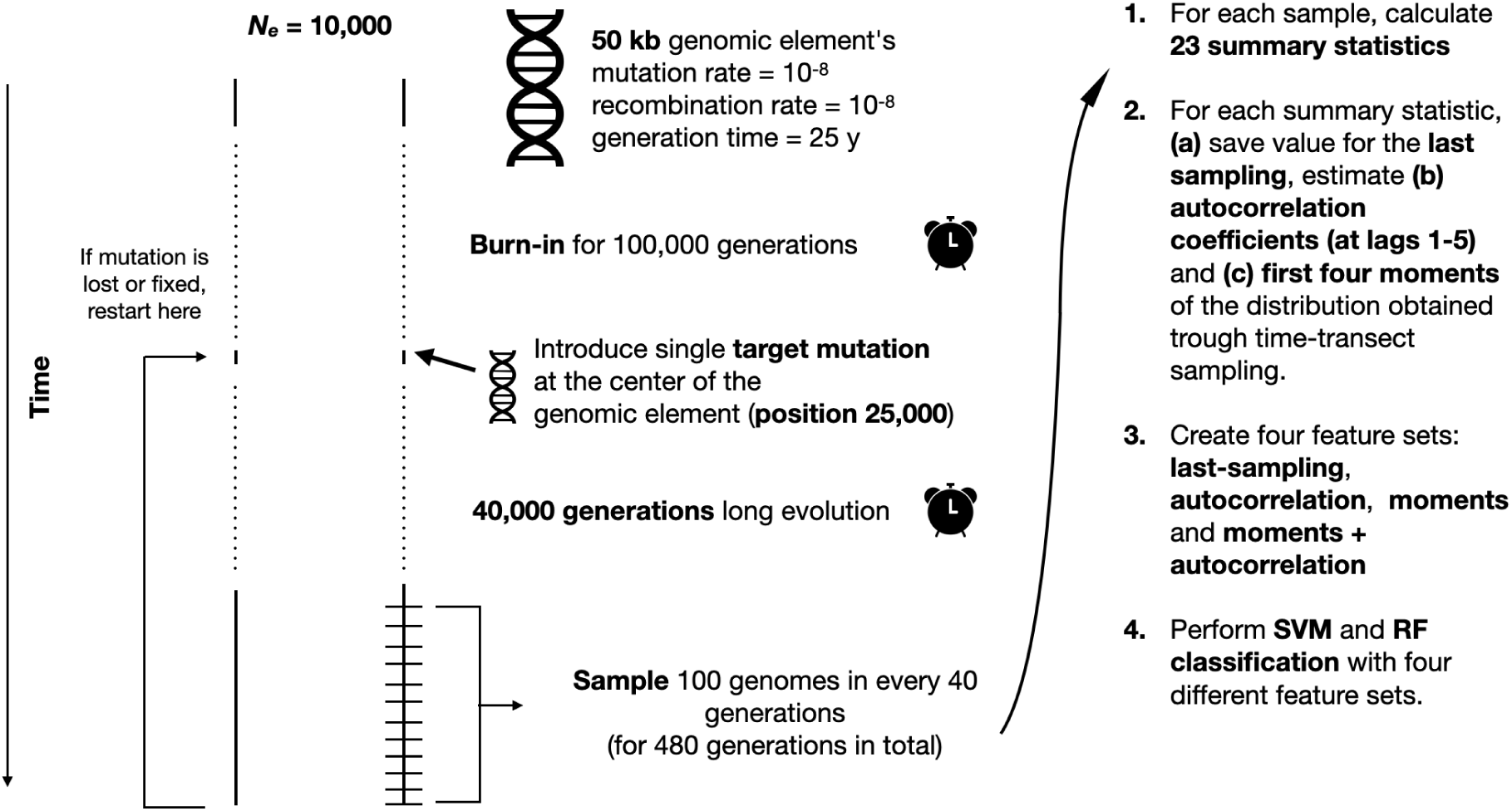
Overview of the simulation workflow. A target mutation was introduced at the center of the genomic element after 100,000 generations. Following 40,000 generations of evolution, 100 genomes are sampled every 40 generations. Summary statistics are calculated for each time point, then analyzed with SVM and RF classification. SVM, support vector machine; RF, random forest.

To ensure the persistence of the target mutation, simulations were conditioned on the allele neither being lost nor fixed during the post-introduction phase. If the mutation was lost or fixed, simulations were restarted from the end of the burn-in period (at generation 100,000). That is, in both balancing selection and neutral evolution scenarios, the target mutation is conditioned to remain segregating in the population. With this framework, we aimed to test whether, among mutations that are segregating for the same time in the population, those under balancing selection can be distinguished from those under neutral evolution.

We note that this target segregation conditioning creates a challenging setting for discriminating balancing selection from neutrality, leading to increased class overlap. Therefore, in a separate set of analyses, we also performed neutral evolution simulations without this conditioning. In total, we conducted four sets of simulations: (i) overdominance simulations with no-fixation-or-loss conditioning (i.e., segregation conditioning), (ii) NFDS simulations with segregation conditioning, and neutral evolution simulations (iii) with and (iv) without segregation conditioning.

Balancing selection simulations were generated under eight selection regimes, spanning weak to strong selection (2*N*_e_*s* ∈ {2, 5, 10, 50, 100, 200, 500, 1,000}). Under balancing selection, the total effect of selection is determined by the compound parameter *hs*, where the dominance (*h*) and selection strength (*s*) parameters of the target mutation were set such that *hs* matched the corresponding *s* values implied by each 2*N*_e_*s* regime. For overdominance, we set the dominance coefficient *h* to a high value to ensure an equilibrium allele frequency of approximately 0.5. Specifically, we set *h* = 20, assumed an effective population size of 10,000, and calculated *s* accordingly. For NFDS simulations, we used the mutationEffect() function in SLiM, which allows genotype fitness to depend dynamically on allele frequency. We defined linear equations such that the fitness of individuals carrying the target mutation decreased with increasing allele frequency. We assumed additive fitness effects by setting the dominance coefficient *h* to 0.5. The equilibrium frequency was set to 0.5 and the mean *s* matched the corresponding overdominance regime when allele frequency was between 0 and 0.5. Specifically, fitness is assigned as 1 + *hs*, where *s* = 2*s_target_* * (1 - 2*f*), *f* denotes the allele frequency of the target mutation and *s_target_* denotes the value of *s* in 2*N*_e_*s* ∈ {2, 5, 10, 50, 100, 200, 500, 1,000}. This linear function ensures *s* = *s_target_*for homozygotes carrying the derived allele at *f* = 0.25, the half point between the frequencies of 0 and 0.5. This choice yields an expectation with a mean selection coefficient comparable to that of the corresponding overdominance regime and allows us to simulate overdominance and NFDS with approximately equivalent fitness effects and comparable allele frequency trajectories (Fig. 2). For neutral evolution with a target mutation, *s* was set to zero.

**Figure 2.**
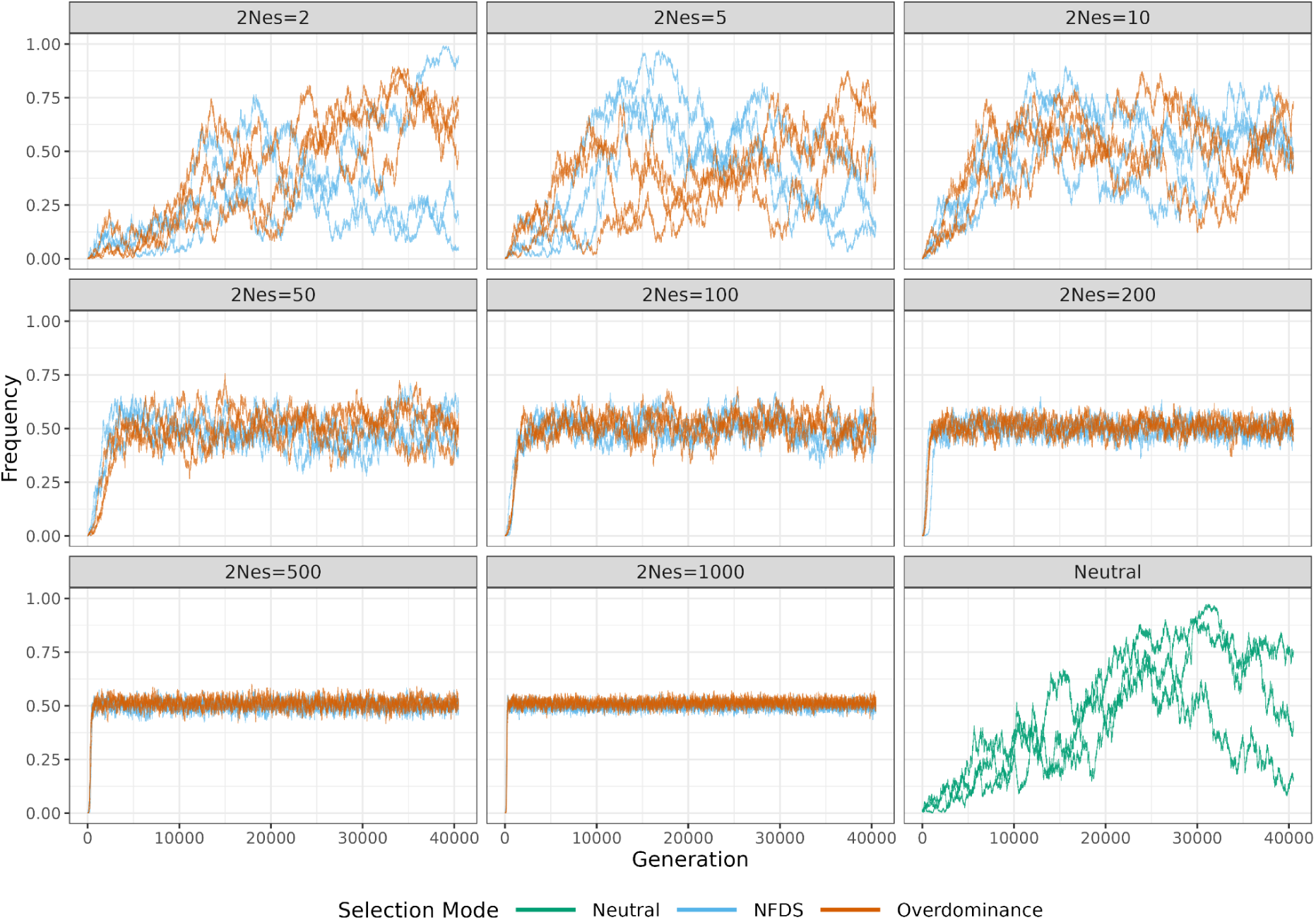
Allele frequency trajectories of the target mutation for three example simulations of overdominance, NFDS and neutral evolution across the total 40,480 simulated generations. Different panels show frequency trajectories for eight different selection strengths (2*N*_e_*s*) used in balancing selection simulations. The simulations were constrained on the variant not being lost or fixed within the last 40,480 generations.

After the introduction of the target mutation, the first 40,000 generations (4*N*_e_) correspond to the TMRCA of neutral alleles in present-day human populations and approximate the lower time bound of long-term balancing selection (Bitarello et al. 2023). During the final 480 generations, corresponding to approximately 12,000 years assuming a generation time of 25 years, we sampled 100 individuals every 40 generations (i.e., every 1,000 years; Fig. 1). This sampling scheme resulted in 13 temporally spaced sampling points and is designed to reflect the availability of human ancient DNA data, the majority of which was sampled from the Holocene Epoch, covering the last 12,000 years [as of 15.04.2026, the Allen Ancient DNA Resource (AADR) data set comprises 17,629 ancient samples, 99.2% (17,494) of which are sampled within the Holocene Epoch, and of these, 67.4% (11,783) are from western Eurasia (Mallick and Reich 2023)].

We performed 1,000 simulations for each balancing selection mode (overdominance and NFDS) and each selection regime, as well as 1,000 simulations for each of two neutral evolution scenarios (with and without segregation conditioning), yielding a total of 18,000 simulations. Owing to stochastic loss or fixation of the target mutation in the segregation-conditioned simulations, achieving this dataset required restarting simulations hundreds of thousands of times (Suppl. Table 1).

### Summary Statistics

For each simulation replicate and each sampling time point, we calculated 23 population genetic summary statistics, including 17 statistics from Isildak et al. (2021), as well as the median allele frequency, trimmed mean allele frequency (allele frequency between 0.1 and 0.9), fraction of intermediate frequency SNPs (allele frequency between 0.3 and 0.7), *β*^(1)^ (Siewert and Voight 2017), and *B*_0,MAF_ (Cheng and DeGiorgio 2020, 2022). The population-genetic summary statistics used in this study are listed in Table 1.

**Table 1.** Summary statistics used in this study.

| **Statistic** | **Information type** | **Reference** |
|---|---|---|
| Tajima’s *D* | SFS/diversity-based | Tajima 1989 |
| Fay and Wu’s *H* | SFS/diversity-based | Fay and Wu 2000 |
| Raggedness Index | SFS/diversity-based | Harpending 1994 |
| Nucleotide Diversity (π) | SFS/diversity-based | Nei and Li 1979 |
| Number of Singletons | SFS/diversity-based |  |
| Mean, Median, and Maximum per-site Mean Pairwise Differences | SFS/diversity-based |  |
| *H*1, *H*12, *H*123, *H*2/*H*1 | Haplotype-based | Garud et al. 2015 |
| Median iHS | Haplotype-based | Voight et al. 2006 |
| Median and Maximum *nS*<sub>L</sub> | Haplotype-based | Ferrer-Admetlla et al. 2014 |
| Number of Haplotypes | Haplotype-based |  |
| Haplotype Diversity | Haplotype-based |  |
| Median Frequency | Frequency-based |  |
| Mean Trimmed Frequency (0.1 < Frequency < 0.9) | Frequency-based |  |
| Fraction of intermediate frequency SNPs (0.3 < Frequency < 0.7; Frac. Inter. Freq.) | Frequency-based |  |
| NCD1 | Balancing selection-specific | Bitarello et al. 2018 |
| *β*<sup>(1)</sup> | Balancing selection-specific | Siewert and Voight 2017 |
| *B*<sub>0,MAF</sub> | Balancing selection-specific | Cheng and DeGiorgio 2022 |

Summary statistics were calculated for a 10 kb window centered on the site where the target mutation is introduced. This window size corresponds approximately to the 95^th^ percentile of the genomic region remaining unrecombined after 40,000 generations assuming a recombination rate of 10^-8^ per site per generation (Siewert and Voight 2020). Calculated statistics included site frequency spectrum–based measures [Tajima’s *D* (Tajima 1989), Fay and Wu’s *H* (Fay and Wu 2000)], haplotype-based statistics [*H*1, *H*12, *H*123, *H*2/*H*1 (Garud et al. 2015), median iHS (Voight et al. 2006), median and maximum *nS*_L_ (Ferrer-Admetlla et al. 2014), haplotype diversity, and the number of unique haplotypes], balancing selection statistics [NCD1 (Bitarello et al. 2018), *β*^(1)^ (Siewert and Voight 2017, 2020), and *B*_0,MAF_ (Cheng and DeGiorgio 2022)], and frequency-based statistics (median allele frequency, trimmed mean allele frequency, and fraction of intermediate frequency SNPs). We also estimated nucleotide diversity π (Nei and Li 1979), the raggedness index (Harpending 1994), per-site mean pairwise nucleotide differences (PWD), and the number of singletons. Mean, median, and maximum values of PWD across all SNPs per window were also included (Table 1), following Isildak et al. (2021).

To compute all summary statistics except *B*_0,MAF_ and *β*^(1)^, we used a modified version of the Python script Preprocessing.py distributed with the BaSe tool (https://github.com/ulasisik/balancing-selection/tree/master/BaSe). To estimate *B*_0,MAF_, we first converted the ms-formatted outputs of SLiM to VCF format with a custom Python script, retaining only the central 10 kb of the 50 kb genomic element. We then used BalLeRMix+ (Cheng and DeGiorgio 2022) tool to parse the input VCF files with the recombination rate parameter set to 10^-8^ per site per generation. The parsed output includes minor allele count data, which can then be used as input for *B*-statistic estimation.

The BalLeRMix+ tool requires a neutral SFS to estimate *B*-statistics. To create this input, we constructed a folded SFS file from the concatenated allele-count data obtained from neutral simulations using a custom Python script. Only the final sampling time point from each neutral simulation replicate was included in the folded SFS. Using this folded SFS file together with the parsed test input, *B*_0,MAF_ was subsequently estimated with BalLeRMix+ using physical genomic positions, no substitution information, and a fixed window size of 10 kb.

To estimate the folded *β*^(1)^ statistic, we used the BetaScan tool (Siewert and Voight 2017, 2020). Similar to BalLeRMix+, BetaScan requires allele-count data as input. We therefore used the parsed allele-count files generated with BalLeRMix+ preprocessing pipeline and estimated *β*^(1)^ with the folded-frequency option, a fixed window size of 10 kb, and a minor-allele-frequency threshold of 0.15.

### Classification

Each simulation produced 13 temporally spaced samples, yielding 13 values for each of the 23 summary statistics. From this data set, we constructed four feature sets. (1) Last sampling: summary statistics estimated for the final sampling time point only, mimicking a situation where only present-day data are available. (2) Time-series moments: the first four moments (mean, standard deviation, skewness, and kurtosis) of each statistic across 13 time points. (3) Time-series autocorrelation: autocorrelation coefficients for each statistic calculated across 13 time points at lags 1-5; that is, for each statistic, autocorrelation coefficients were estimated between values separated by one to five consecutive sampling intervals. (4) Time-series moments with autocorrelation: a combined feature set including both the moments and autocorrelation features. Time-series moments statistics and autocorrelation coefficients were calculated using the respective mean(), sd(), skewness(), kurtosis(), and acf() functions in R (version 4.3.1).

To assess the ability of summary statistics and their time-series derivatives to distinguish (i) balancing selection from neutral evolution and (ii) between different balancing selection modes within an ML context, we applied SVM (Cortes and Vapnik 1995) and RF classifiers (Breiman 2001). For each feature set separately, we first assessed the combined classification performance of all summary statistics and then the performance of each statistic separately. All SVM and RF analyses were performed in R. For RF analyses, we set the number of decision trees at 500, as preliminary analyses indicated that this was sufficient for convergence of model performance, as assessed by the area under the receiver operating characteristic curve (ROC AUC; Suppl. Fig. 1). The minimum node size was set to five observations and the number of predictors considered at each split was set to √*p*, where *p* is the total number of predictors. Analyses were performed with the ranger engine (Wright and Ziegler 2017) in a tidymodels framework (Kuhn and Wickham 2020), where classification trees were constructed by recursively partitioning the data to maximize reductions in Gini impurity (Breiman 2001; Breiman et al. 2017).

SVM analyses were conducted using linear, polynomial, and radial basis function (RBF) kernels. We performed hyperparameter optimization with a grid search for all kernels. Specifically, we explored the following hyperparameters: cost values ranging from 10^-5^ to 10^5^ on a logarithmic scale for all kernels, polynomial degrees 1, 2, and 3 and scale factors ranging from 10^-3^ to 10^-1^ on a logarithmic scale for the polynomial kernel, and σ values ranging from 10^-3^ to 10^-1^ on a logarithmic scale for the RBF kernel. Hyperparameter combinations were generated with a space-filling grid design using the dials and tune R packages and explored with the grid sizes of 20, 60, and 40 for the linear, polynomial, and RBF kernels, respectively. Analyses were conducted with the kernlab engine (Karatzoglou et al. 2004) in a tidymodels framework (Kuhn and Wickham 2020).

For both SVM and RF analyses, each predictor was standardized to have zero mean and unit standard deviation while omitting those with zero variance. This last filtering step was included as a safeguard against rare instances in which variables such as the number of singletons might show zero variance. Model performance was evaluated using stratified 10-fold cross-validation, where the same number of observations from each class are included in each fold, and balanced accuracy and ROC AUC were computed to assess model performance. For each metric, standard errors were calculated from the variation in metric estimates across folds. Because the simulated datasets were largely balanced with respect to class membership, balanced accuracy and overall accuracy were expected to produce similar values. Nevertheless, balanced accuracy was used because the removal of observations containing missing values occasionally resulted in slight class imbalances. The missing values were observed only for the summary statistics trimmed mean frequency and *β*^(1)^ and derived features for fraction of intermediate frequency SNPs. This is because sometimes the 10 kb window did not include any variants within the 0.1 and 0.9 frequency range. As we estimated *β*^(1)^ for variants with >0.15 frequency, this also affected *β*^(1)^ estimation. However, these cases were rare and did not exceed approximately 5% of simulations for any combination of selection mode and selection strength.

For SVM analyses, the models with the hyperparameter combinations having the highest cross-validation ROC AUC scores were chosen as the best models and results of those analyses are presented below. In a separate set of analyses, we assessed the classification performance of each summary statistic individually across the same four feature sets using both RF and SVM classifiers. In SVM classification, we used the linear kernel with a fixed cost hyperparameter set to *C*=1 for this set of analyses.

We also repeated all analyses starting from summary statistic computation to downstream classification tasks with a downsampled data set of *n*=20 haplotypes per sampling point per simulation by choosing 20 haplotypes uniformly at random without replacement among the 200 haplotypes originally sampled by SLiM.

All figures were generated in R using the ggplot2 package (Wickham 2016). The workflow is shown in Fig. 1 and the scripts and pipelines used in all analyses are available in the project’s GitHub repository (https://github.com/taskent/slim-time-transect-balsel-ml).

## Results

We first confirmed that the allele frequencies of the simulated target mutations followed the expected trajectories for each selection regime and selection strength (Fig. 2). The frequency trajectories for balancing selection simulations oscillate around 0.5, with these oscillations covering much narrower frequency ranges for stronger selection regimes, whereas those for the neutral evolution simulations evolve purely stochastically. The number of times the simulations were restarted due to fixation or loss for balancing selection and neutral simulations are given in Suppl. Table 1. In total, we had to repeat the simulations hundreds of thousands of times to collect the dataset.

### Balancing Selection versus Neutral Evolution

We first assessed whether the 23 population-genetic summary statistics computed in this study (Table 1) could reliably distinguish balancing selection from neutral evolution in an ML framework and whether incorporating time-series information improves this performance. To represent balancing selection, here we only used overdominance simulations. Neutral evolution was simulated in two ways: a) conditioning the neutral target mutation to remain segregating in the population for the entire 40,480 generations (i.e., not allowing fixation or loss; see Methods) and b) without such conditioning.

To mimic a scenario in which only present-day data are available, we created a last-sampling feature set that included only the summary statistics calculated at the final (present-day) sampling point (Fig. 1). To exploit the temporal dimension of the data, we constructed two time series-based feature sets: (i) a moments feature set, comprising the first four moments of each summary statistic calculated across the 13 sampling points (spaced 40 generations apart) and (ii) an autocorrelation feature set, comprising autocorrelation coefficients for each summary statistic across the same sampling points at lags 1-5 (Methods). Finally, we created a moments with autocorrelation feature set by combining the features from the moments and autocorrelation feature sets. Summary statistics were calculated using samples of 200 haplotypes per time point.

We first compared balancing selection with neutral evolution conditioned by frequency. We found that classification performance depended on both the learning algorithm and the choice of summary statistics. In RF analyses using the last-sampling feature set, model accuracy was approximately 70-75% for selection strengths 2*N*_e_*s* ≥ 50 (Fig. 3), indicating that these summary statistics have modest classification accuracy to distinguish balancing selection from neutral evolution. Incorporating time-series information into the RF models produced contrasting results. Models based only on the autocorrelation feature set achieved lower accuracy than those based on the last-sampling feature set, with accuracy remaining <65% even for the strongest selection regimes (2*N*_e_*s* ≥ 500; Fig. 3). In contrast, models based on the moments feature set exhibited similar classification accuracy with the last sampling feature set for weak selection regimes and increased accuracy for stronger selection regimes, reaching approximately 83% accuracy for 2*N*_e_*s* ≥ 500 (Fig. 3). These results indicate that incorporating time-series information via moments-based statistics increases accuracy (from ∼1.5% to ∼10% for 2*N*_e_*s* ≥ 50) to distinguish balancing selection from neutral evolution. Adding autocorrelation statistics to the moments feature set did not further improve performance.

**Figure 3.**
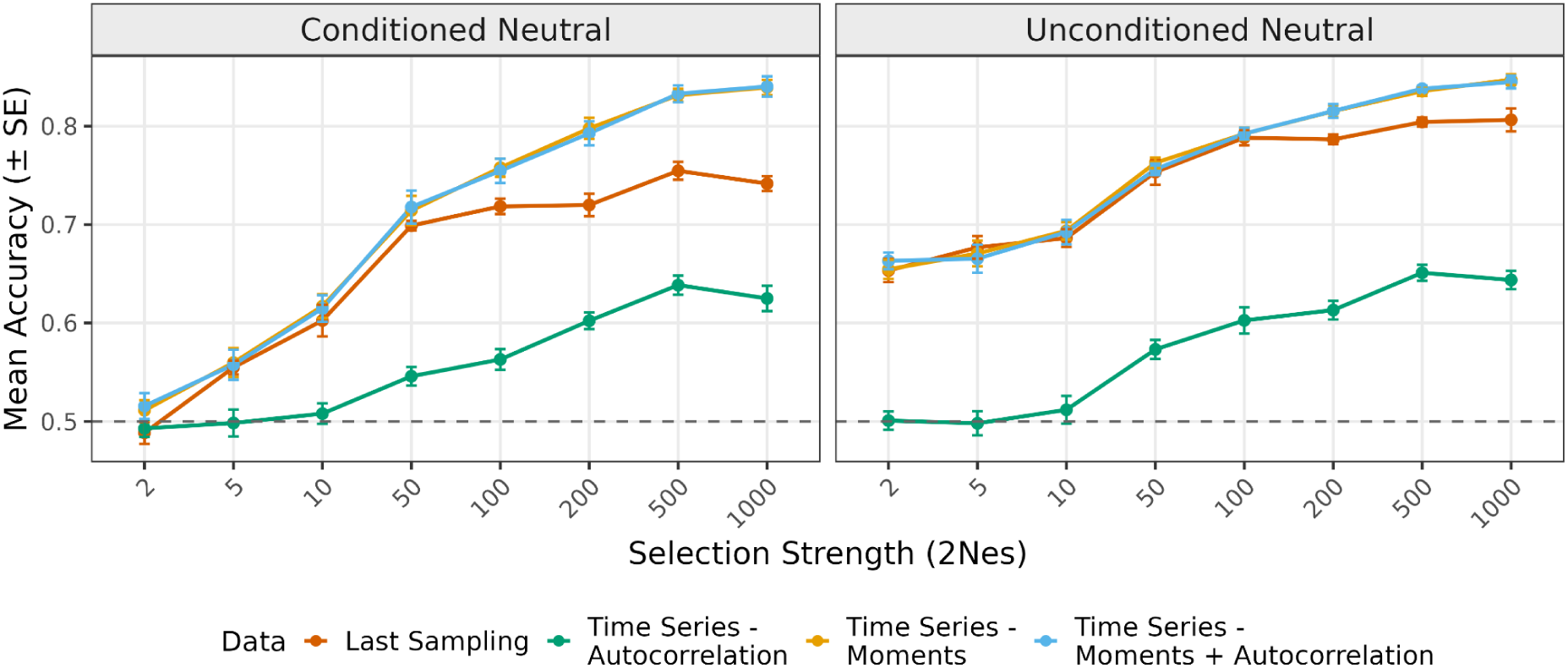
RF classification accuracy for distinguishing balancing selection from neutral evolution as a function of selection strength (2*N*_e_*s*). Models were trained using all 23 summary statistics and four feature sets derived from the same simulations: last sampling, autocorrelation, moments, and moments with autocorrelation. Summary statistics were calculated using *n*=200 haplotypes per sampling point. Points represent mean classification accuracy across the 10 cross-validation folds, with error bars indicating ± SE. The dashed line denotes random expectation (accuracy = 0.5). The Conditioned Neutral panel shows results for neutral simulations in which the target mutation was conditioned to remain segregating for 40,480 generations, whereas the Unconditioned Neutral panel shows results for neutral simulations without this conditioning.

Results of the SVM analyses were qualitatively similar (Suppl. Fig. 2). As in the RF analyses, models based on the autocorrelation statistics exhibited the lowest classification accuracy (in the range of 55-63%). In contrast, time-series moments-based features outperformed last-sampling features under stronger selection, with higher classification accuracy observed for 2*N*_e_*s* ≥ 50 for SVMs with all kernels. Accuracy increased by roughly 1-7% for the linear and polynomial kernels and around 1-5% for the radial kernel (Suppl. Fig. 2). Combining autocorrelation and moments statistics did not further increase classification accuracy (Suppl. Fig. 2). SVM and RF displayed comparable performance across feature sets (Fig. 3, Suppl. Fig. 2) and we report the results of RF analyses in the main text figures. Model performances assessed with ROC AUC were similar to those with accuracy (Suppl. Figs. 5 and 6).

As the sample sizes used in the previous analyses were large (*n* = 200 haplotypes) compared to most available data for natural populations, we decreased sample sizes to *n* = 20 haplotypes per sampling point and repeated RF classification. Downsampling had modest effects on classification performance for all feature sets. The drops in accuracy for moderate to strong selection regimes (2*N*_e_*s* ≥ 50) ranged 4-7%, 1-10%, 1-9%, and 1-9% for the last sampling, autocorrelation, moments, and moments with autocorrelation feature sets, respectively (Suppl. Fig. 3). At 2*N*_e_*s* = 1,000, the model using the moments feature set still achieved 75% accuracy with 20 haplotypes per sampling point, compared with 84% with 200 haplotypes per sampling point (Fig. 3 and Suppl. Fig. 3). Moreover, despite much smaller sample sizes, the gain in performance using the moments features compared to the last sampling features remained almost identical (for 2*N*_e_*s* ≥ 50, 4-8% gain with downsampled data vs. 1.5-10% gain with original data; Suppl. Fig. 3 and Fig. 3).

We then compared balancing selection with neutral simulations without conditioning on segregation. Results of the RF and SVM analyses using these neutral simulations and *n* = 200 showed classification accuracies between approximately 75-85% for all feature sets except autocorrelation across stronger selection regimes (2*N*_e_*s* ≥ 50; Fig. 3, Suppl. Figs. 4). Notably, even under very weak selection (e.g., 2*N*_e_*s* = 2), both RF and SVM achieved moderate classification accuracy (about 65%), which was substantially higher than accuracy using neutral simulations with segregation conditioning (see Discussion). Meanwhile, the relative performance patterns across feature sets were consistent with those observed in segregation-conditioned simulations, with time-series moment-based features improving classification accuracy, whereas autocorrelation-based features provided little to no additional gain. However, the performance gap between the last-sampling and moment-based feature sets was reduced in the absence of segregation conditioning (Fig. 3, Suppl. Figs. 2 and 4). Model performances assessed with ROC AUC were similar to those with accuracy (Suppl. Figs. 7 and 8).

Results of RF classification for downsampled data (*n* = 20) were qualitatively similar. Compared to the last sampling feature set, using the time-series moments feature set led to increases in accuracy for moderate to strong selection regimes (ranging from 1.8% for 2*N*_e_*s* = 50 to 5.3% for 2*N*_e_*s* = 500; Suppl. Fig. 3). For these selection coefficients, the gains in accuracy for downsampled data were even larger compared to the original data (*i.e.*, *n* = 200, where gains were only 0.1% for 2*N*_e_*s* = 50 to 4.1% for 2*N*_e_*s* = 1,000; Fig. 3 and Suppl. Fig. 3). Furthermore, a gain in performance was also observed for weaker selection regimes (2*N*_e_*s* < 50), which was not present using the original data (Fig. 3 and Suppl. Fig. 3). Analogous with the results for the neutral simulations with segregation conditioning, downsampling had modest effects on classification performance of models with all feature sets for moderate to strong selection regimes (2*N*_e_*s* ≥ 50). The drops in accuracy ranged 2-7%, 2-12%, 0.3-6%, and 0.7-6% for the last sampling, autocorrelation, moments, and moments with autocorrelation feature sets, respectively (Suppl. Fig. 3). For the moments feature set and the strongest selection regime (2*N*_e_*s* = 1,000), the model still achieved 78% accuracy (compared to 85% accuracy using *n* = 200; Fig. 3 and Suppl. Fig. 3).

### Classification Performance of Individual Summary Statistics

We then investigated the classification performance of each individual summary statistic. In the RF analyses, 7 statistics individually achieved at least 60% accuracy for the stronger selection regime (i.e., 2*N*_e_*s* ≥ 50) using either the time-series moments or moments with autocorrelation feature sets (Fig. 4). We henceforth refer to this set as the informative statistics. The informative statistics exhibited similar classification accuracies for all selection strengths of 2*N*_e_*s* ≥ 50, indicating that their discriminative performance does not improve substantially after reaching a peak level at around 2*N*_e_*s* = 50 or 100. The only exception to this trend was the maximum of the mean pairwise nucleotide differences across SNPs within the analyzed 10 kb window (Max PWD), whose classification accuracy steadily increased at higher selection strengths, becoming the most accurate statistic for the strongest selection regimes (i.e., 2*N*_e_*s* > 100, Fig. 4; see Discussion). The second best-performing statistic was the fraction of intermediate frequency SNPs (Frac. Inter. Freq.) with around 70% accuracy for moderate to strong selection regimes for the time-series-based moments or moments with autocorrelation feature sets (2*N*_e_*s* ≥ 50; Fig. 4). Statistics based on nucleotide diversity, as well as SFS-based statistics such as π, Tajima’s *D* and Fay & Wu’s *H*, and balancing selection statistics *β*^(1)^, also displayed moderate discriminative performance (classification accuracy ranged from 60% to 70% for selection strengths of 2*N*_e_*s* ≥ 50; Fig. 4). Another balancing selection statistic, NCD1, achieved >60% accuracy only for the stronger selection regimes (2*N*_e_*s* > 50). The remaining statistics, particularly those based on haplotype information, exhibited little to no discriminative ability (Suppl. Figs. 11-14). Results from the SVM analyses were qualitatively similar (Suppl. Figs. 9 and 15-18).

**Figure 4.**
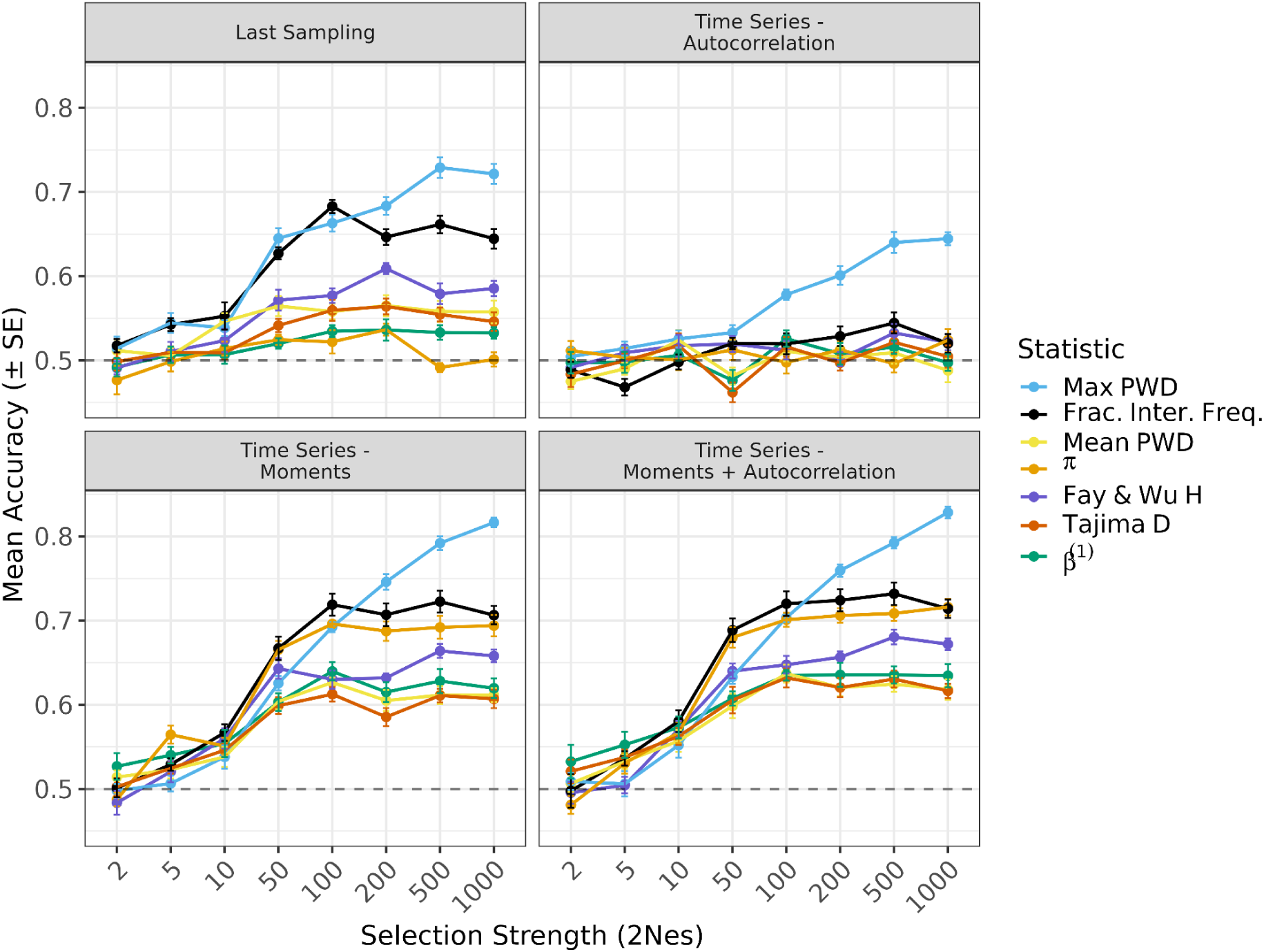
RF classification accuracy of individual summary statistics for discriminating balancing selection from neutral evolution, conditional on no fixation or loss, across selection strengths (2*N*_e_*s*). Summary statistics were calculated using *n*=200 haplotypes per sampling point. Each panel corresponds to a different feature set (last sampling, autocorrelation, moments, and moments with autocorrelation). Points represent mean classification accuracy across the 10 cross-validation folds, with error bars indicating ± SE. Colored lines indicate individual summary statistics. The dashed line denotes random expectation (accuracy = 0.5). Frac. Inter. Freq.: Fraction of intermediate frequency SNPs; Max Het.: Maximum heterozygosity; Mean Het.: Mean heterozygosity; Max PWD: Maximum mean-pairwise-differences; Mean PWD: Mean of mean-pairwise-differences.

For the set of analyses where classifiers were trained using neutral simulations without segregation conditioning, we used a more stringent criteria of classifying a statistic as informative by requiring at least 65% accuracy at 2*N*_e_*s* = 50 using the time-series moments or moments with autocorrelation feature sets, as the models were generally performing better in this setting. Almost all of the informative statistics in the previous analysis emerged as having moderate to high classification accuracies when unconditioned-neutral simulations were used in the analyses (accuracies in the range 70-85% for 2*N*_e_*s* ≥ 50; Fig. 5 and, for the SVM classification, Suppl. Fig. 10). The only exception to this trend was Fay and Wu’s *H*, which surprisingly performed poorly in this setting, with classification accuracies around 60% (Suppl. Fig. 19 and, for the SVM classification, Suppl. Fig. 23). In addition to the aforementioned set of statistics, trimmed mean frequency (i.e., the mean of SNP frequencies in the range 0.1-0.9) and NCD1 also exhibited moderate classification performance for selection strengths 2*N*_e_*s* ≥ 50 (Fig. 5 and, for the SVM classification, Suppl. Fig. 10).

**Figure 5.**
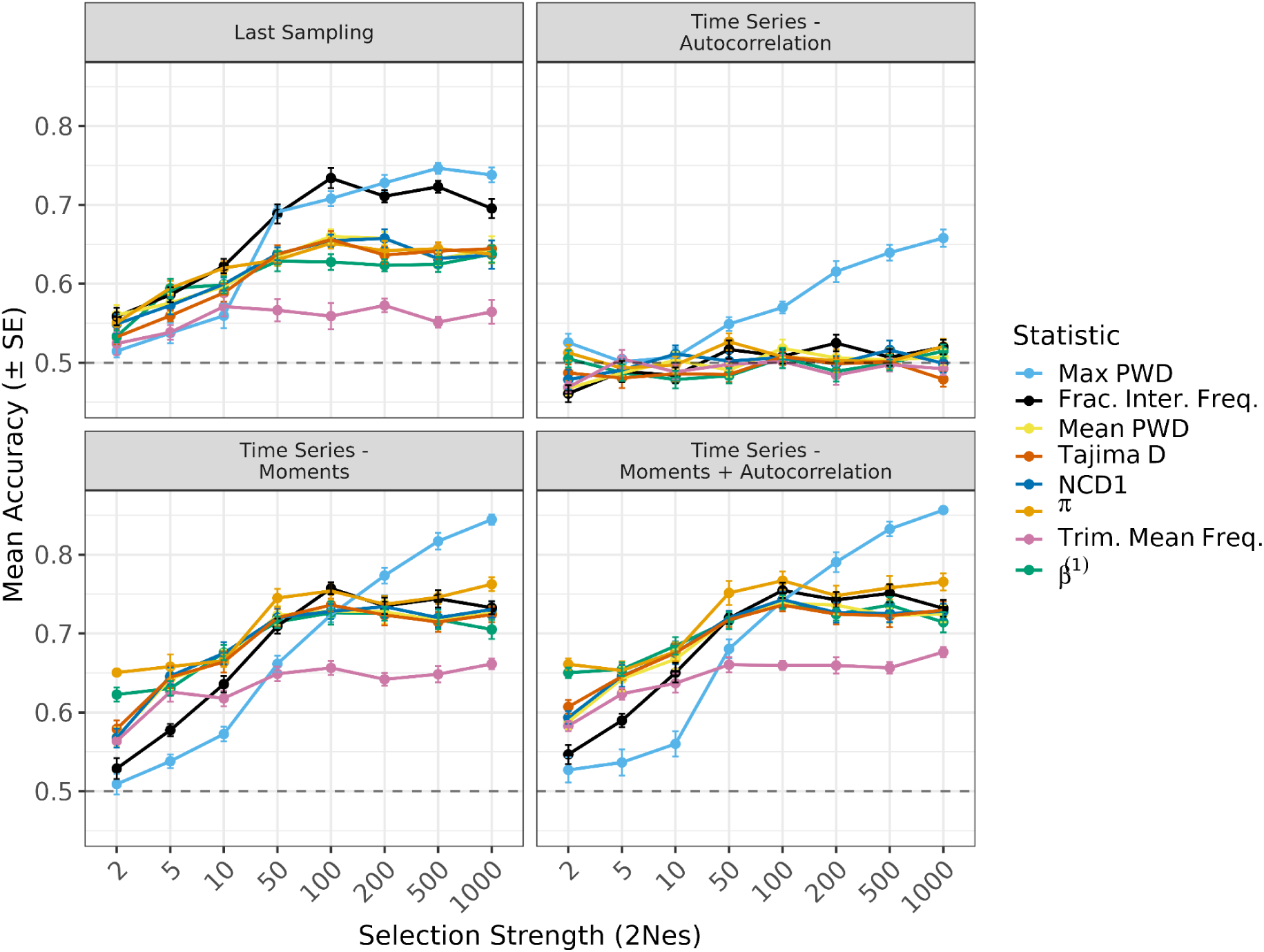
RF classification accuracy of individual summary statistics for discriminating balancing selection from neutral evolution without segregation conditioning across selection strengths (2*N*_e_*s*). Summary statistics were calculated using *n*=200 haplotypes per sampling point. Each panel corresponds to a different feature set (last sampling, autocorrelation, moments, and moments with autocorrelation). Points represent mean classification accuracy across the 10 cross-validation folds, with error bars indicating ± SE. Colored lines indicate individual summary statistics. The dashed line denotes random expectation (accuracy = 0.5). Frac. Inter. Freq.: Fraction of intermediate frequency SNPs; Max Het.: Maximum heterozygosity; Mean Het.: Mean heterozygosity; Max PWD: Maximum mean-pairwise-differences; Mean PWD: Mean of mean-pairwise-differences.

Similar to analyses with segregation conditioned-neutral simulations, Max PWD showed relatively modest performance under weak selection but became the most accurate individual statistic at the strongest selection strengths (2*N*_e_*s* > 100), with a peak accuracy of around 85% for 2*N*_e_*s* = 1,000. All statistics except Max PWD achieved their peak performance at around 2*N*_e_*s* = 50 or 100 and did not exhibit substantial performance gains under stronger selection regimes (Fig. 5 and, for the SVM classification, Suppl. Fig. 10). Individual classification performance of all summary statistics can be found in Suppl. Figs. 19-22 (for the SVM classification, Suppl. Figs. 23-26).

### Overdominance versus NFDS

Overdominance and NFDS are two mechanisms of balancing selection; evidence for both has been documented in natural populations (e.g., Fitzpatrick et al. 2007; Johnston et al. 2013; Gómez-Llano et al. 2024). However, it is challenging to discriminate between these two mechanisms using only present-day genetic data. We reasoned that time-series data, which contains additional information, may help disentangle different balancing selection modes when used together with powerful ML models. Indeed, the results reported above indicate the use of temporal information in discriminating balancing selection from neutral evolution (Fig. 3 and Suppl. Figs. 2-4). Motivated by these observations and similar findings of earlier studies (Soni 2026; Santander et al. 2026), we performed RF and SVM classification to distinguish overdominance from NFDS.

Using the same feature sets (last sampling, autocorrelation, moments, and moments with autocorrelation), we assessed whether population genetic summary statistics could discriminate between overdominance and NFDS. In both RF and SVM analyses, the summary statistics showed very little discriminative power, even at 2*N*_e_*s* = 1,000 (Suppl. Figs. 39 and 40). Results remained similar when we exploited the time-series nature of the data, as classification accuracy did not substantially increase when using the autocorrelation, moments, or moments with autocorrelation feature sets (Suppl. Figs. 39 and 40). Only at 2*N*_e_*s* = 1,000 and when using time-series data was accuracy significantly higher than 50%, but even then, it only reached 53 ± 1%. Model performances assessed with ROC AUC were similar to those with balanced accuracy (Suppl. Figs. 41 and 42). Consistent with these findings, no individual statistic achieved >60% accuracy in either the SVM or RF analyses when evaluated independently (Suppl. Figs. 43-50).

## Discussion

In this study, we systematically evaluated the power of population genetic summary statistics to detect balancing selection and to distinguish between different balancing selection mechanisms using forward-time simulations, time series data, and machine learning approaches, in scenarios compatible with human demographic history and the availability of ancient DNA data from the last 12,000 years. Our results yield several insights into both the strengths and limitations of commonly used statistics and highlight the added value of incorporating temporal information.

First, our results show that the population genetic summary statistics included in this study have moderate to high discriminative ability when used together in an RF or SVM classification set-up to disentangle long-term balancing selection from neutral evolution. When the neutral evolution simulations are conditioned so that the target mutation remains segregating in the population, moderate classification performance is achieved using present-day data only, while classification accuracy increases when temporal information is summarized using moments across sampling time points, particularly under moderate to strong selection regimes (2*N*_e_*s* ≥ 50). The improvement ranges from roughly 1.5% at 2*N*_e_*s* = 50 to about 10% at 2*N*_e_*s* = 1,000. This observation is consistent with the findings of recent studies demonstrating the power of ancient DNA and time-series approaches for selection inference (Mathieson et al. 2015; Dehasque et al. 2020; Mathieson and Terhorst 2022; Kerner et al. 2023; Laval et al. 2024; Irving-Pease et al. 2024; Barrie et al. 2024; Maravall-López et al. 2026).

Second, the classification accuracy increases substantially, especially for weaker selection regimes, when no segregation conditioning is applied to neutral simulations. That is, when the target neutral locus is not conditioned to remain polymorphic in the population, we have more power to detect balancing selection. This finding is expected, as when the aforementioned conditioning is applied, the coalescence time of the lineages at the target neutral locus increases, which leads to the accumulation of polymorphisms in the vicinity of the target neutral locus as well as the degree of differences between the haplotypes carrying the two alleles, effectively mimicking a balancing selection scenario. We note that we evaluated this condition because in empirical data analyses that involve comparisons between candidate variants and neutral variants, researchers may choose to condition the neutral variant set to the similar allele frequencies in order to avoid confounding by technical artifacts or ascertainment.

Surprisingly, we find that even using large sample sizes (200 haplotypes), information across 13 time points, and under strong selection regimes (2*N*_e_*s* = 500 or 1,000), the accuracy of discriminating overdominance from neutrality does not reach 90%, and some of the trajectories (e.g., in Fig. 3) suggest asymptotic behaviour. This result appears to be caused by a subset (∼20%) of neutral loci that tend to be classified as evolving under overdominance (Suppl. Fig. 51). We predict that this misclassification is caused by relatively high stochasticity due to the chosen *N_e_* = 10,000, and that there would be higher power if *N_e_*were higher, which would potentially reduce class overlap compared to the challenging settings that we have simulated.

Third, among the individual summary statistics, the one that captures the maximum of the per-site mean pairwise differences across all segregating sites within the central 10 kb window surrounding the selected mutation (Max PWD) consistently achieves the highest classification accuracy as an RF or SVM classifier, especially under moderate to strong selection regimes and using *n* = 200. Furthermore, while the discriminative abilities of most statistics reach a plateau with increasing selection strengths after moderate selection strengths (e.g., 2*N*_e_*s* = 50), the discriminative ability of Max PWD keeps increasing. These findings likely reflect the fact that stronger balancing selection maintains the selected allele more tightly around its equilibrium frequency of 0.5, thereby increasing the probability that linked polymorphisms are also maintained at intermediate frequencies. Because the mean pairwise difference is maximized at an allele frequency of 0.5, the maximum per-site mean pairwise difference within the region provides an increasingly effective discriminator between balancing selection and neutrality as selection strength increases. For the same reasons, however, the discriminative ability of Max PWD is expected to decrease as equilibrium allele frequency deviates from 0.5, where intermediate-frequency polymorphisms are less likely to occur. Moreover, the classification accuracy of Max PWD depends strongly on sample size. As sample size decreases, its classification performance declines monotonically (Suppl. Fig. 52). This reduction in performance arises because Max PWD is computed using sample allele frequencies and because sampling variance increases with smaller sample sizes, estimation of the maximum per-site mean pairwise differences becomes noisier.

In addition to Max PWD, statistics utilizing SFS- and diversity-based information, such as π, and Tajima’s *D*, exhibit moderate classification performance (accuracies in the range of 59-70% and 72-75% for the moments feature set at 2*N*_e_*s* ≥ 50, with *n* = 200, under neutral simulations with and without segregation conditioning, respectively; Figs. 4 and 5). Frac. Inter. Freq., a frequency-based statistic, is also one of the best-performing predictors. Among the statistics tailored to detect balancing selection, *β*^(1)^ and NCD1 show moderate classification performance in distinguishing balancing selection from neutral evolution. These statistics effectively exploit the hallmark signatures of long-term balancing selection, such as the accumulation of polymorphisms and intermediate frequency alleles in the close vicinity of the balanced locus, which is due to the deep genealogical structure in and around the balanced locus (Hudson and Kaplan 1988; Charlesworth 2006; Bitarello et al. 2018, 2023).

The majority of the summary statistics used in this study retain their discriminative performance with similar classification accuracies using downsampled data (*n* = 20 haplotypes sampled at each sampling point) compared to original data (*n* = 200 haplotypes sampled at each sampling point; Suppl. Figs. 27-38). Counterintuitively, however, some SFS- and diversity-based statistics, such as Tajima’s *D*, Fay and Wu’s *H*, and number of singletons, show slight performance gains with downsampled data for moderate to strong selection regimes (i.e., 2*N*_e_*s* ≥ 50), particularly when the moments features and unconditioned neutral simulations were used in the RF classification (Suppl. Text and Suppl. Figs. 27 and 33). Although the reasons behind these gains in performance were not immediately obvious and thus necessitates further research, we suspect that the increase in sampling variance with downsampling data may differ systematically for overdominance versus neutral evolution when reflected in the SFS, which then aids in separating these two classes in an RF classification set-up.

On the other hand, haplotype-based statistics that depend on extended linkage disequilibrium have little to no discriminative ability in a RF or SVM classification set-up. This result is consistent with the recent findings that such statistics lose power typically within ∼0.01*N*_e_ generations due to the fact that recombination erodes extended haplotype structure around the selected site, leaving little signal for haplotype-based methods to exploit (Soni and Jensen 2024). The power of SFS-based information over haplotype-based information for identifying balancing selection may be considered an advantage in the context of employing time-series data from ancient DNA, because allele frequencies can be calculated directly from ancient DNA sequencing data, while haplotypes can only be inferred indirectly using imputation.

Fourth, the population genetic summary statistics analysed here have effectively no discriminative ability when used in an RF or SVM to distinguish between overdominance and NFDS, at least under the population genetic parameters we explored. The best performance was achieved using time-series data, which supports the idea that temporal information can help distinguish between these two regimes, but even in the best case (at 2*N*_e_*s* = 1,000), accuracy was close to random expectation, suggesting that these two balancing selection mechanisms produce highly similar summary-statistic distributions despite arising from different evolutionary processes.

However, these findings should not be interpreted as evidence that overdominance and NFDS are fundamentally indistinguishable. In recent work utilizing convolutional neural networks with transfer learning, Santander et al. (2026) showed that it is possible to discriminate between overdominance and NFDS with up to around 70% accuracy when only the two classes are considered. Their results therefore suggest that substantially more information is retained when inference is performed directly from genomic data than when it relies on conventional summary statistics. Moreover, Santander et al. (2026) also demonstrated that using ancient alongside present-day samples modestly improves classification accuracy, further highlighting the utility of temporal genomic data in detecting balancing selection and discriminating among its mechanisms.

Our results therefore suggest that the principal limitation lies not in the temporal information itself but in its representation. Deep learning methods that learn feature representations directly from genomic data, rather than relying on predefined summary statistics, may retain spatial and temporal information that is only partially preserved by summary statistic-based approaches. Recent applications of convolutional neural networks to genotype matrices illustrate the promise of this strategy (Santander et al. 2026), while a growing body of work is more broadly demonstrating the utility of deep learning for population genetic inference (Flagel et al. 2019; Torada et al. 2023; Arnab et al. 2025; van den Belt and Alachiotis 2025; Amin et al. 2026; Gower et al. 2021; Mondal et al. 2019; Villanea and Schraiber 2019; Whitehouse and Schrider 2023; Mondal et al. 2025; Laval et al. 2024; Lauterbur et al. 2023; Hamid et al. 2023; Riley et al. 2024; Hejase et al. 2022; Sanchez et al. 2021; Mo and Siepel 2023; Harris et al. 2025; Adrion et al. 2020; Yelmen et al. 2021; Wang et al. 2021).

## Acknowledgements

The lead author used ChatGPT (OpenAI) during the development of this study to assist with programming, code debugging, discussion of methodological approaches, and language editing of the manuscript. All AI-generated suggestions, code, and text were critically evaluated, modified where appropriate, and verified by the authors. All scientific decisions, analyses, interpretations, and conclusions were made by the authors, who take full responsibility for the final manuscript. This study was supported by the Scientific and Technological Research Council of Türkiye (TÜBİTAK) under project number 123C337, as well as grants from the U.S. National Institutes of Health (2R35GM128590), the U.S. National Science Foundation (DEB-2302258), and UKRI Natural Environment Research Council (NE/Y003519/1).

## References

Adrion, Jeffrey R., Jared G. Galloway, and Andrew D. Kern. 2020. “Predicting the Landscape of Recombination Using Deep Learning.” Molecular Biology and Evolution 37 (6): 1790–1808.

Amin, Md Ruhul, Sandipan Paul Arnab, Mohammad Khan, and Michael DeGiorgio. 2026. “Detecting Positive Selection by Modeling Structure within Images of Genetic Variation.” Genome Biology and Evolution 18 (4): evag085.

Andrés, Aida M., Megan Y. Dennis, Warren W. Kretzschmar, et al. 2010. “Balancing Selection Maintains a Form of ERAP2 That Undergoes Nonsense-Mediated Decay and Affects Antigen Presentation.” PLoS Genetics 6 (10): e1001157.

Andrés, Aida M., Melissa J. Hubisz, Amit Indap, et al. 2009. “Targets of Balancing Selection in the Human Genome.” Molecular Biology and Evolution 26 (12): 2755–2764.

Aqil, Alber, Leo Speidel, Pavlos Pavlidis, and Omer Gokcumen. 2023. “Balancing Selection on Genomic Deletion Polymorphisms in Humans.” eLife 12 (January). 10.7554/eLife.79111.

Arnab, Sandipan Paul, Andre Luiz Campelo Dos Santos, Matteo Fumagalli, and Michael DeGiorgio. 2025. “Efficient Detection and Characterization of Targets of Natural Selection Using Transfer Learning.” Molecular Biology and Evolution 42 (5): msaf094.

Arnab, Sandipan Paul, Mohammad Khan, Andre Luiz Campelo Dos Santos, Matteo Fumagalli, and Michael DeGiorgio. 2026. “Identifying Adaptive Footprints in the Presence of Demographic Uncertainty.” Genome Biology and Evolution 18 (4): evag075.

Asthana, Saurabh, Steffen Schmidt, and Shamil Sunyaev. 2005. “A Limited Role for Balancing Selection.” Trends in Genetics: TIG 21 (1): 30–32.

Ayala, Francisco J., and Cathryn A. Campbell. 1974. “Frequency-Dependent Selection.” Annual Review of Ecology and Systematics 5 (1): 115–138.

Bamshad, Michael J., Srinivas Mummidi, Enrique Gonzalez, et al. 2002. “A Strong Signature of Balancing Selection in the 5’ Cis-Regulatory Region of CCR5.” Proceedings of the National Academy of Sciences of the United States of America 99 (16): 10539–10544.

Barrie, William, Yaoling Yang, Evan K. Irving-Pease, et al. 2024. “Elevated Genetic Risk for Multiple Sclerosis Emerged in Steppe Pastoralist Populations.” Nature 625 (7994): 321–328.

Belt, Sjoerd van den, and Nikolaos Alachiotis. 2025. “Fast and Accurate Deep Learning Scans for Signatures of Natural Selection in Genomes Using FASTER-NN.” Communications Biology 8 (1): 58.

Bergelson, J., M. Kreitman, E. A. Stahl, and D. Tian. 2001. “Evolutionary Dynamics of Plant R-Genes.” Science (New York, N.Y.) 292 (5525): 2281–2285.

Bitarello, Bárbara D., Débora Y. C. Brandt, Diogo Meyer, and Aida M. Andrés. 2023. “Inferring Balancing Selection From Genome-Scale Data.” Genome Biology and Evolution 15 (3). 10.1093/gbe/evad032.

Bitarello, Bárbara D., Cesare de Filippo, João C. Teixeira, et al. 2018. “Signatures of Long-Term Balancing Selection in Human Genomes.” Genome Biology and Evolution 10 (3): 939–955.

Brandt, Débora Y. C., Aida M. Andrés, Tim Connallon, and Max Reuter. 2026. “Characterising the Detectable and Invisible Fractions of Genomic Loci under Balancing Selection.” bioRxiv. bioRxiv. 10.64898/2026.01.13.698512.

Breiman, Leo. 2001. “Random Forests.” Machine Learning 45 (1): 5–32.

Breiman, Leo, Jerome H. Friedman, Richard A. Olshen, and Charles J. Stone. 2017. Classification And Regression Trees. 1st Edition. Routledge.

Cagliani, Rachele, Matteo Fumagalli, Stefania Riva, et al. 2008. “The Signature of Long-Standing Balancing Selection at the Human Defensin Beta-1 Promoter.” Genome Biology 9 (9): R143.

Cagliani, Rachele, Stefania Riva, Mara Biasin, et al. 2010. “Genetic Diversity at Endoplasmic Reticulum Aminopeptidases Is Maintained by Balancing Selection and Is Associated with Natural Resistance to HIV-1 Infection.” Human Molecular Genetics 19 (23): 4705–4714.

Cagliani, R., F. R. Guerini, M. Fumagalli, et al. 2012. “A Trans-Specific Polymorphism in ZC3HAV1 Is Maintained by Long-Standing Balancing Selection and May Confer Susceptibility to Multiple Sclerosis.” Molecular Biology and Evolution 29 (6): 1599–1613.

Charlesworth, Deborah. 2006. “Balancing Selection and Its Effects on Sequences in Nearby Genome Regions.” PLoS Genetics 2 (4): e64.

Cheng, Xiaoheng, and Michael DeGiorgio. 2019. “Detection of Shared Balancing Selection in the Absence of Trans-Species Polymorphism.” Molecular Biology and Evolution 36 (1): 177–199.

Cheng, Xiaoheng, and Michael DeGiorgio. 2020. “Flexible Mixture Model Approaches That Accommodate Footprint Size Variability for Robust Detection of Balancing Selection.” Molecular Biology and Evolution 37 (11): 3267–3291.

Cheng, Xiaoheng, and Michael DeGiorgio. 2022. “BalLeRMix +: Mixture Model Approaches for Robust Joint Identification of Both Positive Selection and Long-Term Balancing Selection.” Bioinformatics 38 (3): 861–863.

Christensen, Greg L., Ivaylo P. Ivanov, Stephen P. Wooding, et al. 2006. “Identification of Polymorphisms and Balancing Selection in the Male Infertility Candidate Gene, Ornithine Decarboxylase Antizyme 3.” BMC Medical Genetics 7 (1): 27.

Clarke, B., and F. W. Allendorf. 1979. “Frequency-Dependent Selection due to Kinetic Differences between Allozymes.” Nature 279 (5715): 732–734.

Connallon, Tim, and Andrew G. Clark. 2012. “A General Population Genetic Framework for Antagonistic Selection That Accounts for Demography and Recurrent Mutation.” Genetics 190 (4): 1477–1489.

Connallon, Tim, and Andrew G. Clark. 2014. “Balancing Selection in Species with Separate Sexes: Insights from Fisher’s Geometric Model.” Genetics 197 (3): 991–1006.

Cortes, Corinna, and Vladimir Vapnik. 1995. “Support-Vector Networks.” Machine Learning 20 (3): 273–297.

DeGiorgio, Michael, Sandipan Paul Arnab, and Matteo Fumagalli. 2026. “AI Solutions for Evolutionary Genomics of Nonmodel Species.” Evolution Letters 10 (2): 135–146.

DeGiorgio, Michael, Kirk E. Lohmueller, and Rasmus Nielsen. 2014. “A Model-Based Approach for Identifying Signatures of Ancient Balancing Selection in Genetic Data.” PLoS Genetics 10 (8): e1004561.

Dehasque, Marianne, María C. Ávila-Arcos, David Díez-Del-Molino, et al. 2020. “Inference of Natural Selection from Ancient DNA.” Evolution Letters 4 (2): 94–108.

Fay, J. C., and C. I. Wu. 2000. “Hitchhiking under Positive Darwinian Selection.” Genetics 155 (3): 1405–1413.

Ferrer-Admetlla, Anna, Mason Liang, Thorfinn Korneliussen, and Rasmus Nielsen. 2014. “On Detecting Incomplete Soft or Hard Selective Sweeps Using Haplotype Structure.” Molecular Biology and Evolution 31 (5): 1275–1291.

Fijarczyk, Anna, and Wiesław Babik. 2015. “Detecting Balancing Selection in Genomes: Limits and Prospects.” Molecular Ecology 24 (14): 3529–3545.

Filippo, Cesare de, Felix M. Key, Silvia Ghirotto, et al. 2016. “Recent Selection Changes in Human Genes under Long-Term Balancing Selection.” Molecular Biology and Evolution 33 (6): 1435–1447.

Fitzpatrick, Mark J., Elah Feder, Locke Rowe, and Marla B. Sokolowski. 2007. “Maintaining a Behaviour Polymorphism by Frequency-Dependent Selection on a Single Gene.” Nature 447 (7141): 210–12.

Flagel, Lex, Yaniv Brandvain, and Daniel R. Schrider. 2019. “The Unreasonable Effectiveness of Convolutional Neural Networks in Population Genetic Inference.” Molecular Biology and Evolution 36 (2): 220–238.

Fumagalli, Matteo, Rachele Cagliani, Uberto Pozzoli, et al. 2009. “Widespread Balancing Selection and Pathogen-Driven Selection at Blood Group Antigen Genes.” Genome Research 19 (2): 199–212.

Fumagalli, Matteo, Uberto Pozzoli, Rachele Cagliani, et al. 2009. “Parasites Represent a Major Selective Force for Interleukin Genes and Shape the Genetic Predisposition to Autoimmune Conditions.” The Journal of Experimental Medicine 206 (6): 1395–1408.

Fumagalli, M., R. Cagliani, U. Pozzoli, et al. 2009. “A Population Genetics Study of the Familial Mediterranean Fever Gene: Evidence of Balancing Selection under an Overdominance Regime.” Genes and Immunity 10 (8): 678–686.

Garud, Nandita R., Philipp W. Messer, Erkan O. Buzbas, and Dmitri A. Petrov. 2015. “Recent Selective Sweeps in North American Drosophila Melanogaster Show Signatures of Soft Sweeps.” PLoS Genetics 11 (2): e1005004.

Gasper, Joe, and Willie J. Swanson. 2006. “Molecular Population Genetics of the Gene Encoding the Human Fertilization Protein Zonadhesin Reveals Rapid Adaptive Evolution.” American Journal of Human Genetics 79 (5): 820–830.

Gómez-Llano, Miguel, Ronald D. Bassar, Erik I. Svensson, Simon P. Tye, and Adam M. Siepielski. 2024. “Meta-Analytical Evidence for Frequency-Dependent Selection across the Tree of Life.” Ecology Letters 27 (8): e14477.

Gower, Graham, Pablo Iáñez Picazo, Matteo Fumagalli, and Fernando Racimo. 2021. “Detecting Adaptive Introgression in Human Evolution Using Convolutional Neural Networks.” eLife 10 (e64669): e64669.

Hamid, Iman, Katharine L. Korunes, Daniel R. Schrider, and Amy Goldberg. 2023. “Localizing Post-Admixture Adaptive Variants with Object Detection on Ancestry-Painted Chromosomes.” Molecular Biology and Evolution 40 (4): msad074.

Harpending, H. C. 1994. “Signature of Ancient Population Growth in a Low-Resolution Mitochondrial DNA Mismatch Distribution.” Human Biology 66 (4): 591–600.

Harris, Mariana, Ziyi Mo, Adam Siepel, and Nandita Garud. 2025. “The Persistence and Loss of Hard Selective Sweeps amid Ancient Human Admixture.” bioRxiv. 10.1101/2025.10.14.682443.

He, Zhangyi, Xiaoyang Dai, Wenyang Lyu, Mark Beaumont, and Feng Yu. 2023. “Estimating Temporally Variable Selection Intensity from Ancient DNA Data.” Molecular Biology and Evolution 40 (3). 10.1093/molbev/msad008.

Hejase, Hussein A., Ziyi Mo, Leonardo Campagna, and Adam Siepel. 2022. “A Deep-Learning Approach for Inference of Selective Sweeps from the Ancestral Recombination Graph.” Molecular Biology and Evolution 39 (1). 10.1093/molbev/msab332.

Hill, A. V., C. E. Allsopp, D. Kwiatkowski, et al. 1991. “Common West African HLA Antigens Are Associated with Protection from Severe Malaria.” Nature 352 (6336): 595–600.

Hoffmann, Ary A., and Loren H. Rieseberg. 2008. “Revisiting the Impact of Inversions in Evolution: From Population Genetic Markers to Drivers of Adaptive Shifts and Speciation?” Annual Review of Ecology, Evolution, and Systematics 39 (1): 21–42.

Hudson, R. R., and N. L. Kaplan. 1988. “The Coalescent Process in Models with Selection and Recombination.” Genetics 120 (3): 831–840.

Hudson, R. R., M. Kreitman, and M. Aguadé. 1987. “A Test of Neutral Molecular Evolution Based on Nucleotide Data.” Genetics 116 (1): 153–159.

Hughes, A. L., and M. Nei. 1988. “Pattern of Nucleotide Substitution at Major Histocompatibility Complex Class I Loci Reveals Overdominant Selection.” Nature 335 (6186): 167–170.

Innan, Hideki. 2006. “Modified Hudson–kreitman–aguadé Test and Two-Dimensional Evaluation of Neutrality Tests.” Genetics 173 (3): 1725–1733.

Irving-Pease, Evan K., Alba Refoyo-Martínez, William Barrie, et al. 2024. “The Selection Landscape and Genetic Legacy of Ancient Eurasians.” Nature 625 (7994): 312–320.

Isildak, Ulas, Alessandro Stella, and Matteo Fumagalli. 2021. “Distinguishing between Recent Balancing Selection and Incomplete Sweep Using Deep Neural Networks.” Molecular Ecology Resources 21 (8): 2706–2718.

Johnston, Susan E., Jacob Gratten, Camillo Berenos, Jill G. Pilkington, Tim H. Clutton-Brock, Josephine M. Pemberton, and Jon Slate. 2013. “Life History Trade-Offs at a Single Locus Maintain Sexually Selected Genetic Variation.” Nature 502 (7469): 93–95.

Karatzoglou, Alexandros, Alex Smola, Kurt Hornik, and Achim Zeileis. 2004. “Kernlab-An S4 Package for Kernel Methods in R.” Journal of Statistical Software 11 (9): 1–20.

Kern, Andrew D., and Daniel R. Schrider. 2018. “DiploS/HIC: An Updated Approach to Classifying Selective Sweeps.” G3 (Bethesda, Md.) 8 (6): 1959–1970.

Kerner, Gaspard, Anna-Lena Neehus, Quentin Philippot, et al. 2023. “Genetic Adaptation to Pathogens and Increased Risk of Inflammatory Disorders in Post-Neolithic Europe.” Cell Genomics 3 (2): 100248.

Kimura, M. 1968. “Evolutionary Rate at the Molecular Level.” Nature 217 (5129): 624–626.

Korfmann, Kevin, Oscar E. Gaggiotti, and Matteo Fumagalli. 2023. “Deep Learning in Population Genetics.” Genome Biology and Evolution 15 (2). 10.1093/gbe/evad008.

Kreitman, M., and R. R. Hudson. 1991. “Inferring the Evolutionary Histories of the Adh and Adh-Dup Loci in Drosophila Melanogaster from Patterns of Polymorphism and Divergence.” Genetics 127 (3): 565–582.

Kuhn, M., and H. Wickham. 2020. Tidymodels: A Collection of Packages for Modeling and Machine Learning Using Tidyverse Principles.

Laval, Guillaume, Etienne Patin, Lluis Quintana-Murci, and Gaspard Kerner. 2024. “Deep Estimation of the Intensity and Timing of Natural Selection from Ancient Genomes.” Molecular Ecology Resources 24 (8): e14015.

Lauterbur, M. Elise, Kasper Munch, and David Enard. 2023. “Versatile Detection of Diverse Selective Sweeps with Flex-Sweep.” Molecular Biology and Evolution 40 (6): msad139.

Leffler, Ellen M., Ziyue Gao, Susanne Pfeifer, et al. 2013. “Multiple Instances of Ancient Balancing Selection Shared between Humans and Chimpanzees.” Science 339 (6127): 1578–1582.

Livingstone, F. B. 1964. “Aspects of the Population Dynamics of the Abnormal Hemoglobin and Glucose-6-Phosphate Dehydrogenase Deficiency Genes.” The American Journal of Human Genetics 16 (4): 435–450.

Mahmoudi, Ali, Jere Koskela, Jerome Kelleher, Yao-Ban Chan, and David Balding. 2022. “Bayesian Inference of Ancestral Recombination Graphs.” PLoS Computational Biology 18 (3): e1009960.

Mallick, Swapan, and David Reich. 2023. “The Allen Ancient DNA Resource (AADR): A Curated Compendium of Ancient Human Genomes.” Harvard Dataverse. 10.7910/DVN/FFIDCW.

Maravall-López, Javier, Buu Truong, Gaspard Kerner, et al. 2026. “Ancient DNA Reveals That Natural Selection Has Upregulated the Immune System over the Last 10,000 Years.” In bioRxivorg. BioRxiv, April 14. 10.64898/2026.04.14.718409.

Mathieson, Iain. 2020. “Estimating Time-Varying Selection Coefficients from Time Series Data of Allele Frequencies.” In bioRxiv. November 18. 10.1101/2020.11.17.387761.

Mathieson, Iain, Iosif Lazaridis, Nadin Rohland, et al. 2015. “Genome-Wide Patterns of Selection in 230 Ancient Eurasians.” Nature 528 (7583): 499–503.

Mathieson, Iain, and Jonathan Terhorst. 2022. “Direct Detection of Natural Selection in Bronze Age Britain.” Genome Research 32 (11-12): 2057–2067.

Mo, Ziyi, and Adam Siepel. 2023. “Domain-Adaptive Neural Networks Improve Supervised Machine Learning Based on Simulated Population Genetic Data.” PLoS Genetics 19 (11): e1011032.

Mondal, Mayukh, Jaume Bertranpetit, and Oscar Lao. 2019. “Approximate Bayesian Computation with Deep Learning Supports a Third Archaic Introgression in Asia and Oceania.” Nature Communications 10 (1): 246.

Mondal, Mayukh, Mathilde André, Ajai K. Pathak, Nicolas Brucato, François-Xavier Ricaut, Mait Metspalu, and Anders Eriksson. 2025. “Resolving out of Africa Event for Papua New Guinean Population Using Neural Network.” Nature Communications 16 (1): 6345.

Mughal, Mehreen R., and Michael DeGiorgio. 2019. “Localizing and Classifying Adaptive Targets with Trend Filtered Regression.” Molecular Biology and Evolution 36 (2): 252–270.

Nei, M., and W. H. Li. 1979. “Mathematical Model for Studying Genetic Variation in Terms of Restriction Endonucleases.” Proceedings of the National Academy of Sciences of the United States of America 76 (10): 5269–5273.

Nielsen, Rasmus. 2005. “Molecular Signatures of Natural Selection.” Annual Review of Genetics 39: 197–218.

Norman, Paul J., Laurent Abi-Rached, Ketevan Gendzekhadze, et al. 2007. “Unusual Selection on the KIR3DL1/S1 Natural Killer Cell Receptor in Africans.” Nature Genetics 39 (9): 1092–1099.

Ohta, Tomoko. 2013. “Molecular Evolution: Nearly Neutral Theory.” In eLS. John Wiley & Sons, Ltd, February 15. 10.1002/9780470015902.a0001801.pub4.

Pajic, Petar, Yen-Lung Lin, Duo Xu, and Omer Gokcumen. 2016. “The Psoriasis-Associated Deletion of Late Cornified Envelope Genes LCE3B and LCE3C Has Been Maintained under Balancing Selection since Human Denisovan Divergence.” BMC Evolutionary Biology 16 (1): 265.

Prugnolle, Franck, Andrea Manica, Marie Charpentier, Jean François Guégan, Vanina Guernier, and François Balloux. 2005. “Pathogen-Driven Selection and Worldwide HLA Class I Diversity.” Current Biology: CB 15 (11): 1022–1027.

Quelin, Arnaud, Frédéric Austerlitz, and Flora Jay. 2025. “Assessing Simulation-Based Supervised Machine Learning for Demographic Parameter Inference from Genomic Data.” Heredity 134 (7): 417–426.

Riley, Rebecca, Iain Mathieson, and Sara Mathieson. 2024. “Interpreting Generative Adversarial Networks to Infer Natural Selection from Genetic Data.” Genetics 226 (4): iyae024.

Ruwende, C., S. C. Khoo, R. W. Snow, et al. 1995. “Natural Selection of Hemi- and Heterozygotes for G6PD Deficiency in Africa by Resistance to Severe Malaria.” Nature 376 (6537): 246–249.

Ruzicka, Filip, Martyna K. Zwoinska, Debora Goedert, et al. 2025. “A Century of Theories of Balancing Selection.” In bioRxiv. February 14. 10.1101/2025.02.12.637871.

Sanchez, Théophile, Jean Cury, Guillaume Charpiat, and Flora Jay. 2021. “Deep Learning for Population Size History Inference: Design, Comparison and Combination with Approximate Bayesian Computation.” Molecular Ecology Resources 21 (8): 2645–60.

Santander, Cindy Gilda, Andre Luiz Campelo Dos Santos, Sandipan Paul Arnab, Matteo Fumagalli, and Michael DeGiorgio. 2026. “Negative Frequency-Dependent Selection: A Positive Outlook with Deep Learning.” Philosophical Transactions of the Royal Society of London. Series B, Biological Sciences 381 (1952): 20250170.

Schrider, Daniel R., and Andrew D. Kern. 2016. “S/HIC: Robust Identification of Soft and Hard Sweeps Using Machine Learning.” PLoS Genetics 12 (3): e1005928.

Ségurel, Laure, Emma E. Thompson, Timothée Flutre, et al. 2012. “The ABO Blood Group Is a Trans-Species Polymorphism in Primates.” Proceedings of the National Academy of Sciences of the United States of America 109 (45): 18493–18498.

Sellis, Diamantis, Benjamin J. Callahan, Dmitri A. Petrov, and Philipp W. Messer. 2011. “Heterozygote Advantage as a Natural Consequence of Adaptation in Diploids.” Proceedings of the National Academy of Sciences of the United States of America 108 (51): 20666–20671.

Sheehan, Sara, and Yun S. Song. 2016. “Deep Learning for Population Genetic Inference.” PLoS Computational Biology 12 (3): e1004845.

Siewert, Katherine M., and Benjamin F. Voight. 2017. “Detecting Long-Term Balancing Selection Using Allele Frequency Correlation.” Molecular Biology and Evolution 34 (11): 2996–3005.

Siewert, Katherine M., and Benjamin F. Voight. 2020. “BetaScan2: Standardized Statistics to Detect Balancing Selection Utilizing Substitution Data.” Genome Biology and Evolution 12 (2): 3873–3877.

Simonsen, K. L., G. A. Churchill, and C. F. Aquadro. 1995. “Properties of Statistical Tests of Neutrality for DNA Polymorphism Data.” Genetics 141 (1): 413–429.

Smith, J. Maynard, and R. Hoekstra. 1980. “Polymorphism in a Varied Environment: How Robust Are the Models?” Genetics Research 35 (1): 45–57.

Song, Siliang, Piaopiao Chen, Xukang Shen, and Jianzhi Zhang. 2025. “Adaptive Tracking with Antagonistic Pleiotropy Results in Seemingly Neutral Molecular Evolution.” Nature Ecology & Evolution 9 (12): 2358–2373.

Soni, Vivak. 2026. “Can Ancient DNA and Other Forms of Time-Sampled Data Aid in the Inference of Negative Frequency-Dependent Selection?” Philosophical Transactions of the Royal Society of London. Series B, Biological Sciences 381 (1952): 20250171.

Soni, Vivak, and Jeffrey D. Jensen. 2024. “Temporal Challenges in Detecting Balancing Selection from Population Genomic Data.” G3, ahead of print, March 29. 10.1093/g3journal/jkae069.

Soni, Vivak, Michiel Vos, and Adam Eyre-Walker. 2022. “A New Test Suggests Hundreds of Amino Acid Polymorphisms in Humans Are Subject to Balancing Selection.” PLoS Biology 20 (6): e3001645.

Spurgin, Lewis G., and David S. Richardson. 2010. “How Pathogens Drive Genetic Diversity: MHC, Mechanisms and Misunderstandings.” Proceedings. Biological Sciences / The Royal Society 277 (1684): 979–988.

Tajima, F. 1989. “Statistical Method for Testing the Neutral Mutation Hypothesis by DNA Polymorphism.” Genetics 123 (3): 585–595.

Tajima, F. 1993. “Statistical Analysis of DNA Polymorphism.” The Japanese Journal of Genetics 68 (6): 567–595.

Teixeira, João C., Cesare de Filippo, Antje Weihmann, et al. 2015. “Long-Term Balancing Selection in LAD1 Maintains a Missense Trans-Species Polymorphism in Humans, Chimpanzees, and Bonobos.” Molecular Biology and Evolution 32 (5): 1186–1196.

Tian, Dacheng, Hitoshi Araki, Eli Stahl, Joy Bergelson, and Martin Kreitman. 2002. “Signature of Balancing Selection in Arabidopsis.” Proceedings of the National Academy of Sciences of the United States of America 99 (17): 11525–11530.

Tishkoff, S. A., R. Varkonyi, N. Cahinhinan, et al. 2001. “Haplotype Diversity and Linkage Disequilibrium at Human G6PD: Recent Origin of Alleles That Confer Malarial Resistance.” Science (New York, N.Y.) 293 (5529): 455–462.

Torada, Luis, Lucrezia Lorenzon, Alice Beddis, et al. 2019. “ImaGene: A Convolutional Neural Network to Quantify Natural Selection from Genomic Data.” BMC Bioinformatics 20 (Suppl 9): 337.

Verrelli, Brian C., John H. McDonald, George Argyropoulos, et al. 2002. “Evidence for Balancing Selection from Nucleotide Sequence Analyses of Human G6PD.” American Journal of Human Genetics 71 (5): 1112–1128.

Villanea, Fernando A., and Joshua G. Schraiber. 2019. “Multiple Episodes of Interbreeding between Neanderthal and Modern Humans.” Nature Ecology & Evolution 3 (1): 39–44.

Voight, Benjamin F., Sridhar Kudaravalli, Xiaoquan Wen, and Jonathan K. Pritchard. 2006. “A Map of Recent Positive Selection in the Human Genome.” PLoS Biology 4 (3): e72.

Wang, Zhanpeng, Jiaping Wang, Michael Kourakos, Nhung Hoang, Hyong Hark Lee, Iain Mathieson, and Sara Mathieson. 2021. “Automatic Inference of Demographic Parameters Using Generative Adversarial Networks.” Molecular Ecology Resources 21 (8): 2689–2705.

Whitehouse, Logan S., and Daniel R. Schrider. 2023. “Timesweeper: Accurately Identifying Selective Sweeps Using Population Genomic Time Series.” Genetics 224 (3). 10.1093/genetics/iyad084.

Wickham, Hadley. 2016. Ggplot2: Elegant Graphics for Data Analysis. 2nd ed. Use R! Springer International Publishing. PDF.

Wilson, J. N., K. Rockett, B. Keating, et al. 2006. “A Hallmark of Balancing Selection Is Present at the Promoter Region of Interleukin 10.” Genes and Immunity 7 (8): 680–683.

Wright, Marvin N., and Andreas Ziegler. 2017. “Ranger: A Fast Implementation of Random Forests for High Dimensional Data in C++ and R.” Journal of Statistical Software 77 (1): 1–17.

Xue, Alexander T., Daniel R. Schrider, Andrew D. Kern, and Ag1000g Consortium. 2021. “Discovery of Ongoing Selective Sweeps within Anopheles Mosquito Populations Using Deep Learning.” Molecular Biology and Evolution 38 (3): 1168–1183.

Yelmen, Burak, Aurélien Decelle, Linda Ongaro, Davide Marnetto, Corentin Tallec, Francesco Montinaro, Cyril Furtlehner, Luca Pagani, and Flora Jay. 2021. “Creating Artificial Human Genomes Using Generative Neural Networks.” PLoS Genetics 17 (2): e1009303.

