## Supplementary Materials for "Evaluating Temporal Genomic Data for Detecting and Characterizing Balancing Selection"

**Supplementary Table 1.** The number of times the simulations were restarted due to fixation or loss of the target mutation

| Selection Mode | Selection Strength ($2N_es$) | Mean | Median | Standard Deviation | Maximum | Minimum |
|---|---|---|---|---|---|---|
| Balancing Selection (Overdominance) | 2 | 10,174.7 | 6,812.5 | 10,555.0 | 87,670.0 | 55.0 |
| | 5 | 3,357.4 | 2,399.0 | 3,313.5 | 26,068.0 | 13.0 |
| | 10 | 1,214.8 | 858.0 | 1,175.7 | 9,381.0 | 1.0 |
| | 50 | 203.0 | 139.5 | 209.1 | 1,589.0 | 0.0 |
| | 100 | 100.8 | 69.0 | 102.6 | 724.0 | 0.0 |
| | 200 | 47.7 | 34.0 | 47.7 | 401.0 | 0.0 |
| | 500 | 20.2 | 14.0 | 20.0 | 172.0 | 0.0 |
| | 1,000 | 9.9 | 6.0 | 10.7 | 85.0 | 0.0 |
| Balancing Selection (NFDS) | 2 | 9,426.5 | 6,872.5 | 8,913.4 | 52,805.0 | 4.0 |
| | 5 | 3,522.9 | 2,512.5 | 3,344.5 | 23,907.0 | 1.0 |
| | 10 | 1,254.4 | 832.5 | 1,214.2 | 8,380.0 | 0.0 |
| | 50 | 192.0 | 135.5 | 189.5 | 1,307.0 | 0.0 |
| | 100 | 97.9 | 66.0 | 101.1 | 869.0 | 0.0 |
| | 200 | 50.1 | 34.5 | 50.8 | 336.0 | 0.0 |
| | 500 | 19.8 | 13.0 | 19.5 | 130.0 | 0.0 |
| | 1000 | 9.6 | 7.0 | 10.5 | 80.0 | 0.0 |
| Neutral Evolution (Initial Frequency = 0.01) | - | 130.9 | 99.0 | 118.8 | 863.0 | 0.0 |

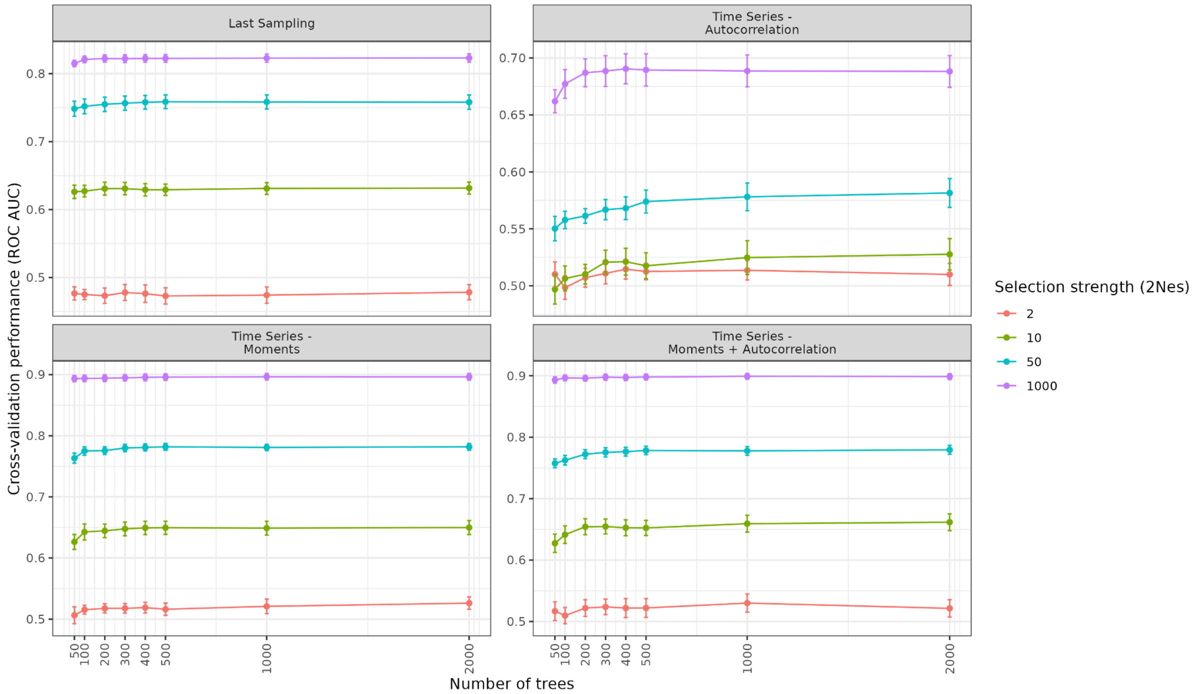


**Supplementary Figure 1.** RF classification performance, measured by ROC AUC, distinguishing balancing selection from neutral evolution as a function of the number of decision trees. Models were trained using all 23 summary statistics and four different feature sets derived from the same simulations: last sampling, autocorrelation, moments, and moments with autocorrelation. Summary statistics were calculated using *n*=200 haplotypes per sampling point. Performance is shown for four representative selection strengths ($2N_es$ = 2, 10, 50, and 1000). Points represent mean ROC AUC across the 10 cross-validation folds, with error bars indicating ± SE.

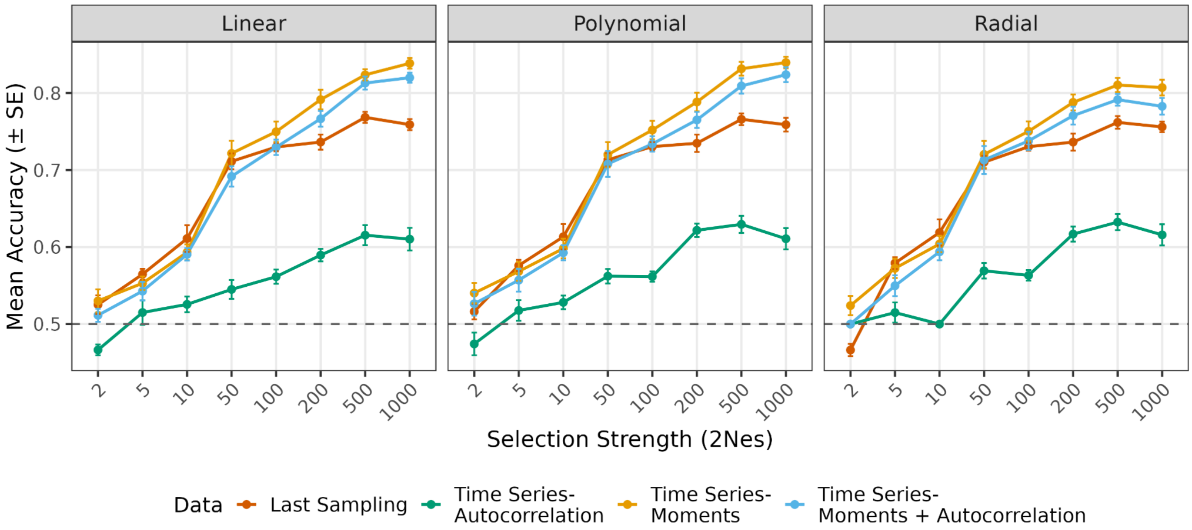


**Supplementary Figure 2.** SVM classification accuracy for distinguishing balancing selection from neutral evolution, conditional on no fixation or loss, as a function of selection strength ($2N_es$). Models were trained using all 23 summary statistics and four different feature sets derived from the same simulations: last sampling, autocorrelation, moments, and moments with autocorrelation. Summary statistics were calculated using $n$=200 haplotypes per sampling point. Points represent mean classification accuracy across the 10 cross-validation folds, with error bars indicating ± SE. Each panel corresponds to SVM analyses with different kernels. The dashed line denotes random expectation (accuracy = 0.5).

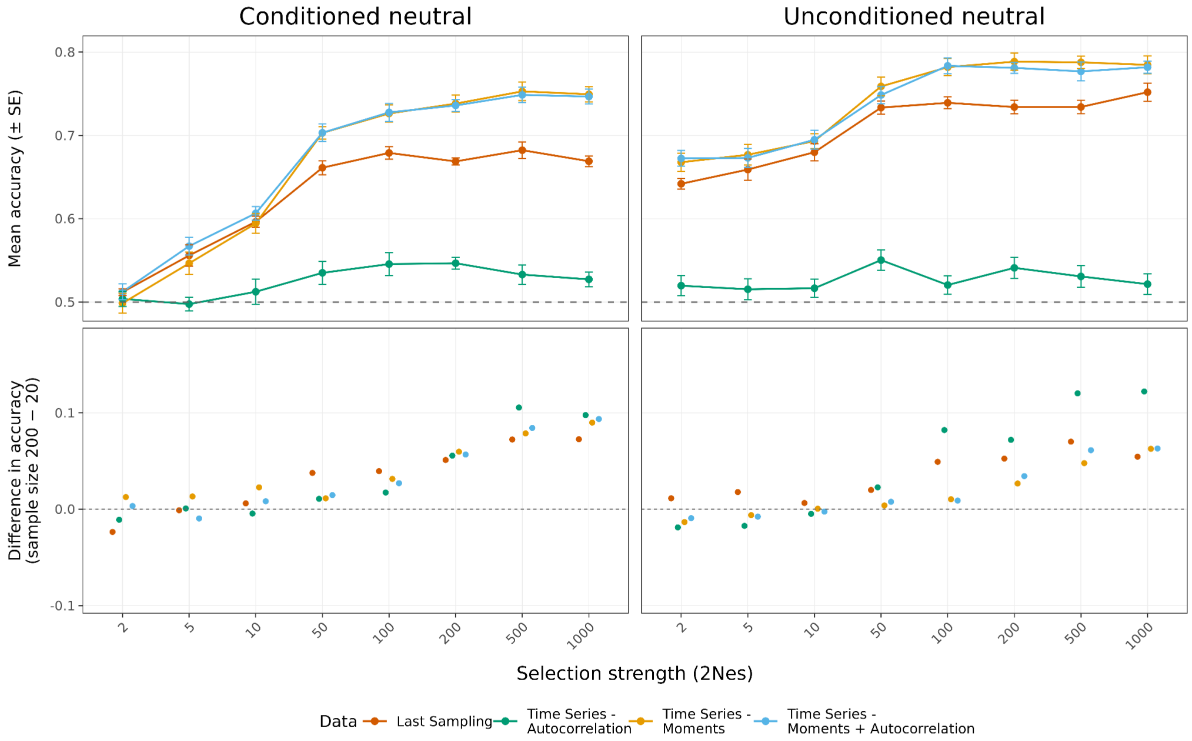


**Supplementary Figure 3.** RF classification accuracy for distinguishing balancing selection from neutral evolution across selection strength (2$N_e$$s$) using downsampled data ($n$=20 haplotypes per sampling point) and accuracy differences of the same models with original ($n$=200 haplotypes per sampling point) versus downsampled data. Models were trained using all 23 summary statistics and four feature sets derived from the same simulations: last sampling, autocorrelation, moments, and moments with autocorrelation. Points represent mean classification accuracy across the 10 cross-validation folds, with error bars indicating ± SE. The dashed line denotes random expectation (accuracy = 0.5). The Conditioned Neutral panel shows results for neutral simulations in which the target mutation was conditioned to remain segregating for 40,480 generations, whereas the Unconditioned Neutral panel shows results for neutral simulations without this conditioning.

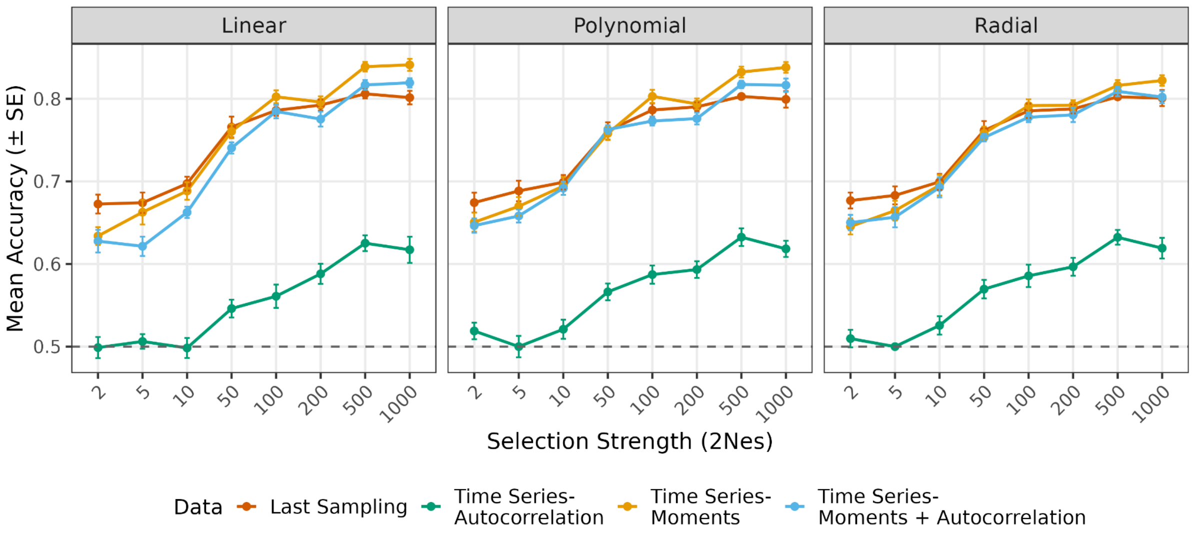


**Supplementary Figure 4.** SVM classification accuracy for distinguishing balancing selection from neutral simulations without segregation conditioning as a function of selection strength ($2N_es$). Models were trained using all 23 summary statistics and four different feature sets derived from the same simulations: last sampling, autocorrelation, moments, and moments with autocorrelation. Summary statistics were calculated using $n$=200 haplotypes per sampling point. Points represent mean classification accuracy across the 10 cross-validation folds, with error bars indicating ± SE. Each panel corresponds to SVM analyses with different kernels. The dashed line denotes random expectation (accuracy = 0.5).

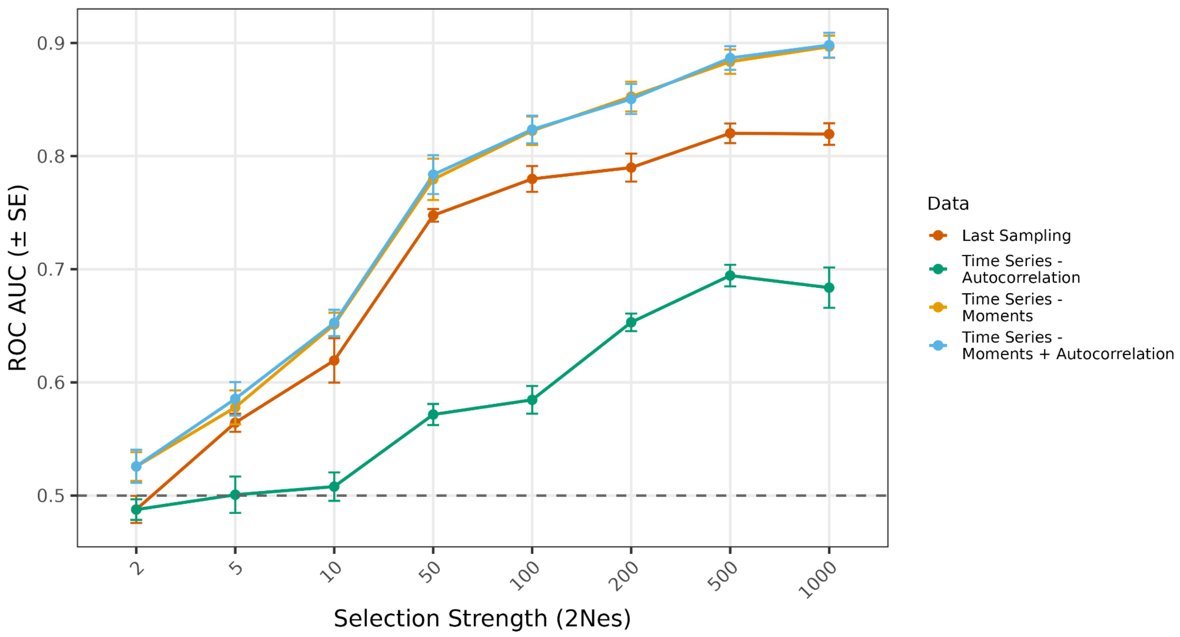


**Supplementary Figure 5.** RF classification performance, measured by ROC AUC, for distinguishing balancing selection from neutral evolution, conditional on no fixation or loss, as a function of selection strength ($2N_es$). Models were trained using all 23 summary statistics and four different feature sets derived from the same simulations: last sampling, autocorrelation, moments, and moments with autocorrelation. Summary statistics were calculated using $n$=200 haplotypes per sampling point. Points represent mean ROC AUC across the 10 cross-validation folds, with error bars indicating ± SE. The dashed line denotes random expectation (ROC AUC = 0.5).

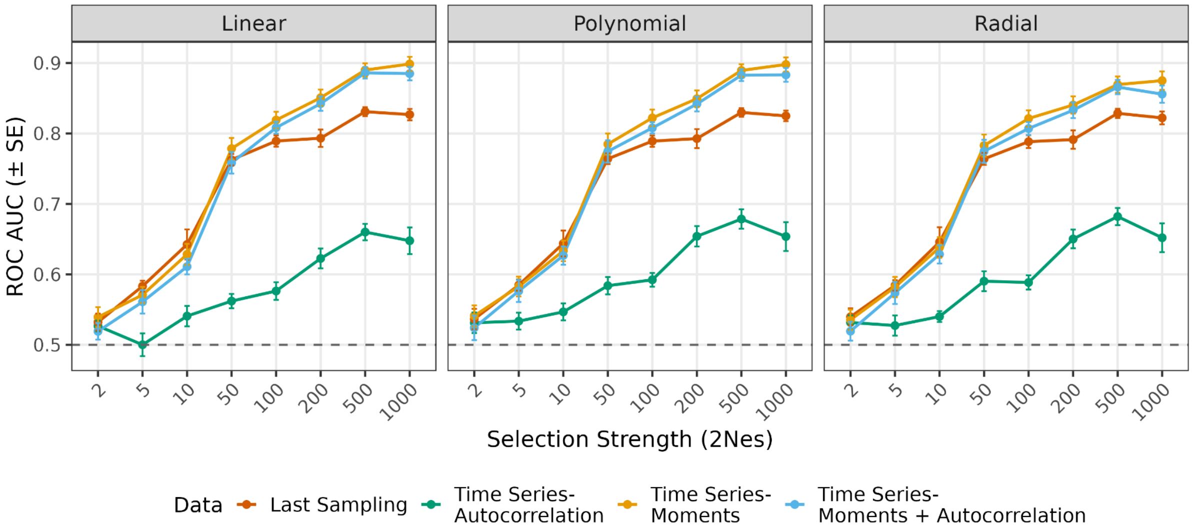


**Supplementary Figure 6.** SVM classification performance, measured by ROC AUC, for distinguishing balancing selection from neutral evolution, conditional on no fixation or loss, as a function of selection strength ($2N_es$). Models were trained using all 23 summary statistics and four different feature sets derived from the same simulations: last sampling, autocorrelation, moments, and moments with autocorrelation. Summary statistics were calculated using $n$=200 haplotypes per sampling point. Points represent mean ROC AUC across the 10 cross-validation folds, with error bars indicating ± SE. Each panel corresponds to SVM analyses with different kernels. The dashed line denotes random expectation (ROC AUC = 0.5).

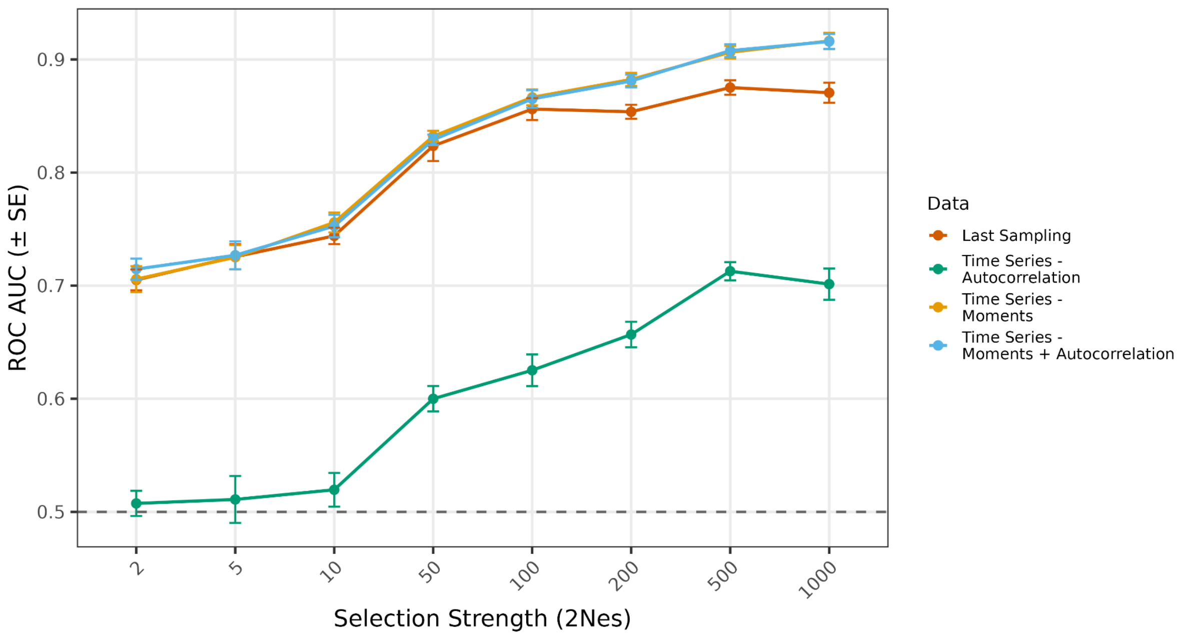


**Supplementary Figure 7.** RF classification performance, measured by ROC AUC, for distinguishing balancing selection from neutral evolution without segregation conditioning as a function of selection strength ($2N_e s$). Models were trained using all 23 summary statistics and four different feature sets derived from the same simulations: last sampling, autocorrelation, moments, and moments with autocorrelation. Summary statistics were calculated using $n$=200 haplotypes per sampling point. Points represent mean ROC AUC across the 10 cross-validation folds, with error bars indicating ± SE. The dashed line denotes random expectation (ROC AUC = 0.5).

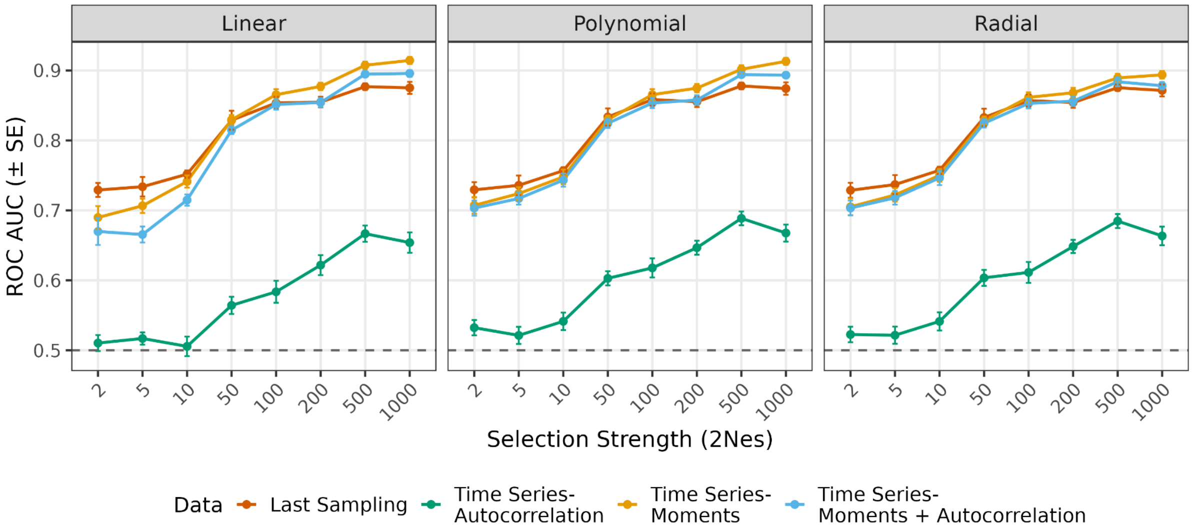


**Supplementary Figure 8.** SVM classification performance, measured by ROC AUC, for distinguishing balancing selection from neutral evolution without segregation conditioning as a function of selection strength ($2N_es$). Models were trained using all 23 summary statistics and four different feature sets derived from the same simulations: last sampling, autocorrelation, moments, and moments with autocorrelation. Summary statistics were calculated using $n$=200 haplotypes per sampling point. Points represent mean ROC AUC across the 10 cross-validation folds, with error bars indicating ± SE. Each panel corresponds to SVM analyses with different kernels. The dashed line denotes random expectation (ROC AUC = 0.5).

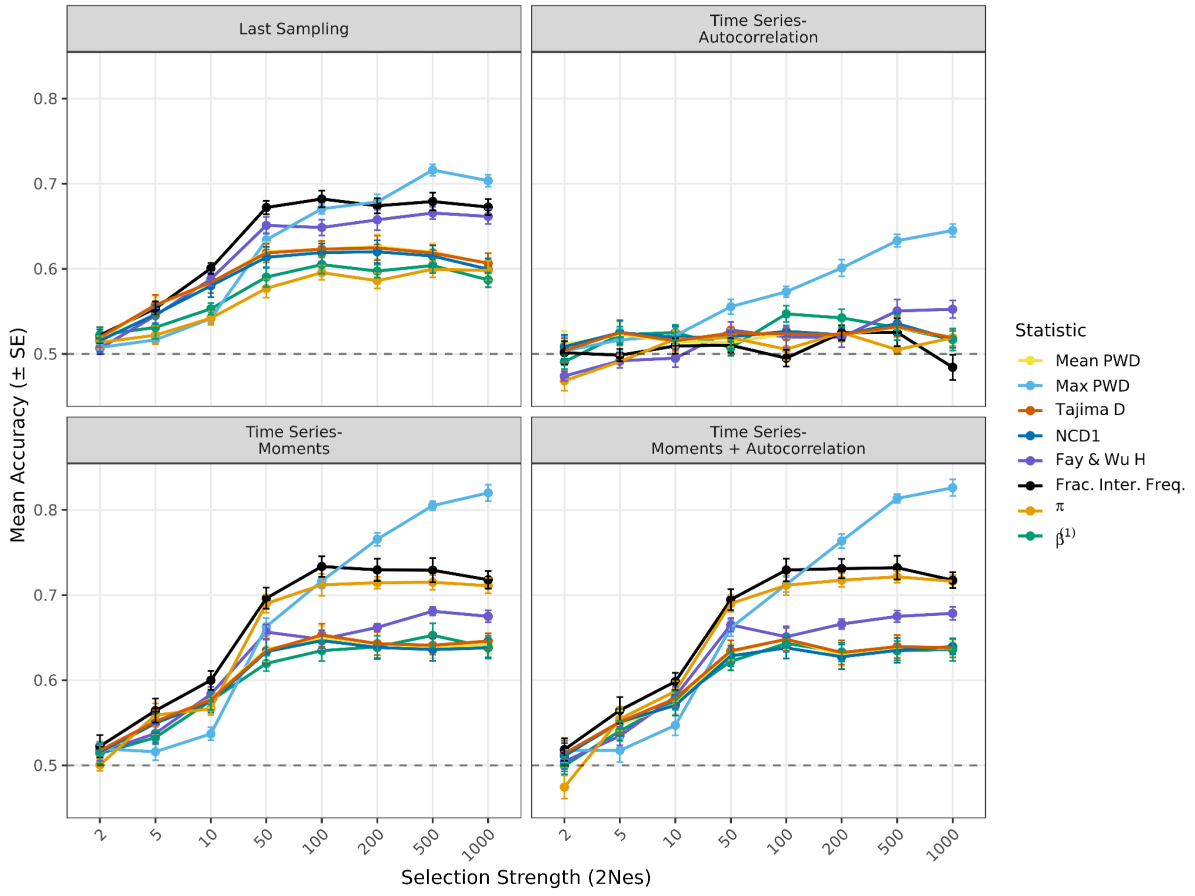


**Supplementary Figure 9.** Linear SVM classification accuracy of individual summary statistics for discriminating balancing selection from neutral evolution, conditional on no fixation or loss, across selection strengths ($2N_es$). Each panel corresponds to a different feature set (last sampling, autocorrelation, moments, and moments with autocorrelation). Summary statistics were calculated using $n$=200 haplotypes per sampling point. Points represent mean classification accuracy across the 10 cross-validation folds, with error bars indicating ± SE. Colored lines indicate individual summary statistics. The dashed line denotes random expectation (accuracy = 0.5).

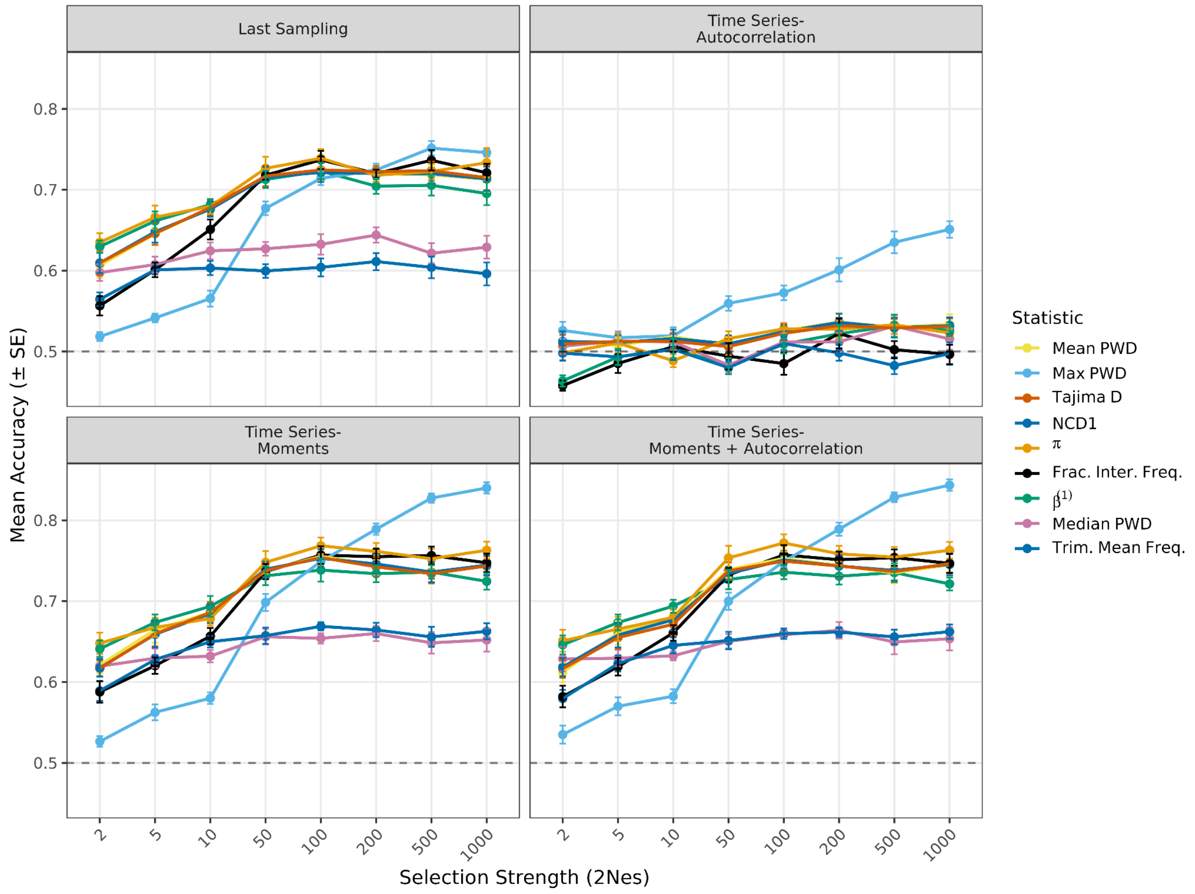


**Supplementary Figure 10.** Linear SVM classification accuracy of individual summary statistics for discriminating balancing selection from neutral evolution without segregation conditioning across selection strengths ($2N_es$). Each panel corresponds to a different feature set (last sampling, autocorrelation, moments, and moments with autocorrelation). Summary statistics were calculated using $n$=200 haplotypes per sampling point. Points represent mean classification accuracy across the 10 cross-validation folds, with error bars indicating ± SE. Colored lines indicate individual summary statistics. The dashed line denotes random expectation (accuracy = 0.5).

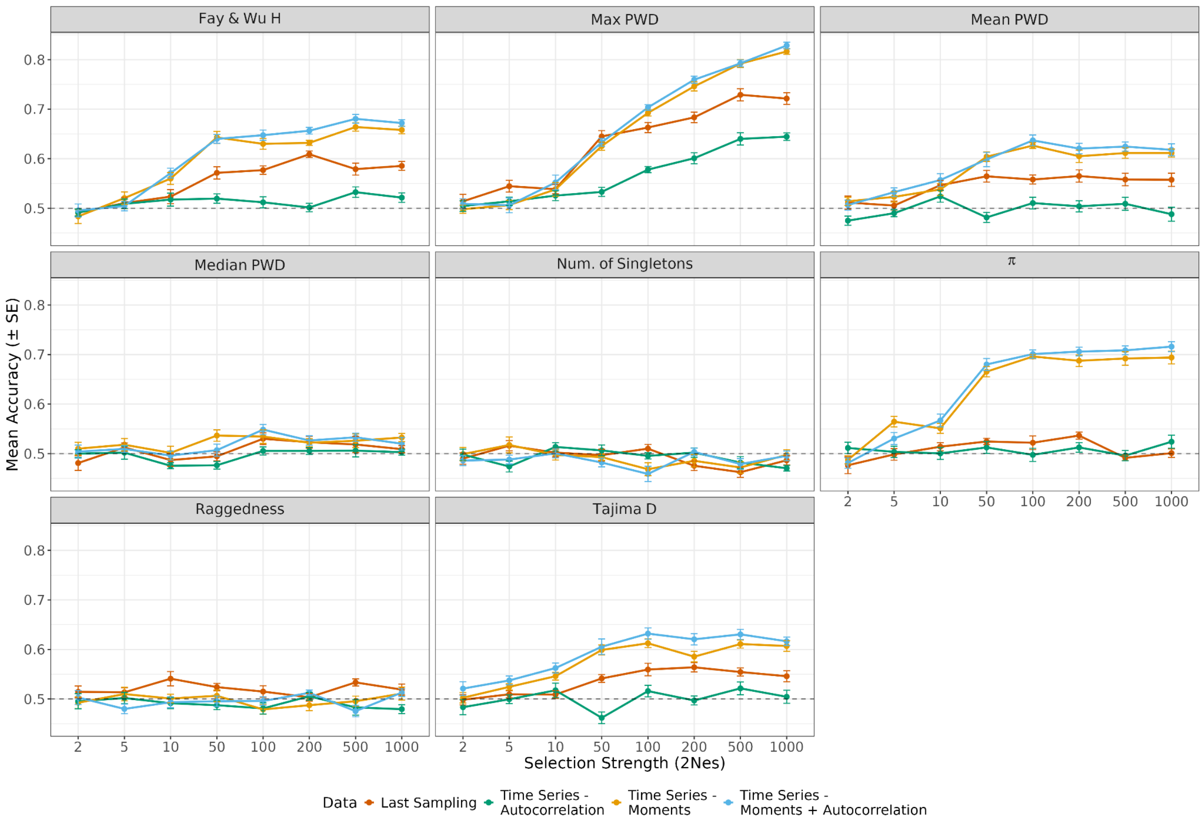


**Supplementary Figure 11.** RF classification accuracy for discriminating balancing selection from neutral evolution, conditional on no fixation or loss, across selection strengths ($2N_es$) using summary statistics derived from the site frequency spectrum (SFS). Each panel shows results for a single SFS-based statistic (panel titles), evaluated using four different feature sets derived from the same simulations: last sampling, autocorrelation, moments, and moments with autocorrelation. Summary statistics were calculated using $n$=200 haplotypes per sampling point. Points represent mean classification accuracy across the 10 cross-validation folds, with error bars indicating ± SE. The dashed line denotes random expectation (accuracy = 0.5).

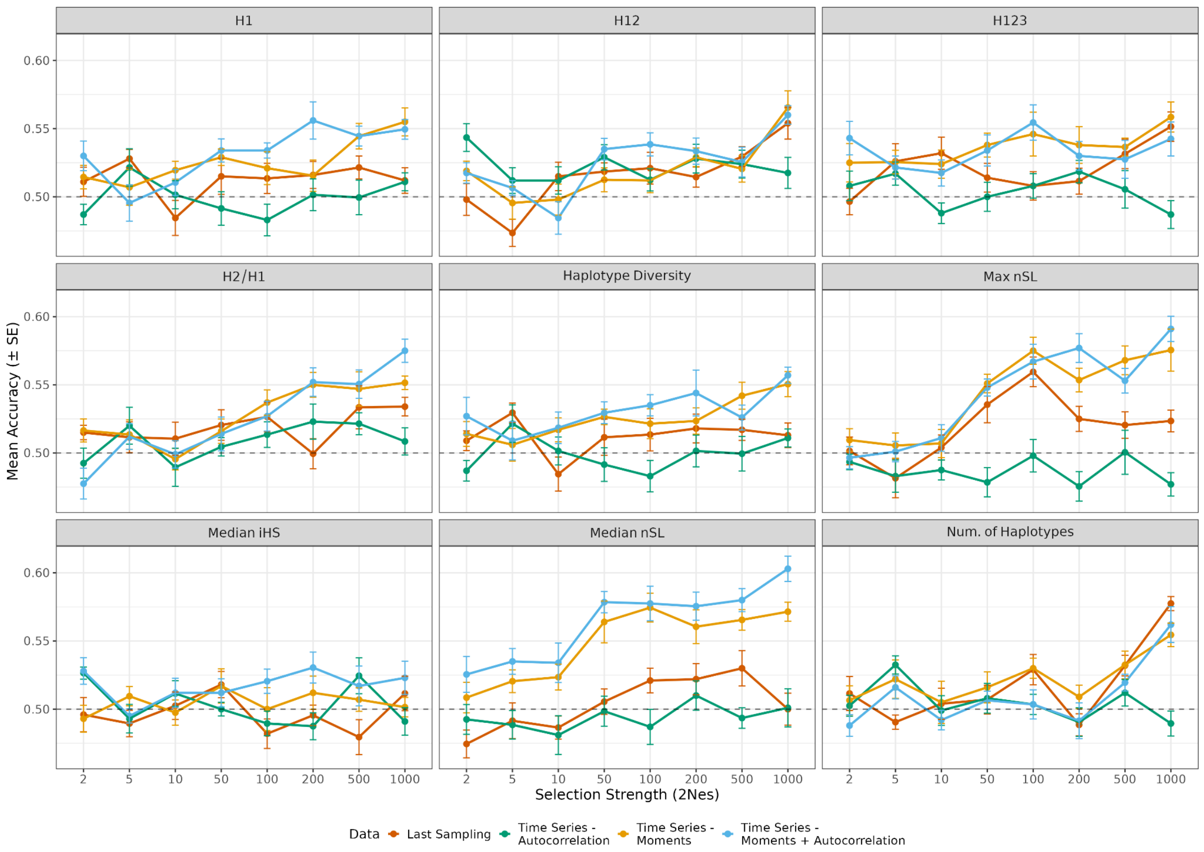


**Supplementary Figure 12.** RF classification accuracy for discriminating balancing selection from neutral evolution, conditional on no fixation or loss, across selection strengths ($2N_es$) using summary statistics based on haplotype-level information. Each panel shows results for a single haplotype-based statistic (panel titles), evaluated using four different feature sets derived from the same simulations: last sampling, autocorrelation, moments, and moments with autocorrelation. Summary statistics were calculated using $n$=200 haplotypes per sampling point. Points represent mean classification accuracy across the 10 cross-validation folds, with error bars indicating ± SE. The dashed line denotes random expectation (accuracy = 0.5).

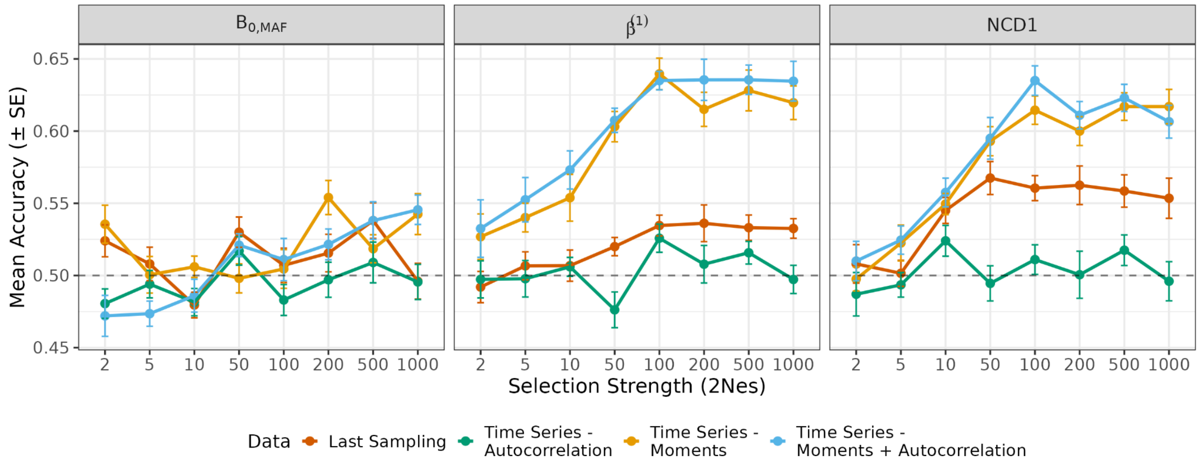


**Supplementary Figure 13.** RF classification accuracy for discriminating balancing selection from neutral evolution, conditional on no fixation or loss, across selection strengths ($2N_es$) using summary statistics specifically designed to detect balancing selection. Each panel shows results for a single balancing selection statistic (panel titles), evaluated using four different feature sets derived from the same simulations: last sampling, autocorrelation, moments, and moments with autocorrelation. Summary statistics were calculated using *n*=200 haplotypes per sampling point. Points represent mean classification accuracy across the 10 cross-validation folds, with error bars indicating ± SE. The dashed line denotes random expectation (accuracy = 0.5).

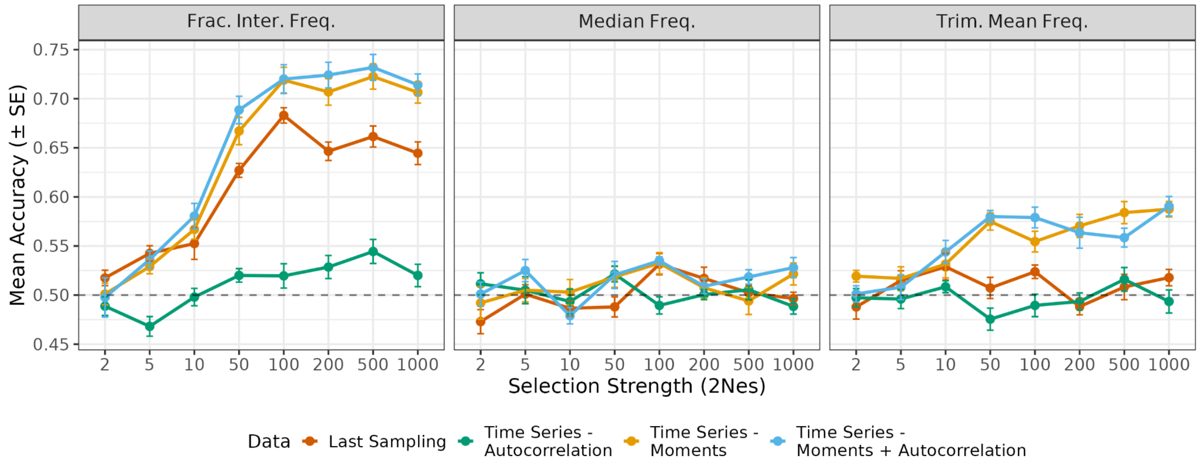


**Supplementary Figure 14.** RF classification accuracy for discriminating balancing selection from neutral evolution, conditional on no fixation or loss, across selection strengths ($2N_e s$) using summary statistics based on frequency information. Each panel shows results for a single frequency-based statistic (panel titles), evaluated using four different feature sets derived from the same simulations: last sampling, autocorrelation, moments, and moments with autocorrelation. Summary statistics were calculated using $n$=200 haplotypes per sampling point. Points represent mean classification accuracy across the 10 cross-validation folds, with error bars indicating ± SE. The dashed line denotes random expectation (accuracy = 0.5).

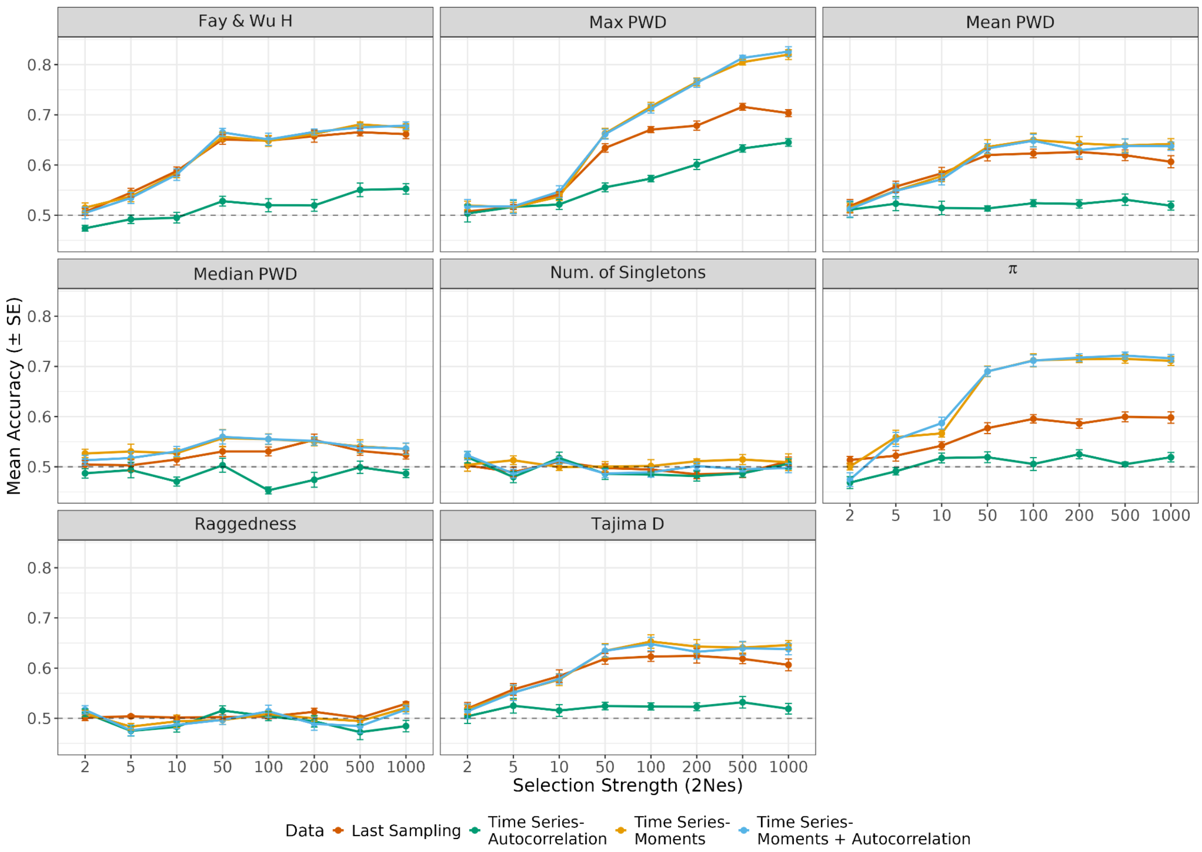


**Supplementary Figure 15.** Linear SVM classification accuracy for discriminating balancing selection from neutral evolution, conditional on no fixation or loss, across selection strengths ($2N_es$) using summary statistics derived from SFS. Each panel shows results for a single SFS-based statistic (panel titles), evaluated using four different feature sets derived from the same simulations: last sampling, autocorrelation, moments, and moments with autocorrelation. Summary statistics were calculated using *n*=200 haplotypes per sampling point. Points represent mean classification accuracy across the 10 cross-validation folds, with error bars indicating ± SE. The dashed line denotes random expectation (accuracy = 0.5).

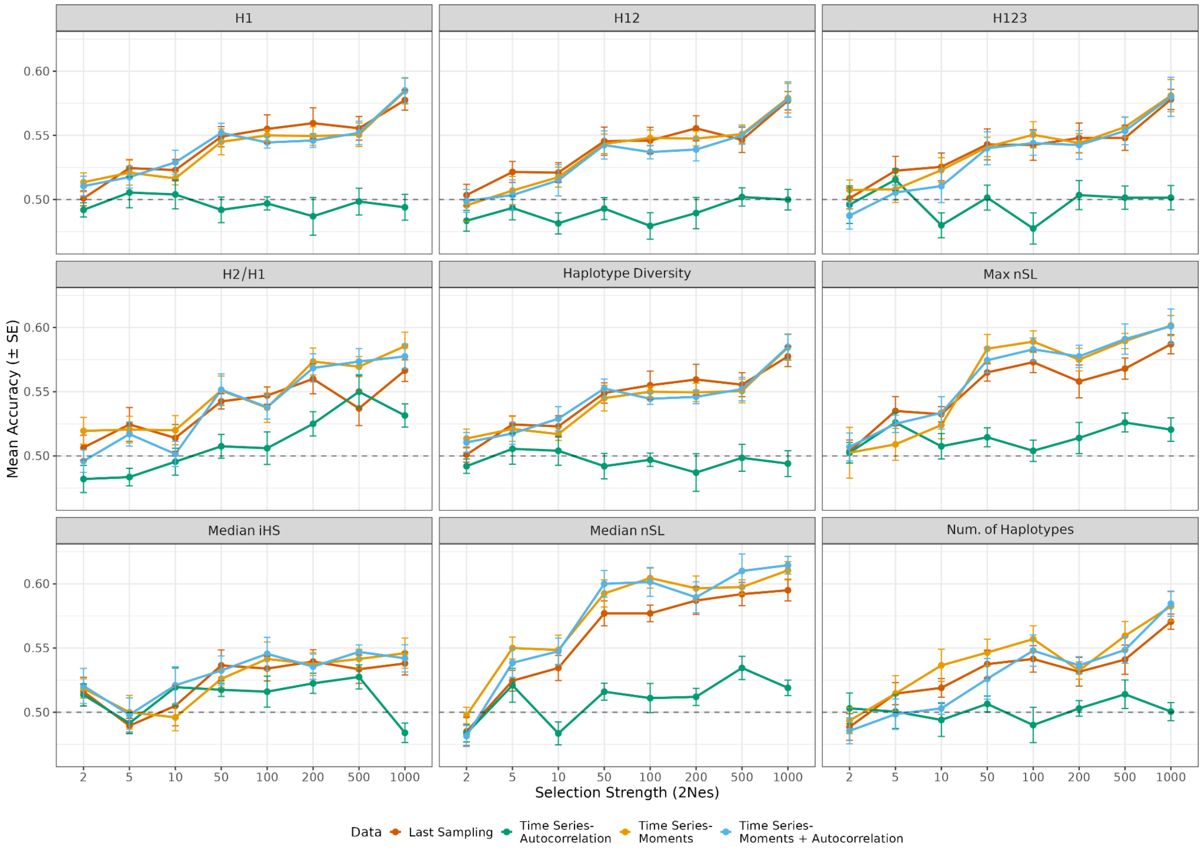


**Supplementary Figure 16.** Linear SVM classification accuracy for discriminating balancing selection from neutral evolution, conditional on no fixation or loss, across selection strengths ($2N_es$) using summary statistics based on haplotype-level information. Each panel shows results for a single haplotype-based statistic (panel titles), evaluated using four different feature sets derived from the same simulations: last sampling, autocorrelation, moments, and moments with autocorrelation. Summary statistics were calculated using $n$=200 haplotypes per sampling point. Points represent mean classification accuracy across the 10 cross-validation folds, with error bars indicating ± SE. The dashed line denotes random expectation (accuracy = 0.5).

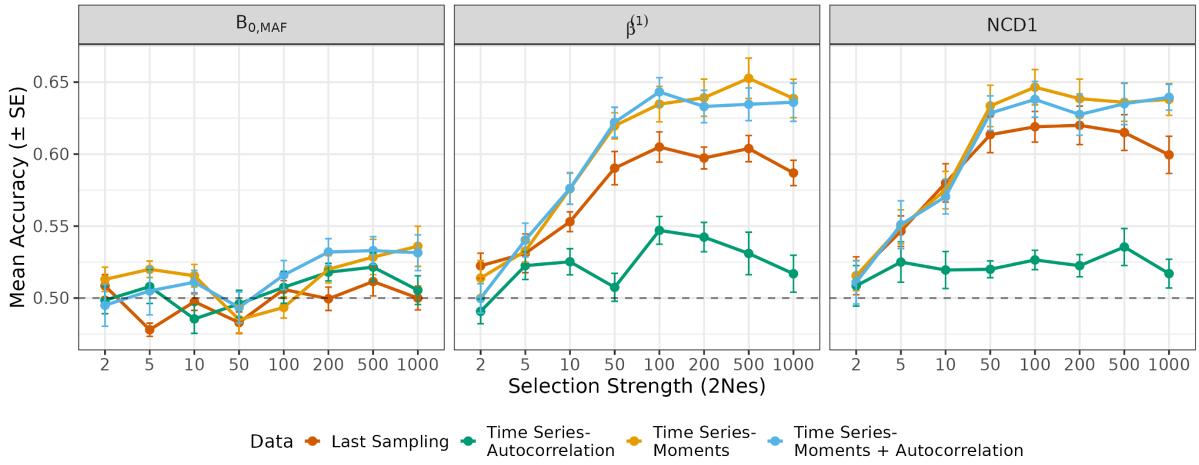


**Supplementary Figure 17.** Linear SVM classification accuracy for discriminating balancing selection from neutral evolution, conditional on no fixation or loss, across selection strengths ($2N_es$) using summary statistics specifically designed to detect balancing selection. Each panel shows results for a single balancing selection statistic (panel titles), evaluated using four different feature sets derived from the same simulations: last sampling, autocorrelation, moments, and moments with autocorrelation. Summary statistics were calculated using *n*=200 haplotypes per sampling point. Points represent mean classification accuracy across the 10 cross-validation folds, with error bars indicating ± SE. The dashed line denotes random expectation (accuracy = 0.5).

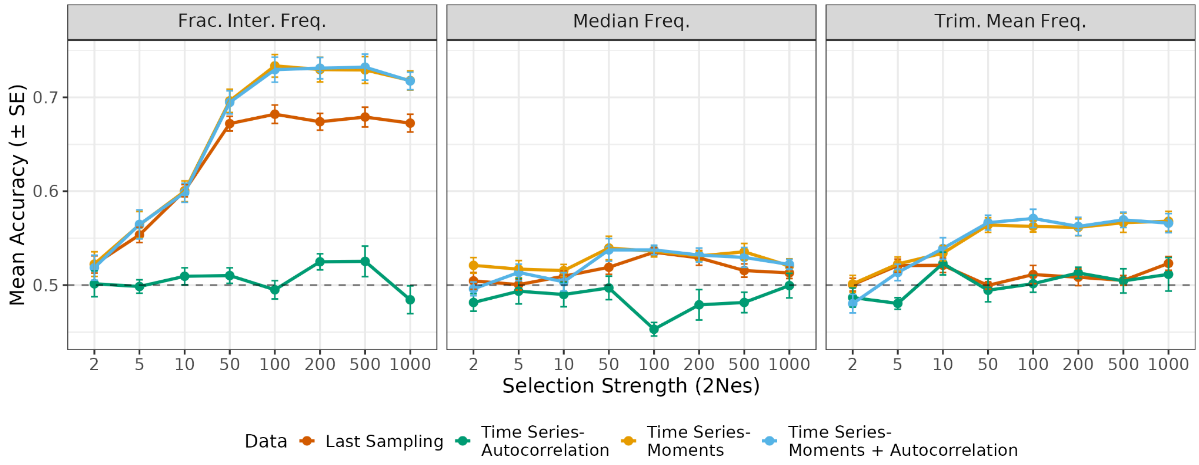


**Supplementary Figure 18.** Linear SVM classification accuracy for discriminating balancing selection from neutral evolution, conditional on no fixation or loss, across selection strengths ($2N_es$) using frequency-based summary statistics. Each panel shows results for a single frequency-based statistic (panel titles), evaluated using four different feature sets derived from the same simulations: last sampling, autocorrelation, moments, and moments with autocorrelation. Summary statistics were calculated using $n$=200 haplotypes per sampling point. Points represent mean classification accuracy across the 10 cross-validation folds, with error bars indicating ± SE. The dashed line denotes random expectation (accuracy = 0.5).

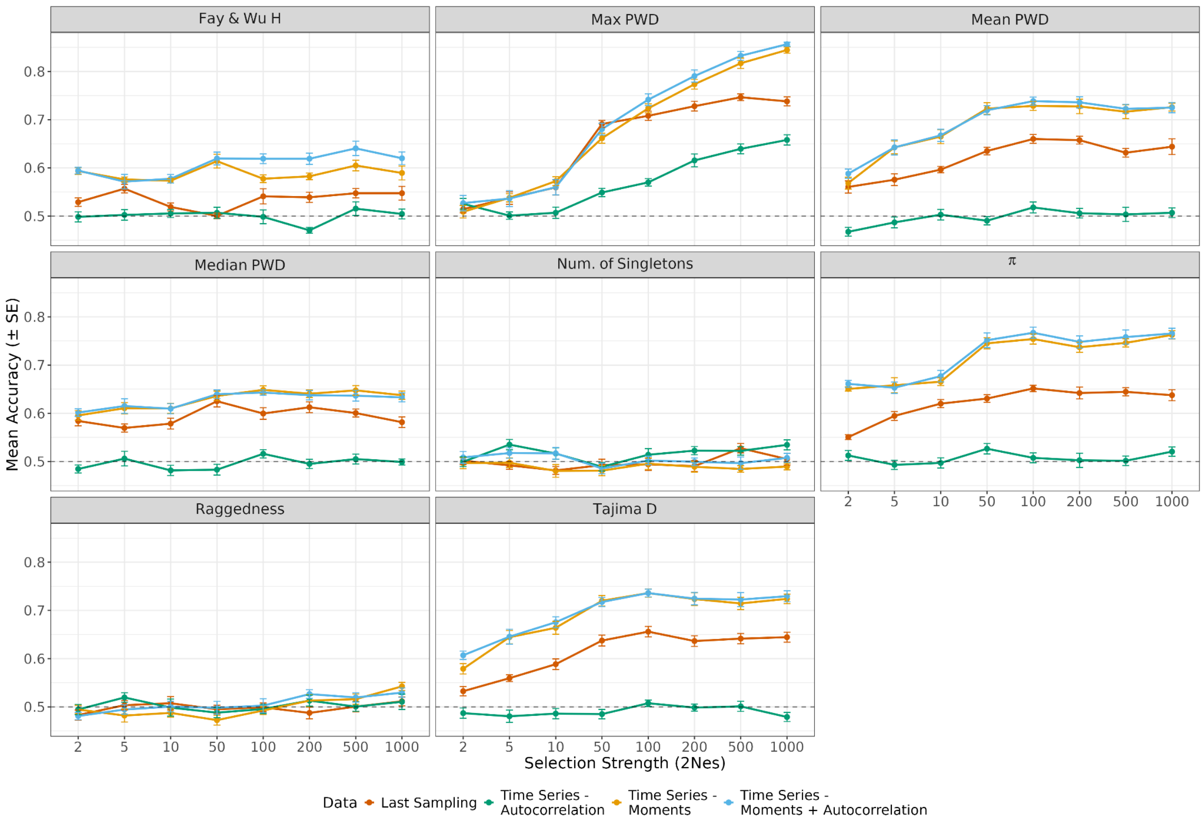


**Supplementary Figure 19.** RF classification accuracy for discriminating balancing selection from neutral evolution without segregation conditioning across selection strengths ($2N_es$) using summary statistics derived from the site frequency spectrum (SFS). Each panel shows results for a single SFS-based statistic (panel titles), evaluated using four different feature sets derived from the same simulations: last sampling, autocorrelation, moments, and moments with autocorrelation. Summary statistics were calculated using $n$=200 haplotypes per sampling point. Points represent mean classification accuracy across the 10 cross-validation folds, with error bars indicating ± SE. The dashed line denotes random expectation (accuracy = 0.5).

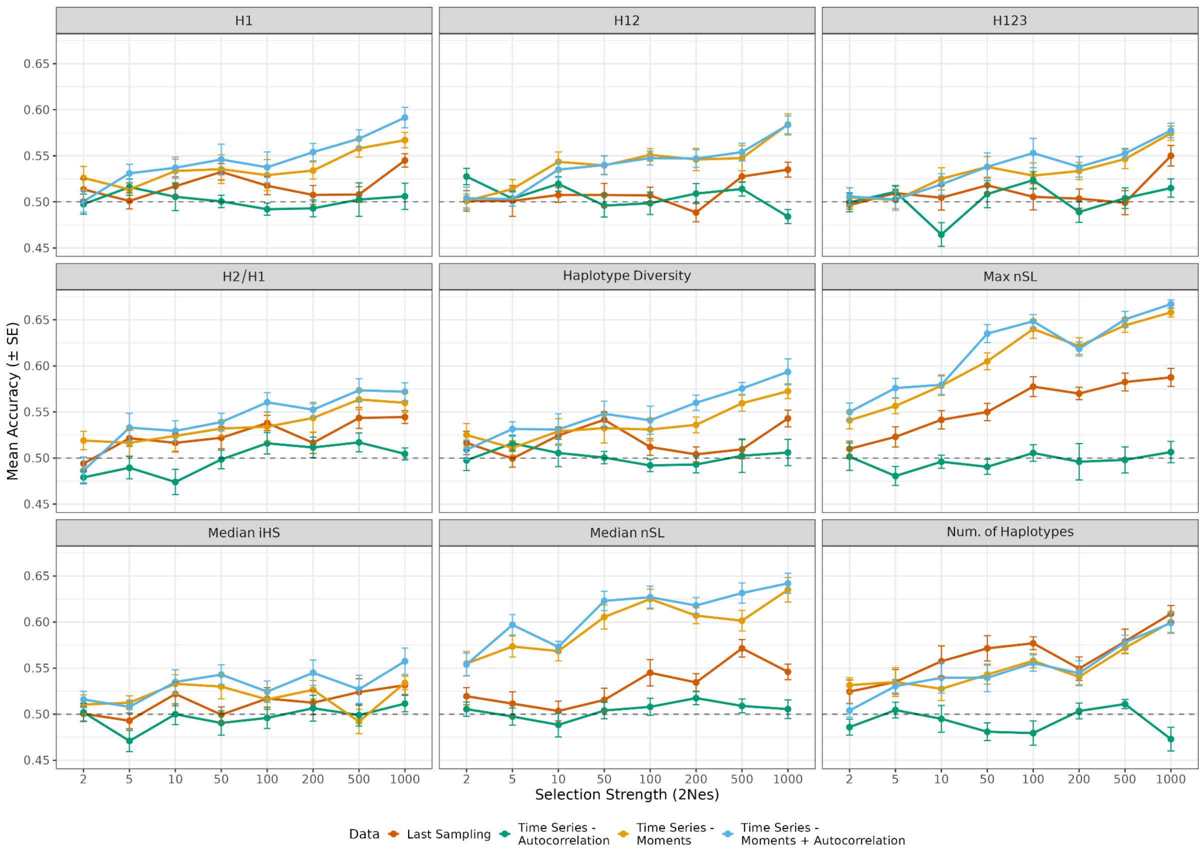


**Supplementary Figure 20.** RF classification accuracy for discriminating balancing selection from neutral evolution without segregation conditioning across selection strengths ($2N_es$) using summary statistics based on haplotype-level information. Each panel shows results for a single haplotype-based statistic (panel titles), evaluated using four different feature sets derived from the same simulations: last sampling, autocorrelation, moments, and moments with autocorrelation. Summary statistics were calculated using $n$=200 haplotypes per sampling point. Points represent mean classification accuracy across the 10 cross-validation folds, with error bars indicating ± SE. The dashed line denotes random expectation (accuracy = 0.5).

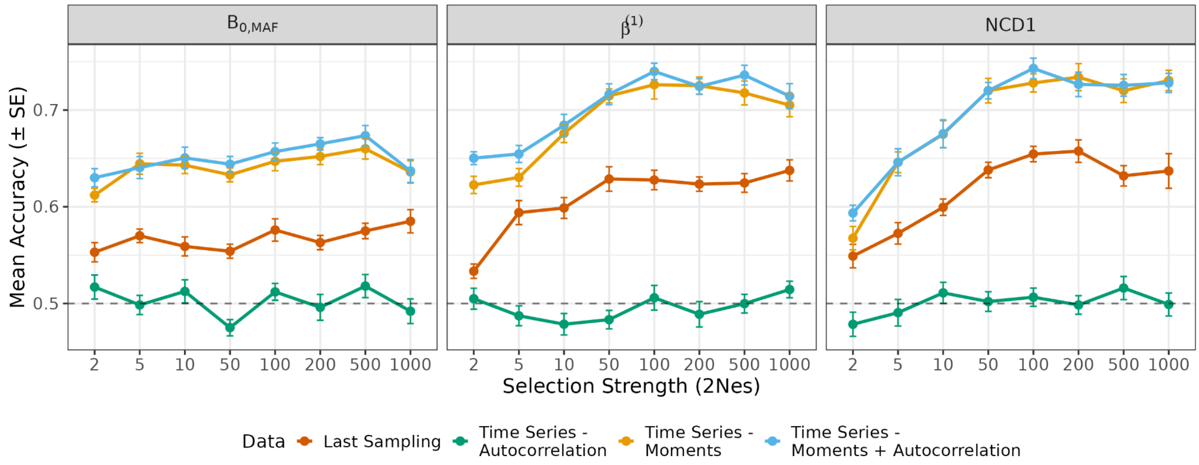


**Supplementary Figure 21.** RF classification accuracy for discriminating balancing selection from neutral evolution without segregation conditioning across selection strengths ($2N_es$) using summary statistics specifically designed to detect balancing selection. Each panel shows results for a single balancing selection statistic (panel titles), evaluated using four different feature sets derived from the same simulations: last sampling, autocorrelation, moments, and moments with autocorrelation. Summary statistics were calculated using *n*=200 haplotypes per sampling point. Points represent mean classification accuracy across the 10 cross-validation folds, with error bars indicating ± SE. The dashed line denotes random expectation (accuracy = 0.5).

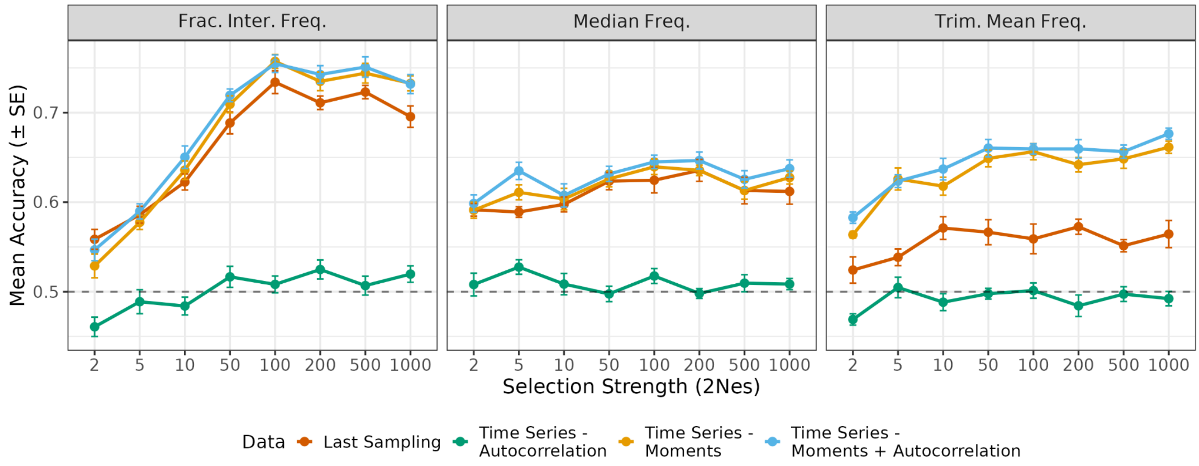


**Supplementary Figure 22.** RF classification accuracy for discriminating balancing selection from neutral evolution without segregation conditioning across selection strengths ($2N_es$) using summary statistics based on frequency information. Each panel shows results for a single frequency-based statistic (panel titles), evaluated using four different feature sets derived from the same simulations: last sampling, autocorrelation, moments, and moments with autocorrelation. Summary statistics were calculated using *n*=200 haplotypes per sampling point. Points represent mean classification accuracy across the 10 cross-validation folds, with error bars indicating ± SE. The dashed line denotes random expectation (accuracy = 0.5).

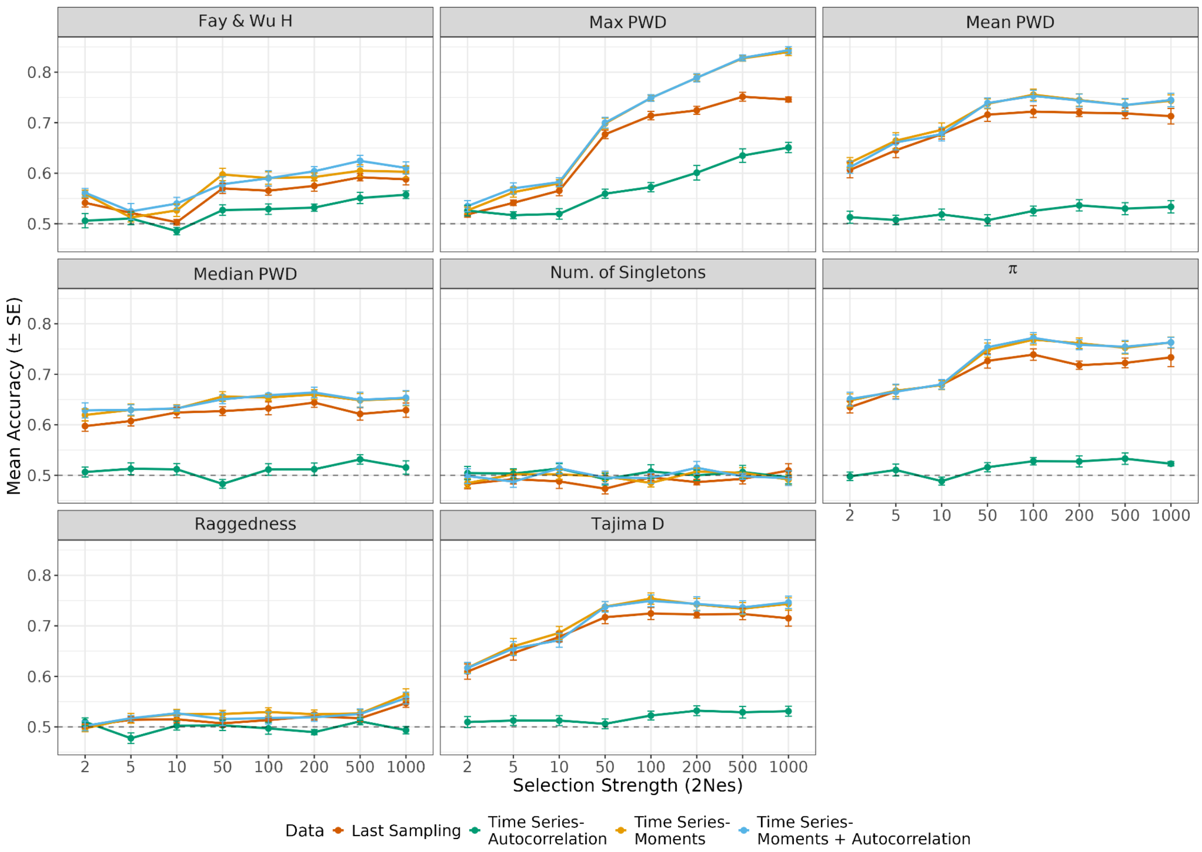


**Supplementary Figure 23.** Linear SVM classification accuracy for discriminating balancing selection from neutral evolution without segregation conditioning across selection strengths ($2N_es$) using summary statistics derived from SFS. Each panel shows results for a single SFS-based statistic (panel titles), evaluated using four different feature sets derived from the same simulations: last sampling, autocorrelation, moments, and moments with autocorrelation. Summary statistics were calculated using $n$=200 haplotypes per sampling point. Points represent mean classification accuracy across the 10 cross-validation folds, with error bars indicating ± SE. The dashed line denotes random expectation (accuracy = 0.5).

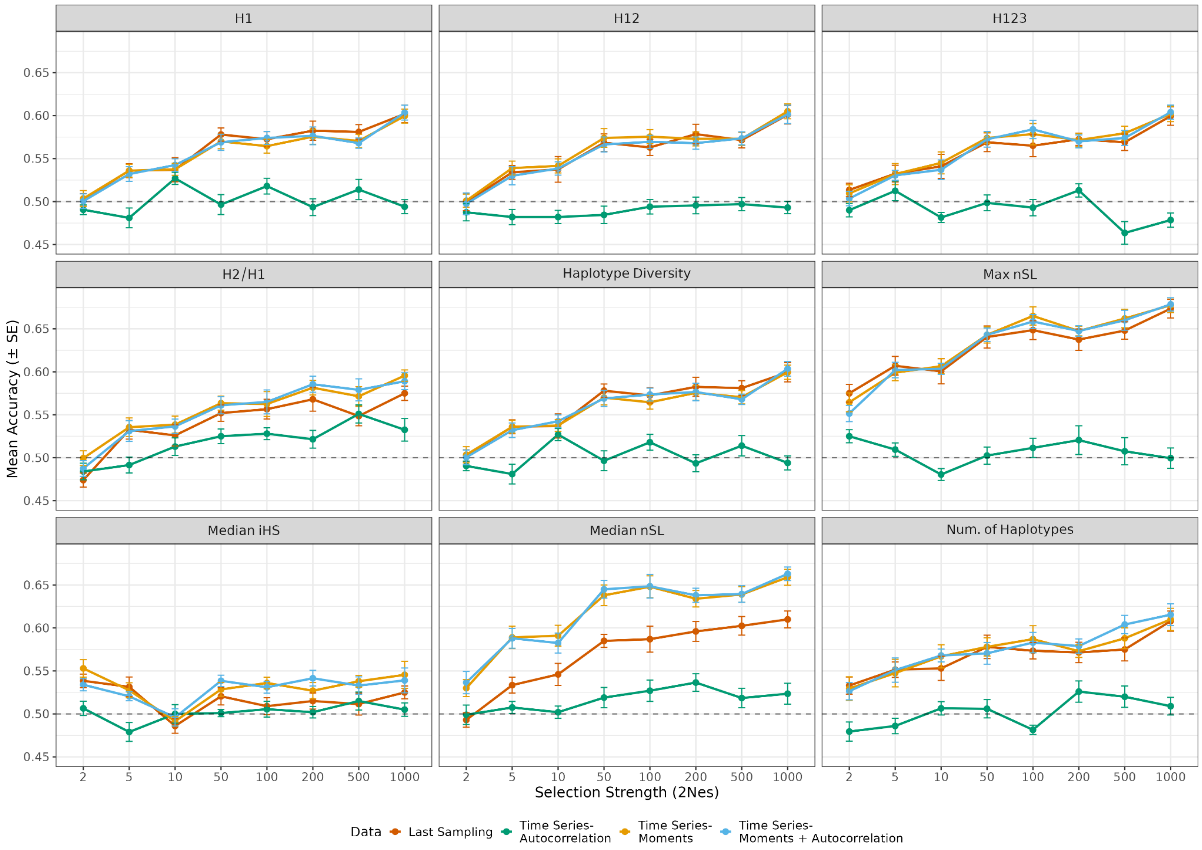


**Supplementary Figure 24.** Linear SVM classification accuracy for discriminating balancing selection from neutral evolution without segregation conditioning across selection strengths ($2N_es$) using summary statistics based on haplotype-level information. Each panel shows results for a single haplotype-based statistic (panel titles), evaluated using four different feature sets derived from the same simulations: last sampling, autocorrelation, moments, and moments with autocorrelation. Summary statistics were calculated using $n$=200 haplotypes per sampling point. Points represent mean classification accuracy across the 10 cross-validation folds, with error bars indicating ± SE. The dashed line denotes random expectation (accuracy = 0.5).

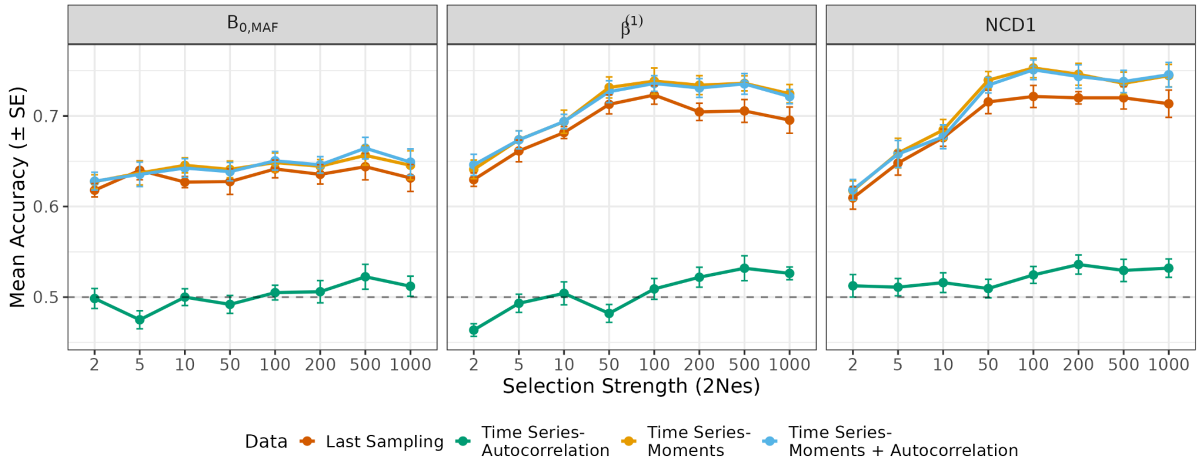


**Supplementary Figure 25.** Linear SVM classification accuracy for discriminating balancing selection from neutral evolution without segregation conditioning across selection strengths ($2N_es$) using summary statistics specifically designed to detect balancing selection. Each panel shows results for a single balancing selection statistic (panel titles), evaluated using four different feature sets derived from the same simulations: last sampling, autocorrelation, moments, and moments with autocorrelation. Summary statistics were calculated using $n$=200 haplotypes per sampling point. Points represent mean classification accuracy across the 10 cross-validation folds, with error bars indicating ± SE. The dashed line denotes random expectation (accuracy = 0.5).

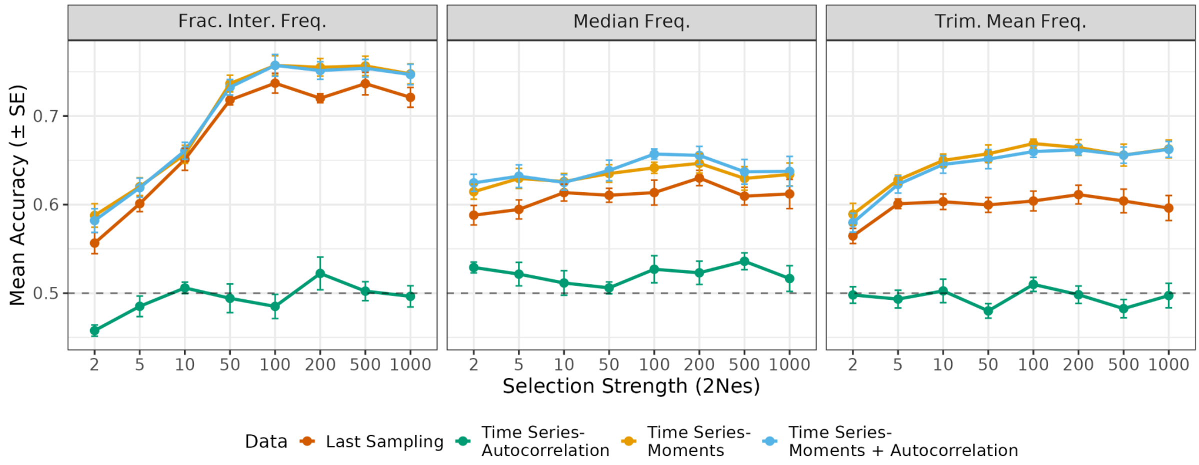


**Supplementary Figure 26.** Linear SVM classification accuracy for discriminating balancing selection from neutral evolution without segregation conditioning across selection strengths ($2N_es$) using frequency-based summary statistics. Each panel shows results for a single frequency-based statistic (panel titles), evaluated using four different feature sets derived from the same simulations: last sampling, autocorrelation, moments, and moments with autocorrelation. Summary statistics were calculated using $n$=200 haplotypes per sampling point. Points represent mean classification accuracy across the 10 cross-validation folds, with error bars indicating ± SE. The dashed line denotes random expectation (accuracy = 0.5).

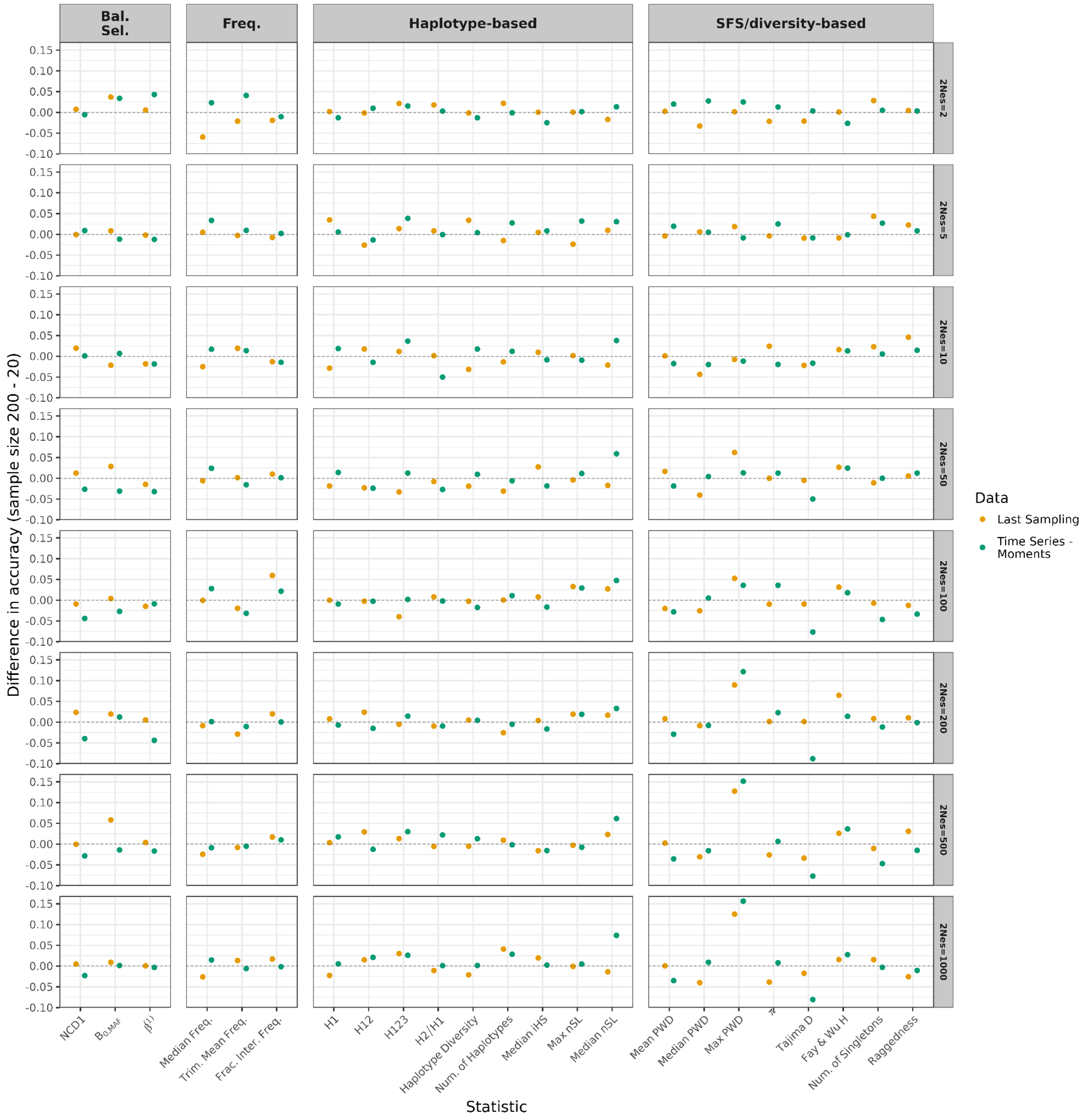


**Supplementary Figure 27.** Differences in mean RF classification accuracy for discriminating balancing selection from neutral evolution, conditional on no fixation or loss, using individual summary statistics estimated for original (*n*=200 haplotypes) vs. downsampled data (*n*=20 haplotypes). Horizontal panels show selection strengths used to generate balancing selection simulations. Vertical panels show the type of genetic data each statistic utilizes. Statistics were evaluated using two different feature sets derived from the same simulations: last sampling and moments. Trim. Mean Freq.: Trimmed mean frequency (0.1 < frequency < 0.9); Frac. Inter. Freq.: Fraction of intermediate frequency SNPs; Median freq.: Median frequency. For other names of statistics, see Table 1.

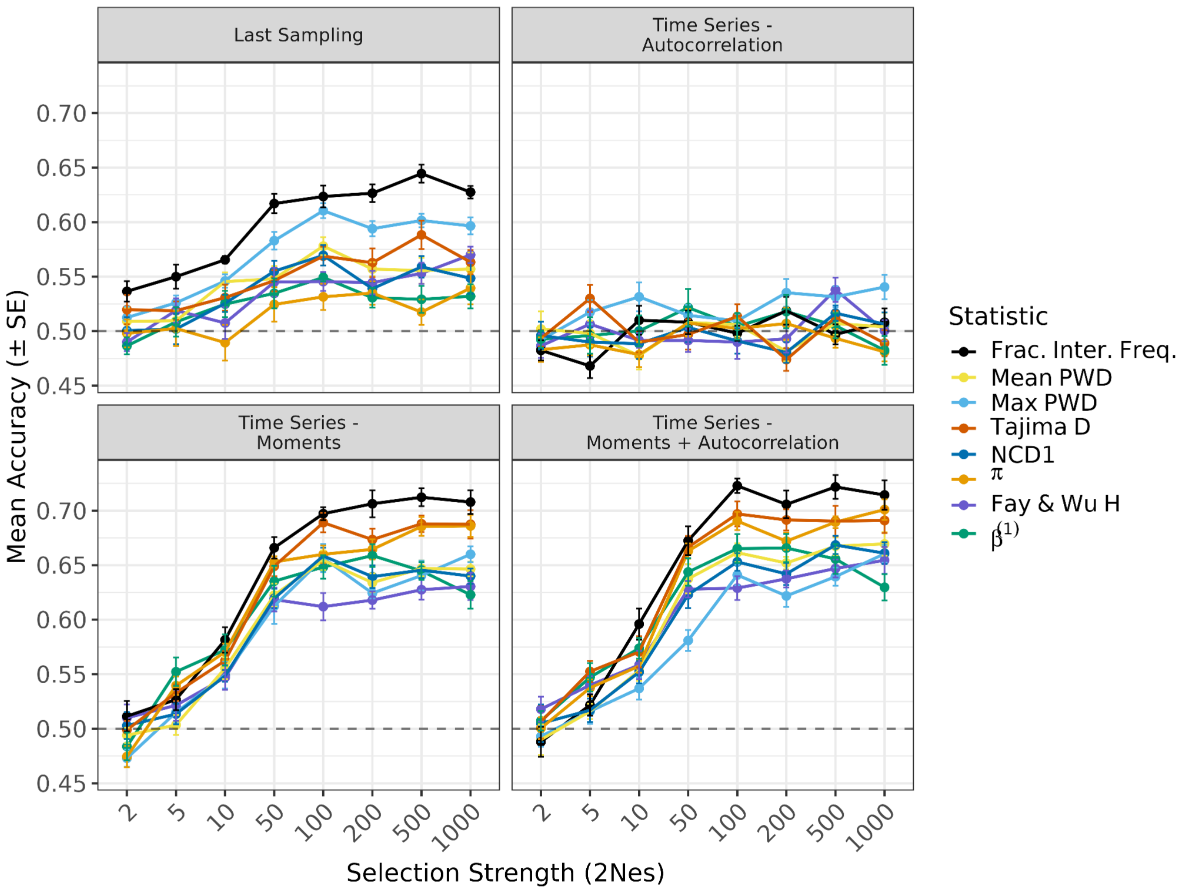


**Supplementary Figure 28.** RF classification accuracy of individual summary statistics estimated for downsampled data ($n$=20 haplotypes) for discriminating balancing selection from neutral evolution, conditional on no fixation or loss, across selection strengths ($2N_es$). Each panel corresponds to a different feature set (last sampling, autocorrelation, moments, and moments with autocorrelation). Points represent mean classification accuracy across the 10 cross-validation folds, with error bars indicating ± SE. Colored lines indicate individual summary statistics. The dashed line denotes random expectation (accuracy = 0.5). Frac. Inter. Freq.: Fraction of intermediate frequency SNPs; Max PWD: Maximum mean-pairwise-differences; Mean PWD: Mean of mean-pairwise-differences.

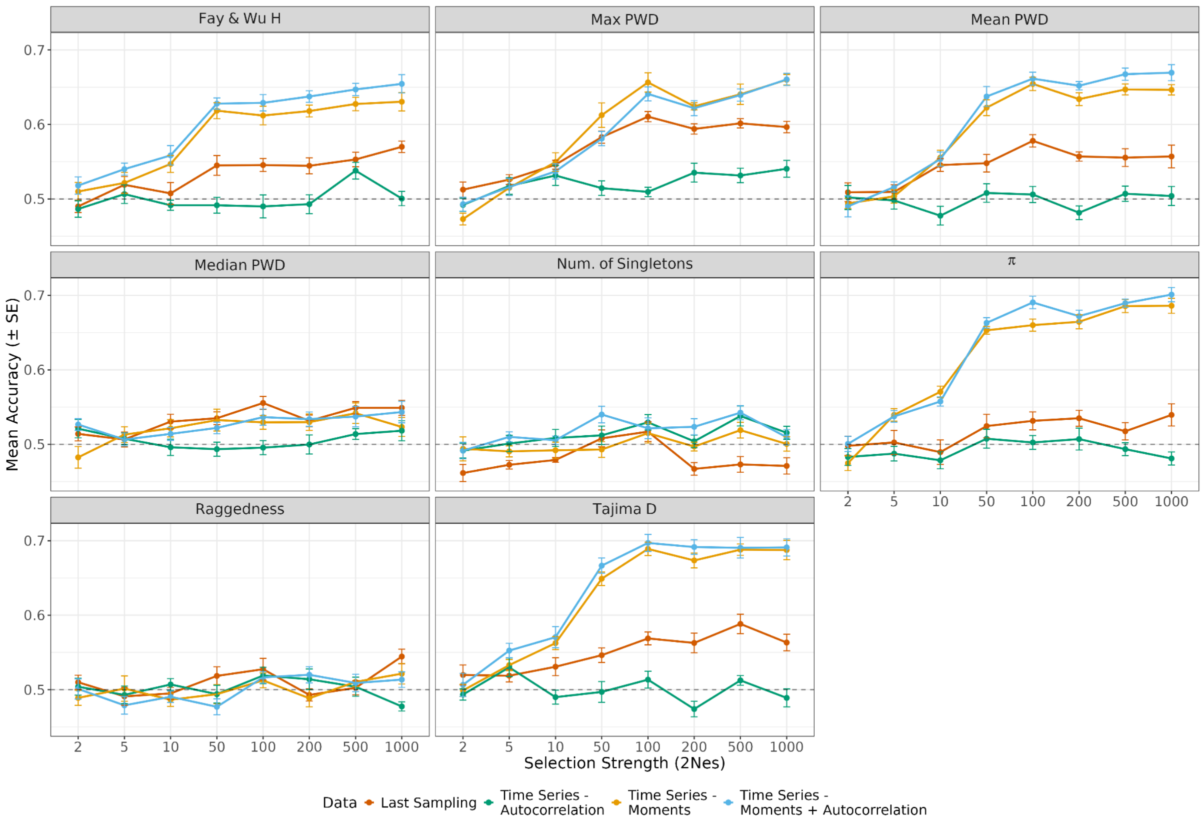


**Supplementary Figure 29.** RF classification accuracy for discriminating balancing selection from neutral evolution, conditional on no fixation or loss, across selection strengths ($2N_es$) using summary statistics derived from the site frequency spectrum (SFS) on downsampled data ($n$=20 haplotypes). Each panel shows results for a single SFS-based statistic (panel titles), evaluated using four different feature sets derived from the same simulations: last sampling, autocorrelation, moments, and moments with autocorrelation. Points represent mean classification accuracy across the 10 cross-validation folds, with error bars indicating ± SE. The dashed line denotes random expectation (accuracy = 0.5).

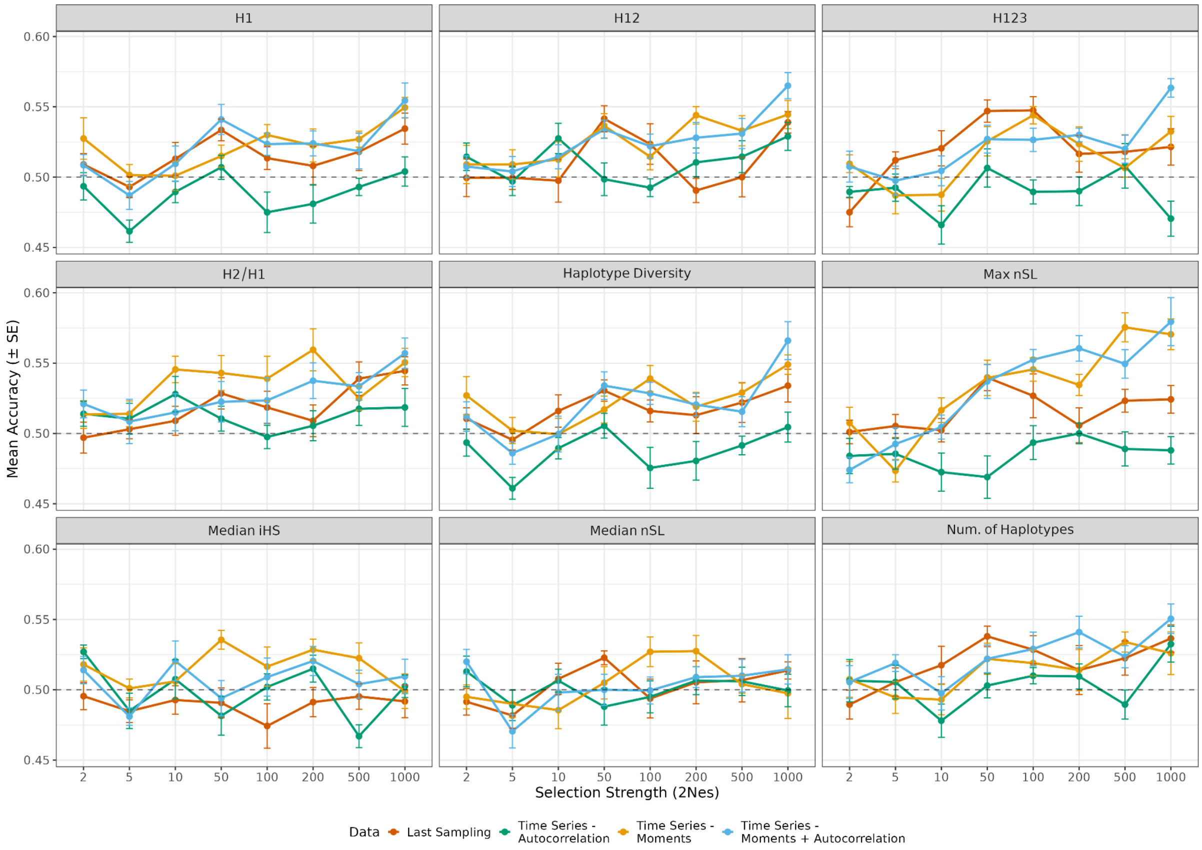


**Supplementary Figure 30.** RF classification accuracy for discriminating balancing selection from neutral evolution, conditional on no fixation or loss, across selection strengths ($2N_es$) using summary statistics based on haplotype-level information and estimated for downsampled data ($n$=20 haplotypes). Each panel shows results for a single haplotype-based statistic (panel titles), evaluated using four different feature sets derived from the same simulations: last sampling, autocorrelation, moments, and moments with autocorrelation. Points represent mean classification accuracy across the 10 cross-validation folds, with error bars indicating ± SE. The dashed line denotes random expectation (accuracy = 0.5).

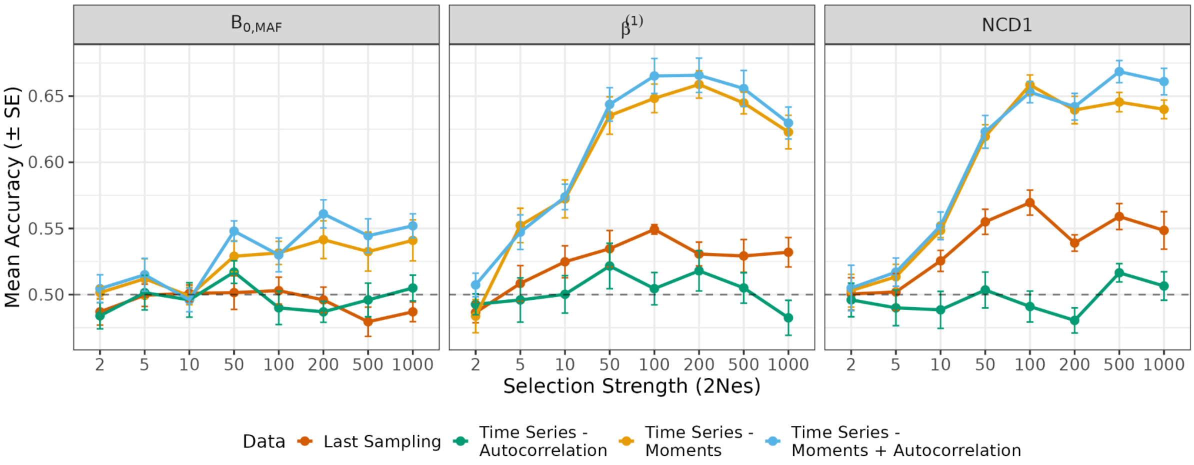


**Supplementary Figure 31.** RF classification accuracy for discriminating balancing selection from neutral evolution, conditional on no fixation or loss, across selection strengths ($2N_es$) using summary statistics specifically designed to detect balancing selection and estimated for downsampled data (*n*=20 haplotypes). Each panel shows results for a single balancing selection statistic (panel titles), evaluated using four different feature sets derived from the same simulations: last sampling, autocorrelation, moments, and moments with autocorrelation. Points represent mean classification accuracy across the 10 cross-validation folds, with error bars indicating ± SE. The dashed line denotes random expectation (accuracy = 0.5).

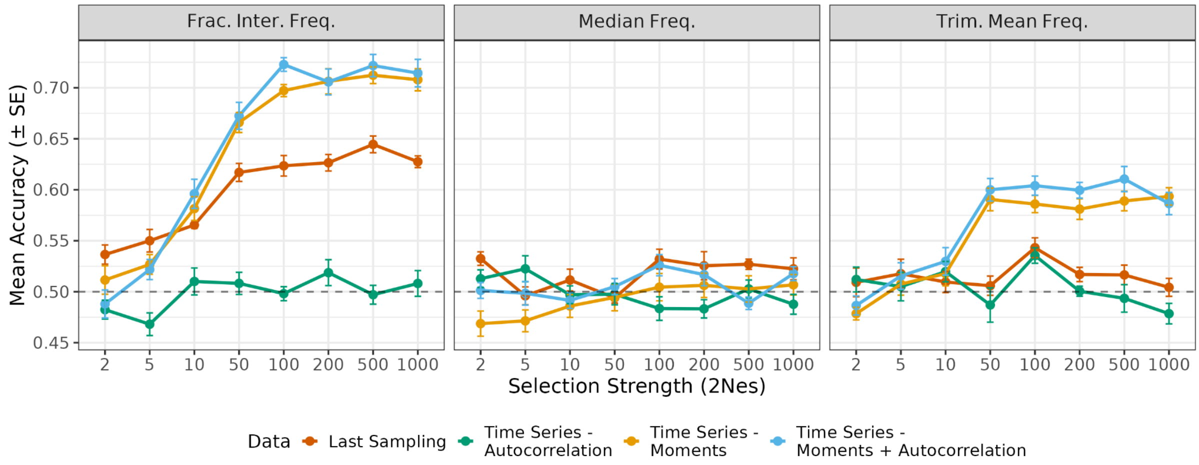


**Supplementary Figure 32.** RF classification accuracy for discriminating balancing selection from neutral evolution, conditional on no fixation or loss, across selection strengths ($2N_e s$) using summary statistics based on frequency information and estimated for downsampled data ($n$=20 haplotypes). Each panel shows results for a single frequency-based statistic (panel titles), evaluated using four different feature sets derived from the same simulations: last sampling, autocorrelation, moments, and moments with autocorrelation. Points represent mean classification accuracy across the 10 cross-validation folds, with error bars indicating ± SE. The dashed line denotes random expectation (accuracy = 0.5).

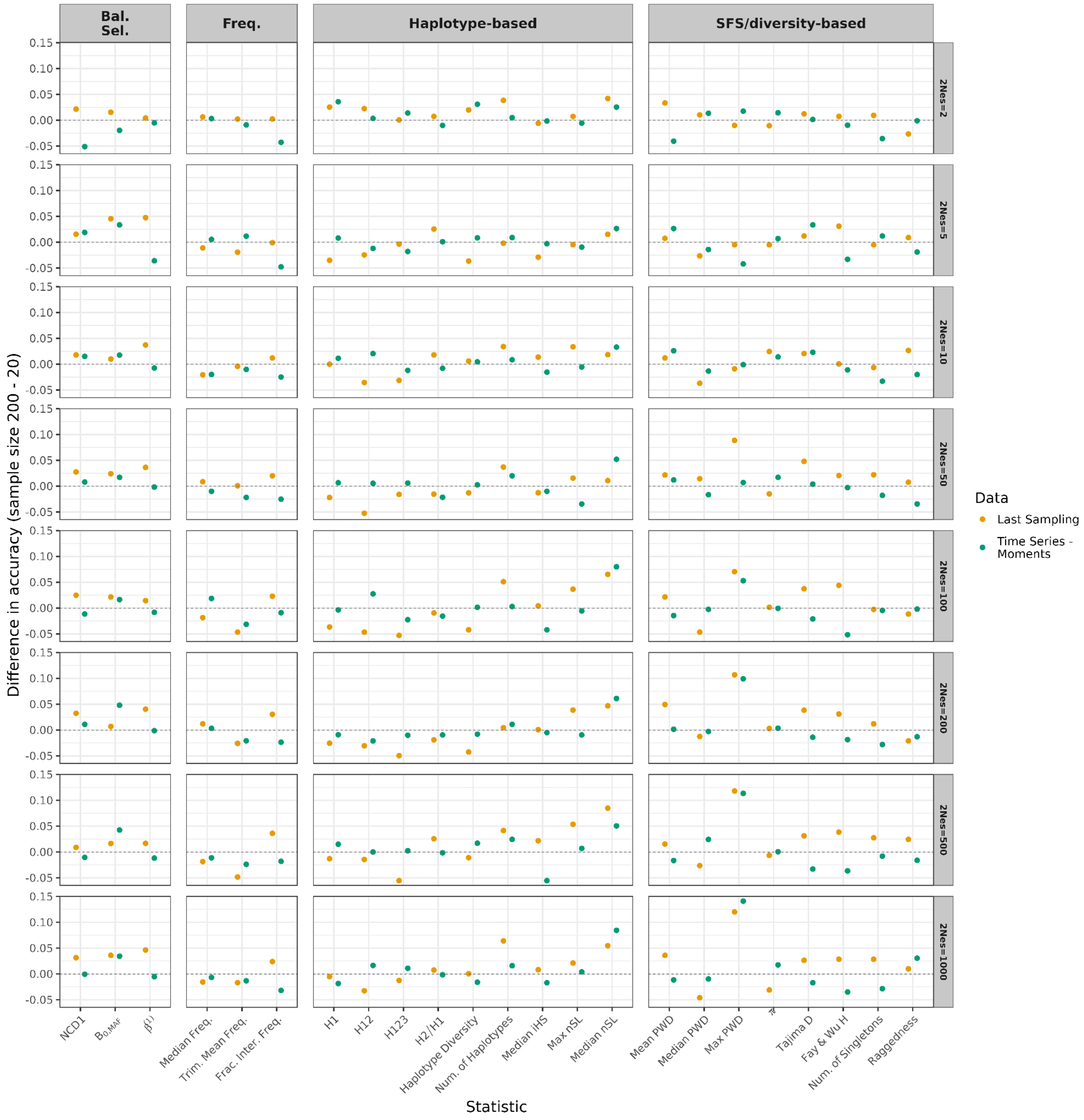


**Supplementary Figure 33.** Differences in mean RF classification accuracy for discriminating balancing selection from neutral evolution without segregation conditioning using individual summary statistics estimated for original (*n*=200 haplotypes) vs. downsampled data (*n*=20 haplotypes). Horizontal panels show selection strengths used to generate balancing selection simulations. Vertical panels show the type of genetic data each statistic utilizes. Statistics were evaluated using two different feature sets derived from the same simulations: last sampling and moments. Trim. Mean Freq.: Trimmed mean frequency (0.1 < frequency < 0.9); Frac. Inter. Freq.: Fraction of intermediate frequency SNPs; Median freq.: Median frequency. For other names of statistics, see Table 1.

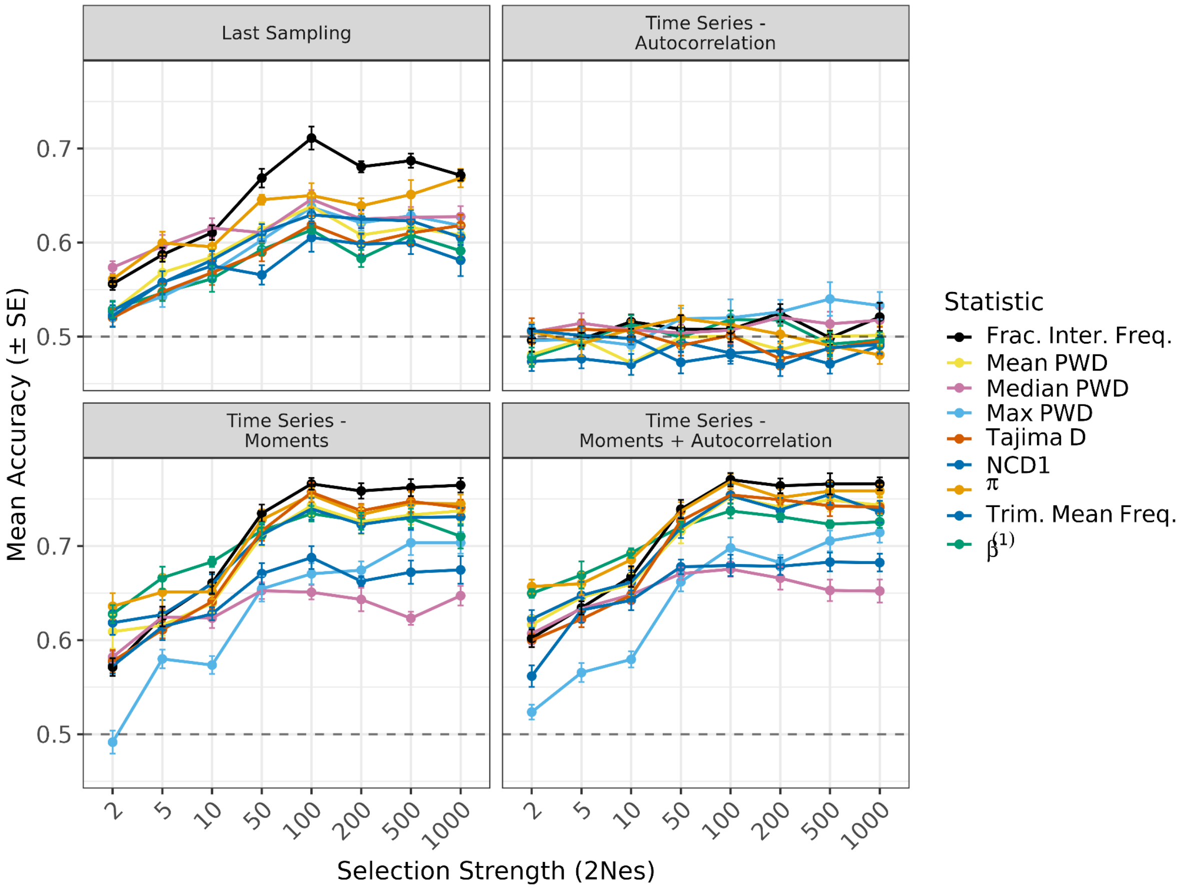


**Supplementary Figure 34.** RF classification accuracy of individual summary statistics estimated for downsampled data (*n*=20 haplotypes) for discriminating balancing selection from neutral evolution without segregation conditioning across selection strengths ($2N_e s$). Each panel corresponds to a different feature set (last sampling, autocorrelation, moments, and moments with autocorrelation). Points represent mean classification accuracy across the 10 cross-validation folds, with error bars indicating ± SE. Colored lines indicate individual summary statistics. The dashed line denotes random expectation (accuracy = 0.5). Frac. Inter. Freq.: Fraction of intermediate frequency SNPs; Mean, Median $H_o$: Mean and median heterozygosity, respectively; Median $H_o/H_e$: Median observed over expected heterozygosity; Max PWD, Mean PWD, Median PWD: Maximum, mean, and median per-site mean-pairwise-differences, respectively. Trim. Mean Freq.: Trimmed mean frequency (0.1 < frequency < 0.9).

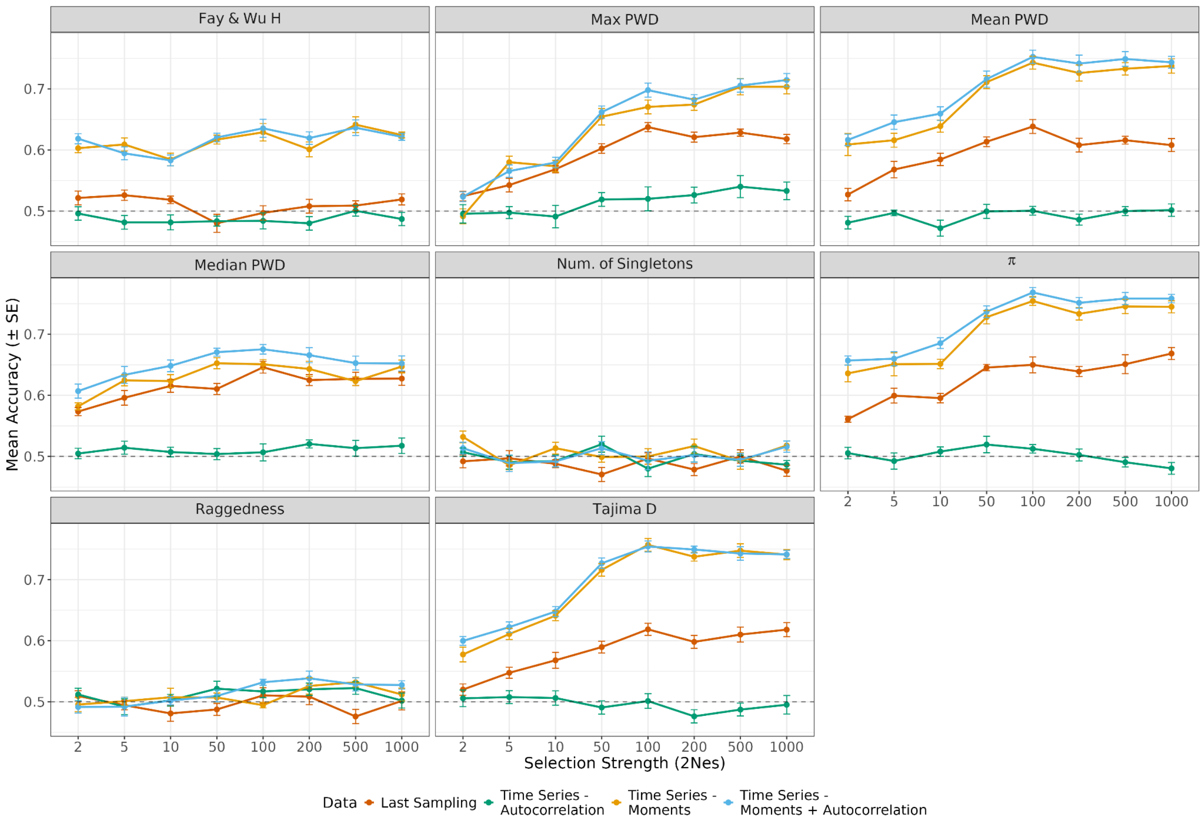


**Supplementary Figure 35.** RF classification accuracy for discriminating balancing selection from neutral evolution without segregation conditioning across selection strengths ($2N_es$) using summary statistics derived from the site frequency spectrum (SFS) on downsampled data ($n$=20 haplotypes per sampling point). Each panel shows results for a single SFS-based statistic (panel titles), evaluated using four different feature sets derived from the same simulations: last sampling, autocorrelation, moments, and moments with autocorrelation. Points represent mean classification accuracy across the 10 cross-validation folds, with error bars indicating ± SE. The dashed line denotes random expectation (accuracy = 0.5).

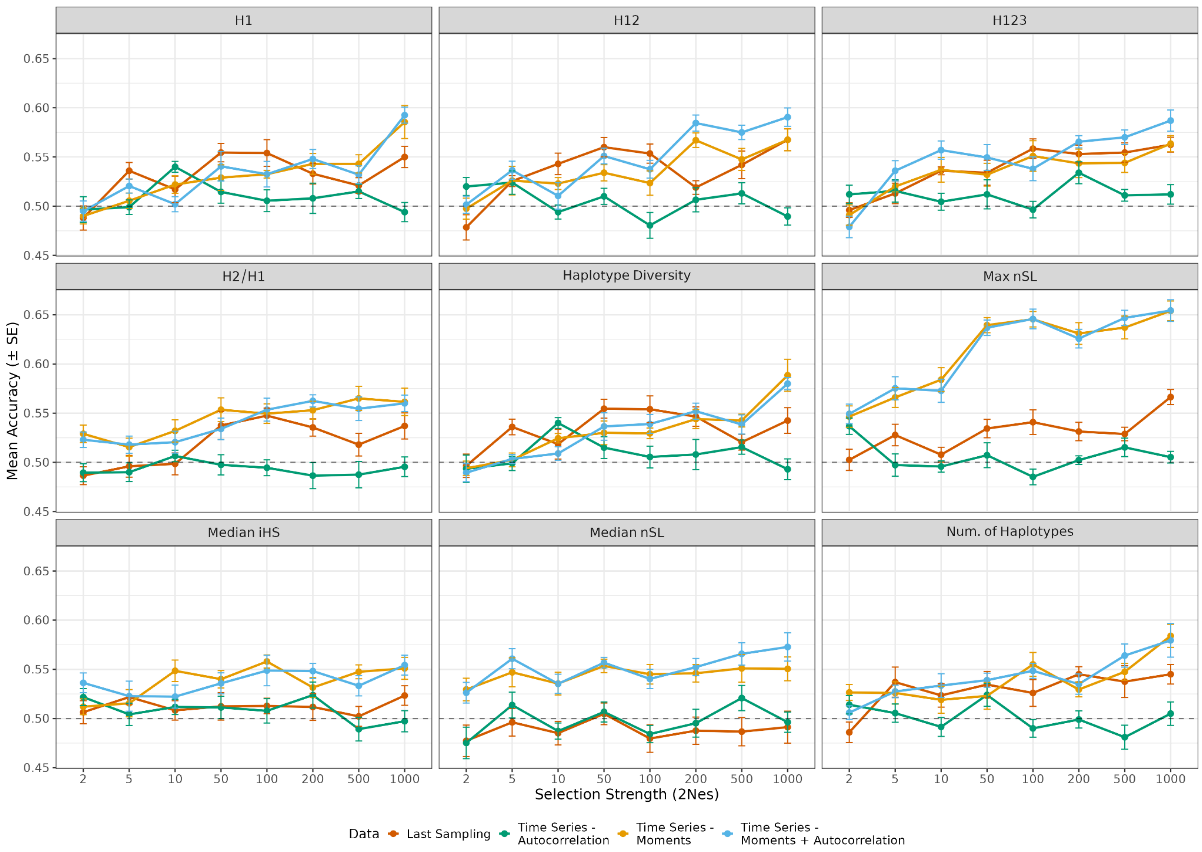


**Supplementary Figure 36.** RF classification accuracy for discriminating balancing selection from neutral evolution without segregation conditioning across selection strengths ($2N_es$) using summary statistics based on haplotype-level information and estimated for downsampled data ($n$=20 haplotypes per sampling point). Each panel shows results for a single haplotype-based statistic (panel titles), evaluated using four different feature sets derived from the same simulations: last sampling, autocorrelation, moments, and moments with autocorrelation. Points represent mean classification accuracy across the 10 cross-validation folds, with error bars indicating ± SE. The dashed line denotes random expectation (accuracy = 0.5).

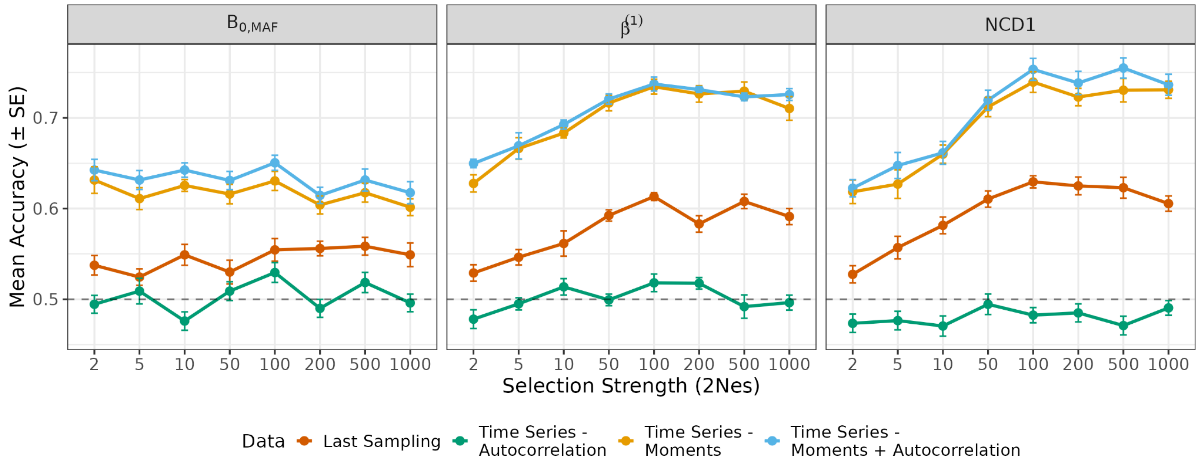


**Supplementary Figure 37.** RF classification accuracy for discriminating balancing selection from neutral evolution without segregation conditioning across selection strengths ($2N_es$) using summary statistics specifically designed to detect balancing selection and estimated for downsampled data ($n$=20 haplotypes per sampling point). Each panel shows results for a single balancing selection statistic (panel titles), evaluated using four different feature sets derived from the same simulations: last sampling, autocorrelation, moments, and moments with autocorrelation. Points represent mean classification accuracy across the 10 cross-validation folds, with error bars indicating ± SE. The dashed line denotes random expectation (accuracy = 0.5).

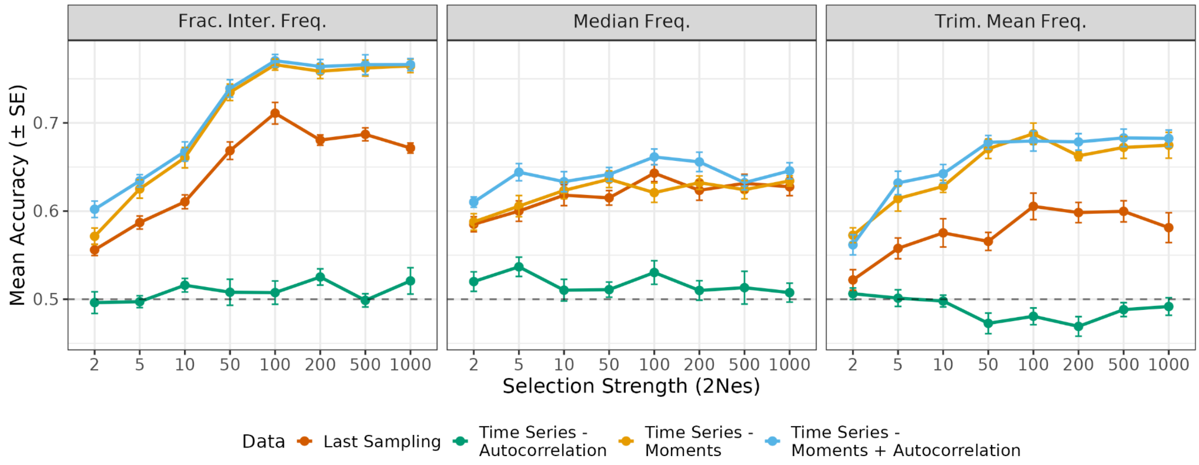


**Supplementary Figure 38.** RF classification accuracy for discriminating balancing selection from neutral evolution without segregation conditioning across selection strengths ($2N_es$) using summary statistics based on frequency information and estimated for downsampled data ($n$=20 haplotypes per sampling point). Each panel shows results for a single frequency-based statistic (panel titles), evaluated using four different feature sets derived from the same simulations: last sampling, autocorrelation, moments, and moments with autocorrelation. Points represent mean classification accuracy across the 10 cross-validation folds, with error bars indicating ± SE. The dashed line denotes random expectation (accuracy = 0.5).

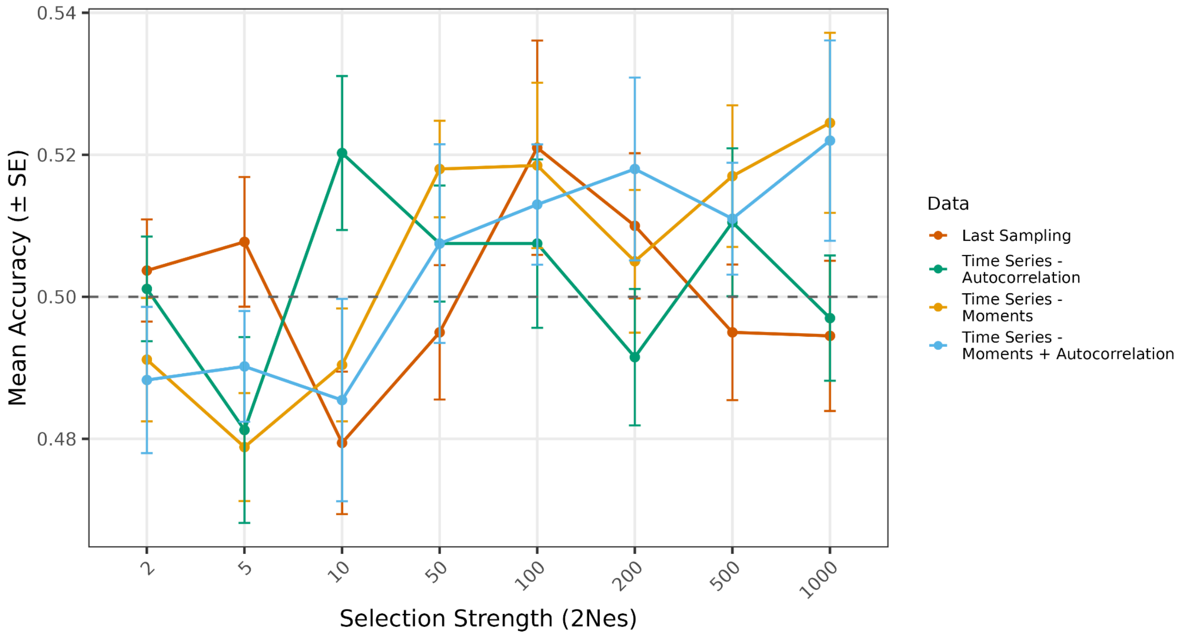


**Supplementary Figure 39.** RF classification accuracy for distinguishing overdominance from NFDS as a function of selection strength ($2N_es$). Models were trained using all 23 summary statistics and four different feature sets derived from the same simulations: last sampling, autocorrelation, moments, and moments with autocorrelation. Summary statistics were calculated using $n$=200 haplotypes per sampling point. Points represent mean classification accuracy across the 10 cross-validation folds, with error bars indicating ± SE. The dashed line denotes random expectation (accuracy = 0.5).

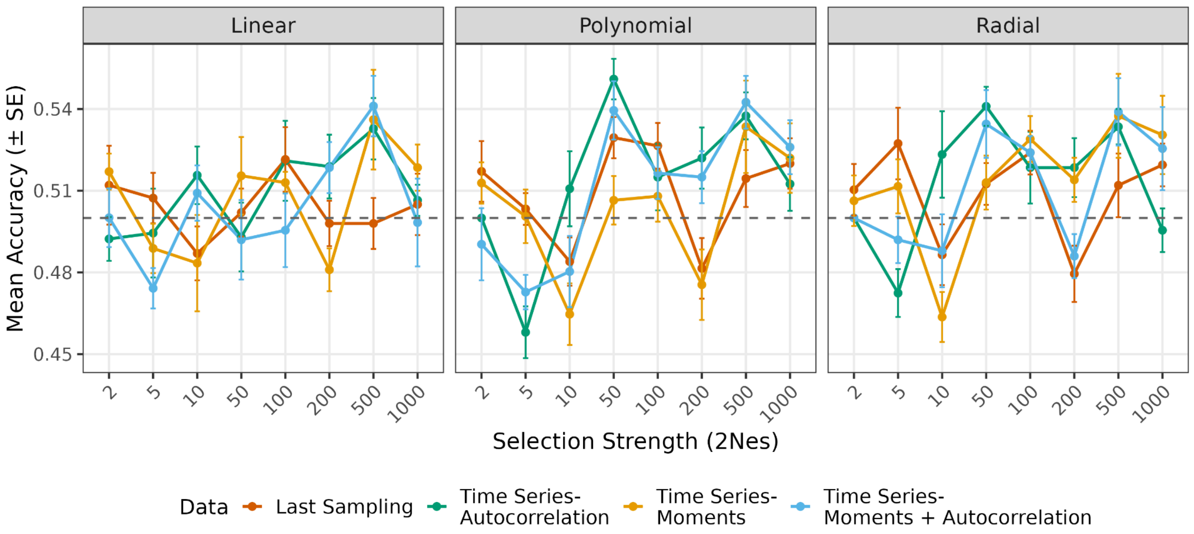


**Supplementary figure 40.** SVM classification accuracy for distinguishing overdominance from NFDS as a function of selection strength ($2N_es$). Models were trained using all 23 summary statistics and four different feature sets derived from the same simulations: last sampling, autocorrelation, moments, and moments with autocorrelation. Summary statistics were calculated using $n$=200 haplotypes per sampling point. Points represent mean classification accuracy across the 10 cross-validation folds, with error bars indicating ± SE. Each panel corresponds to SVM analyses with different kernels. The dashed line denotes random expectation (accuracy = 0.5).

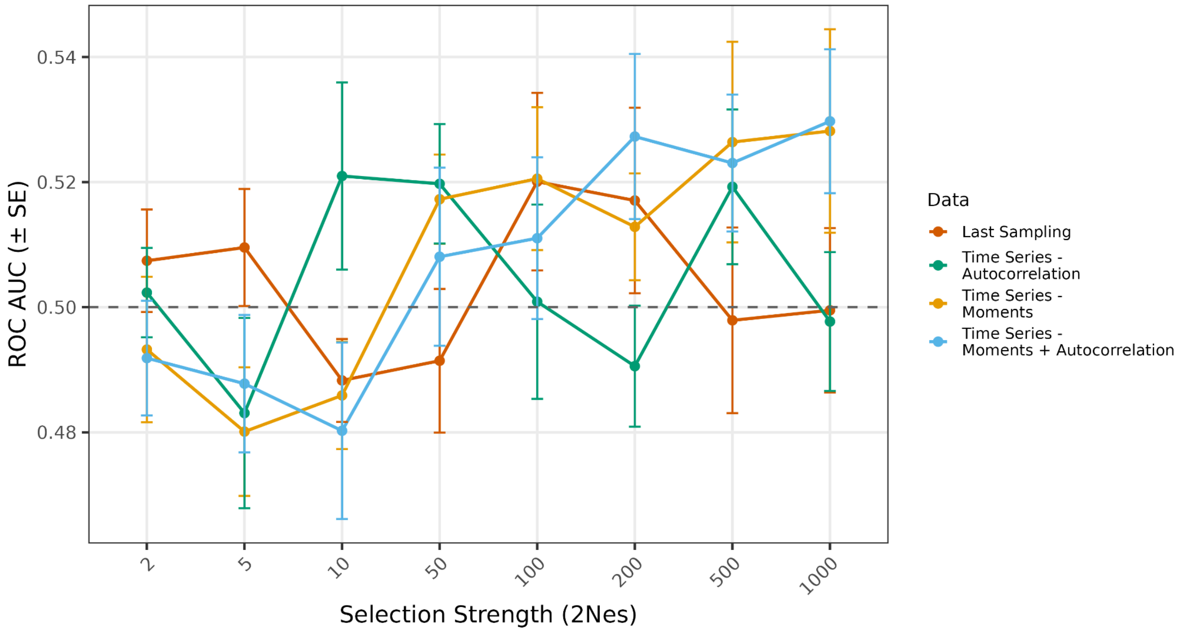


**Supplementary Figure 41.** RF classification performance, measured by ROC AUC, for distinguishing overdominance from NFDS as a function of selection strength ($2N_es$). Models were trained using all 23 summary statistics and four different feature sets derived from the same simulations: last sampling, autocorrelation, moments, and moments with autocorrelation. Summary statistics were calculated using $n$=200 haplotypes per sampling point. Points represent mean ROC AUC across the 10 cross-validation folds, with error bars indicating ± SE. The dashed line denotes random expectation (ROC AUC = 0.5).

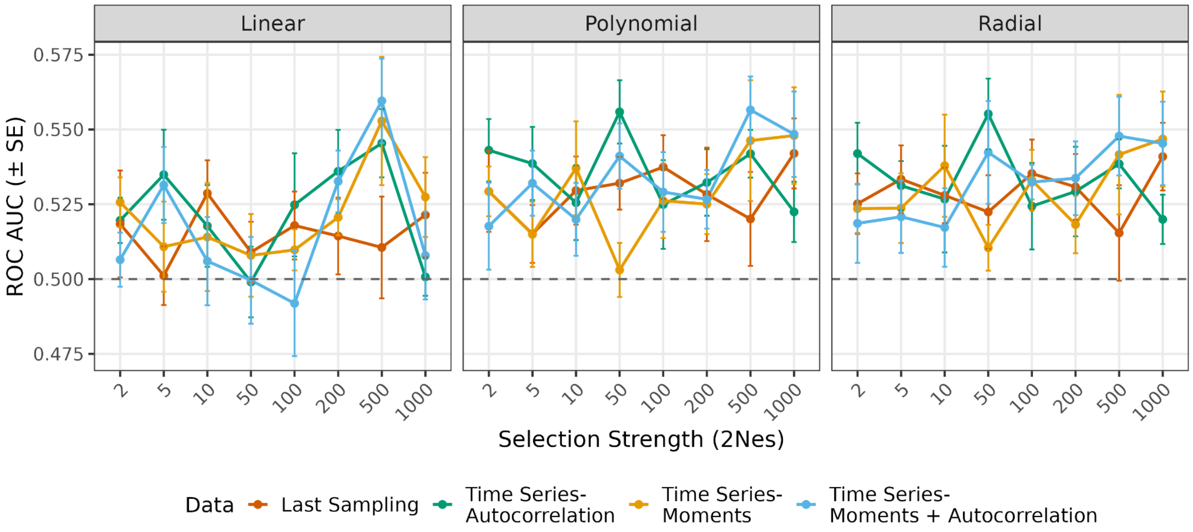


**Supplementary Figure 42.** SVM classification performance, measured by ROC AUC, for distinguishing overdominance from NFDS as a function of selection strength ($2N_es$). Models were trained using all 23 summary statistics and four different feature sets derived from the same simulations: last sampling, autocorrelation, moments, and moments with autocorrelation. Summary statistics were calculated using *n*=200 haplotypes per sampling point. Points represent mean ROC AUC across the 10 cross-validation folds, with error bars indicating ± SE. Each panel corresponds to SVM analyses with different kernels. The dashed line denotes random expectation (ROC AUC = 0.5).

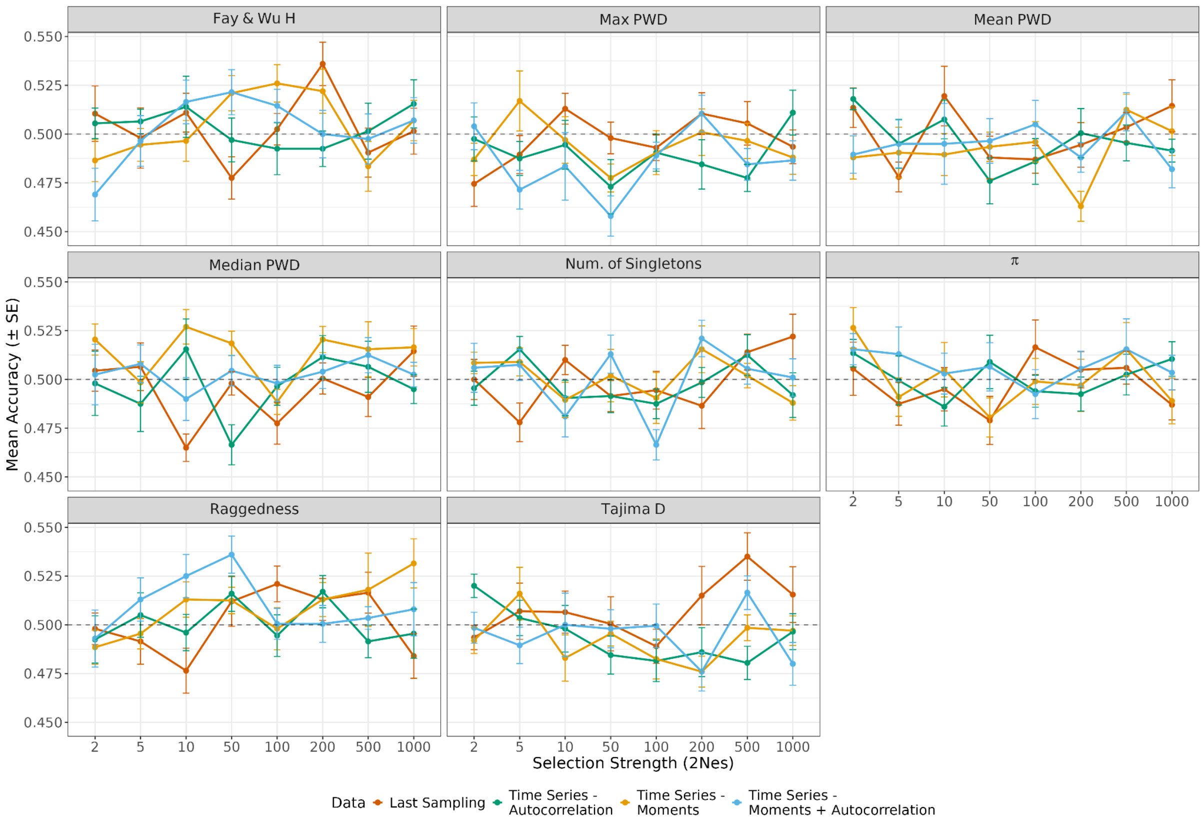


**Supplementary Figure 43.** RF classification accuracy for discriminating overdominance from NFDS across selection strengths ($2N_es$) using summary statistics derived from the site frequency spectrum (SFS). Each panel shows results for a single SFS-based statistic (panel titles), evaluated using four different feature sets derived from the same simulations: last sampling, autocorrelation, moments, and moments with autocorrelation. Summary statistics were calculated using $n$=200 haplotypes per sampling point. Points represent mean classification accuracy across the 10 cross-validation folds, with error bars indicating ± SE. The dashed line denotes random expectation (accuracy = 0.5).

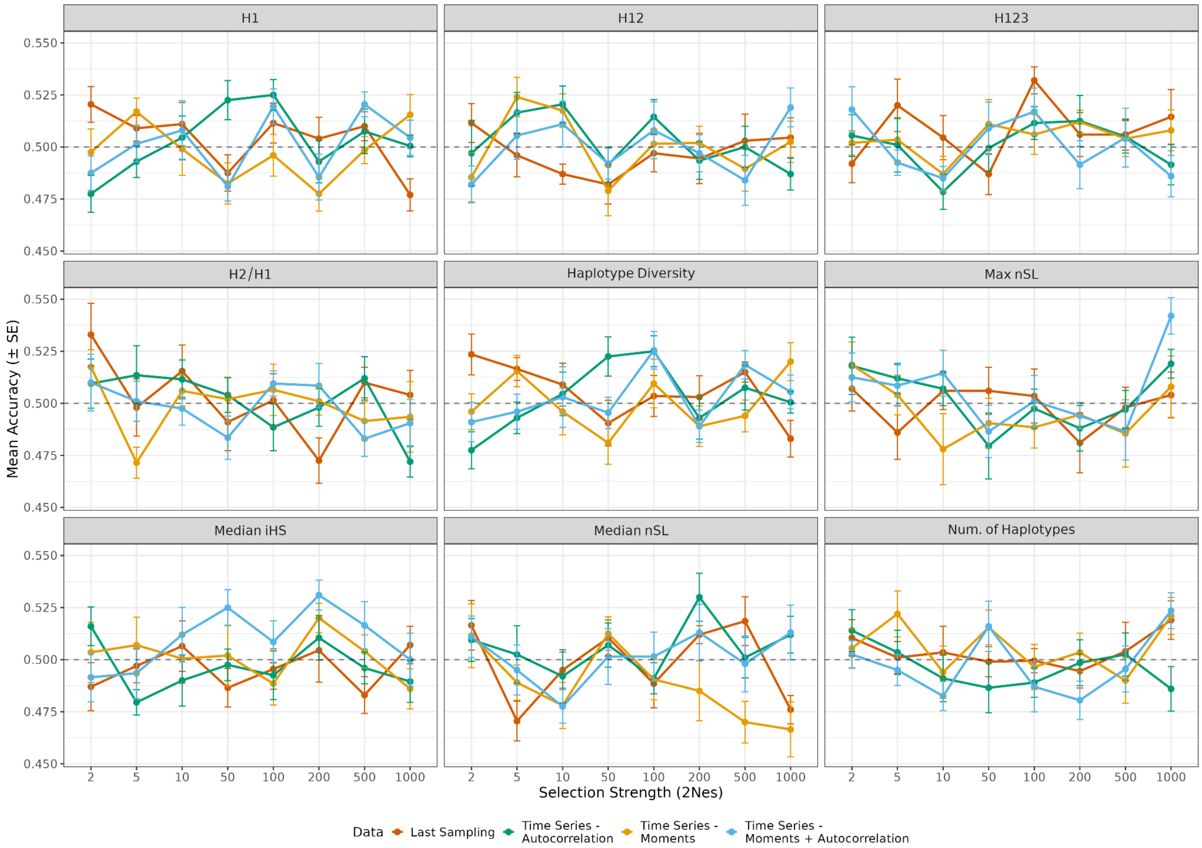


**Supplementary Figure 44.** RF classification accuracy for discriminating overdominance from NFDS across selection strengths ($2N_es$) using summary statistics based on haplotype-level information. Each panel shows results for a single haplotype-based statistic (panel titles), evaluated using four different feature sets derived from the same simulations: last sampling, autocorrelation, moments, and moments with autocorrelation. Summary statistics were calculated using $n$=200 haplotypes per sampling point. Points represent mean classification accuracy across the 10 cross-validation folds, with error bars indicating ± SE. The dashed line denotes random expectation (accuracy = 0.5).

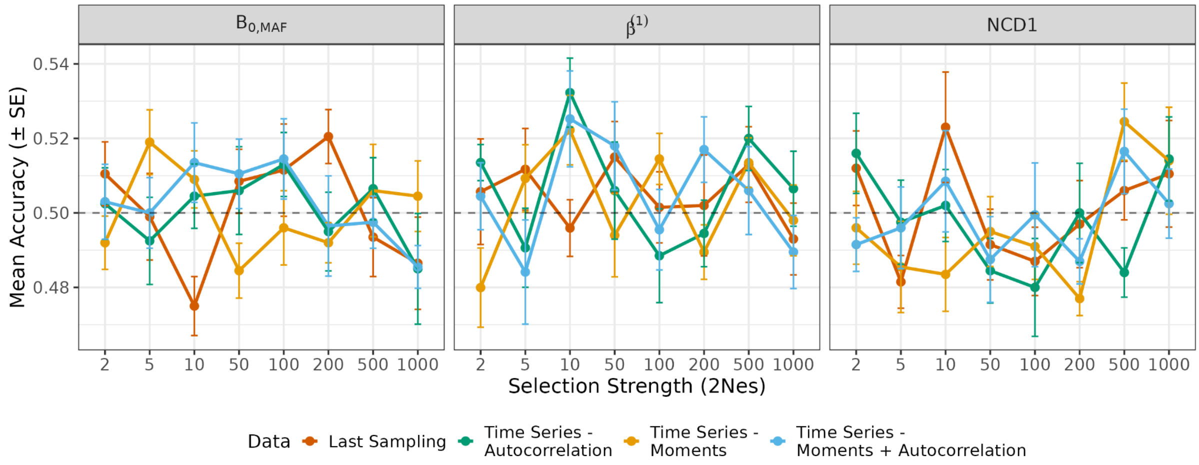


**Supplementary Figure 45.** RF classification accuracy for discriminating overdominance from NFDS across selection strengths ($2N_es$) using summary statistics specifically designed to detect balancing selection. Each panel shows results for a single balancing selection statistic (panel titles), evaluated using four different feature sets derived from the same simulations: last sampling, autocorrelation, moments, and moments with autocorrelation. Summary statistics were calculated using $n$=200 haplotypes per sampling point. Points represent mean classification accuracy across the 10 cross-validation folds, with error bars indicating ± SE. The dashed line denotes random expectation (accuracy = 0.5).

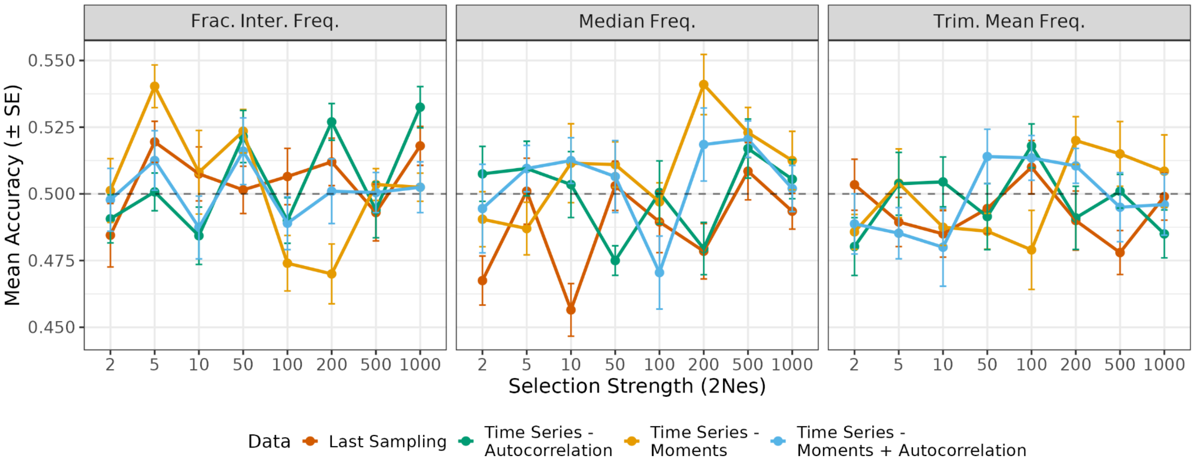


**Supplementary Figure 46.** RF classification accuracy for discriminating overdominance from NFDS across selection strengths ($2N_es$) based on frequency information. Each panel shows results for a single frequency-based statistic (panel titles), evaluated using four different feature sets derived from the same simulations: last sampling, autocorrelation, moments, and moments with autocorrelation. Summary statistics were calculated using $n$=200 haplotypes per sampling point. Points represent mean classification accuracy across the 10 cross-validation folds, with error bars indicating ± SE. The dashed line denotes random expectation (accuracy = 0.5).

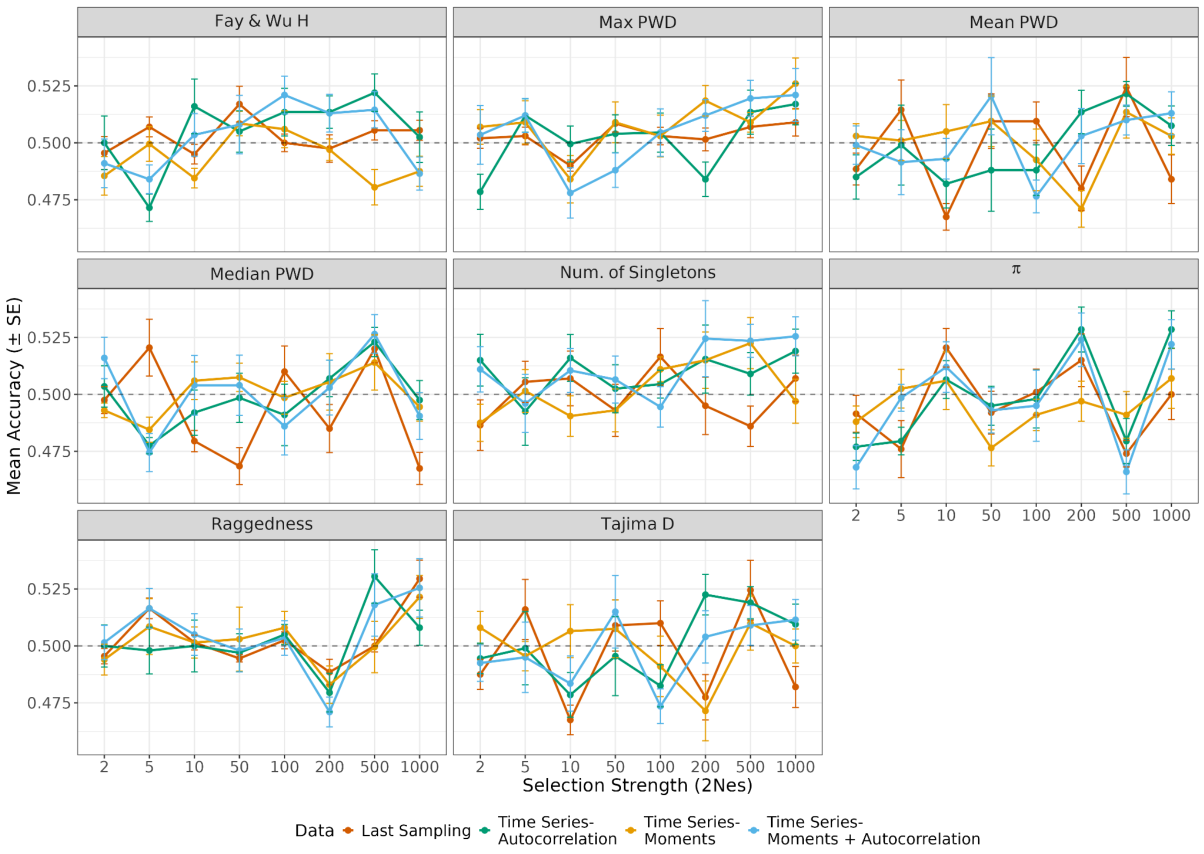


**Supplementary Figure 47.** Linear SVM classification accuracy for discriminating overdominance from NFDS across selection strengths ($2N_es$) using summary statistics derived from the site frequency spectrum (SFS). Each panel shows results for a single SFS-based statistic (panel titles), evaluated using four different feature sets derived from the same simulations: last sampling, autocorrelation, moments, and moments with autocorrelation. Summary statistics were calculated using $n$=200 haplotypes per sampling point. Points represent mean classification accuracy across the 10 cross-validation folds, with error bars indicating ± SE. The dashed line denotes random expectation (accuracy = 0.5).

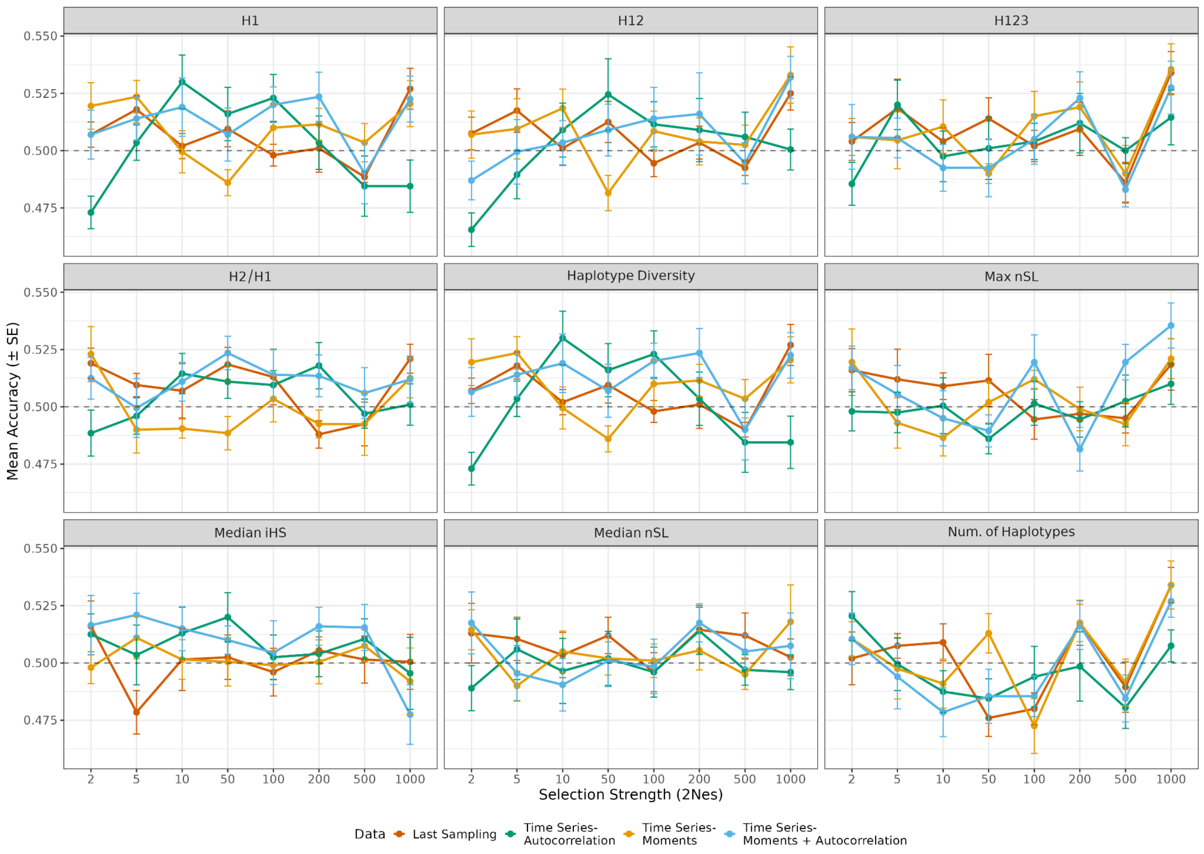


**Supplementary Figure 48.** Linear SVM classification accuracy for discriminating overdominance from NFDS across selection strengths ($2N_es$) using summary statistics derived from the site frequency spectrum (SFS). Each panel shows results for a single SFS-based statistic (panel titles), evaluated using four different feature sets derived from the same simulations: last sampling, autocorrelation, moments, and moments with autocorrelation. Summary statistics were calculated using $n$=200 haplotypes per sampling point. Points represent mean classification accuracy across the 10 cross-validation folds, with error bars indicating ± SE. The dashed line denotes random expectation (accuracy = 0.5).

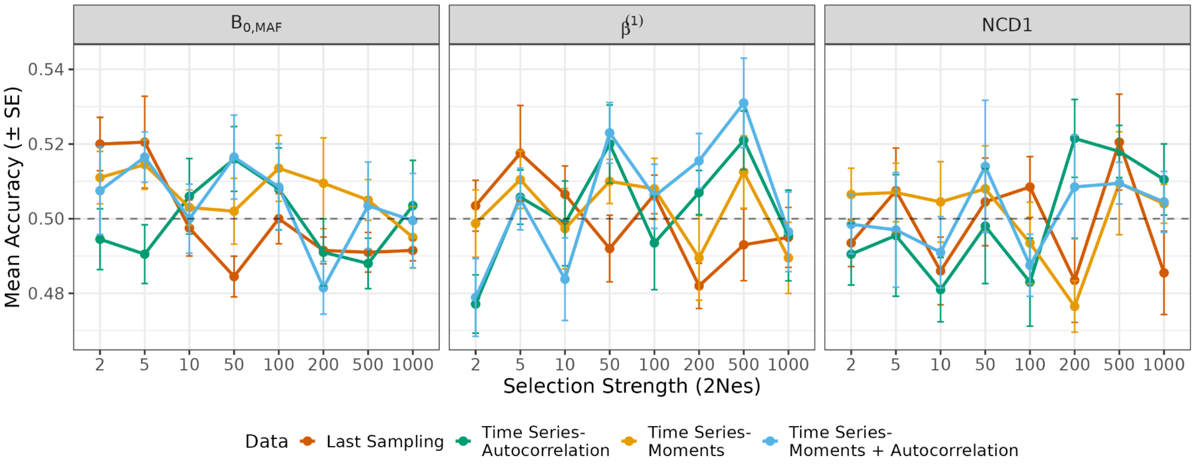


**Supplementary Figure 49.** Linear SVM classification accuracy for discriminating overdominance from NFDS across selection strengths ($2N_es$) using summary statistics specifically designed to detect balancing selection. Each panel shows results for a single balancing selection statistic (panel titles), evaluated using four different feature sets derived from the same simulations: last sampling, autocorrelation, moments, and moments with autocorrelation. Summary statistics were calculated using $n$=200 haplotypes per sampling point. Points represent mean classification accuracy across the 10 cross-validation folds, with error bars indicating ± SE. The dashed line denotes random expectation (accuracy = 0.5).

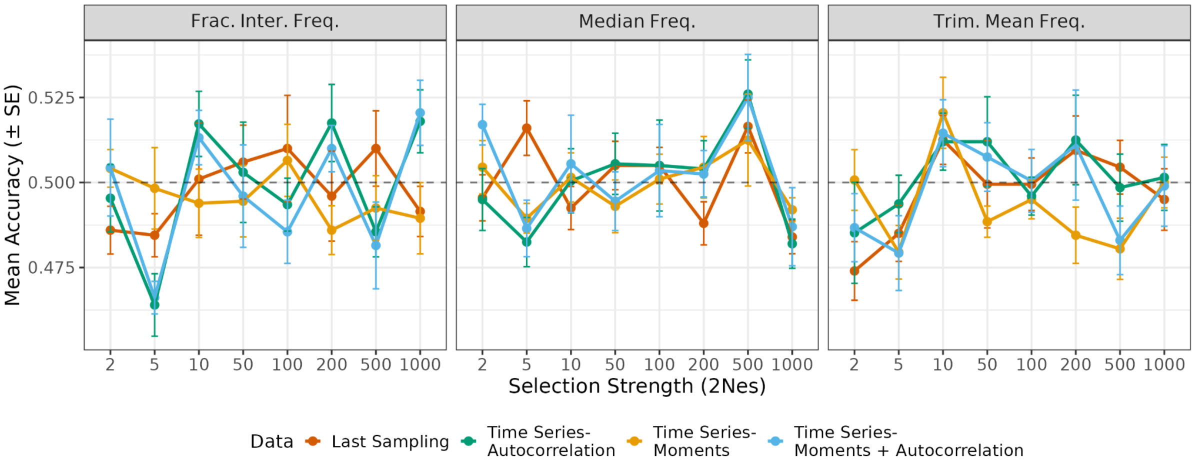


**Supplementary Figure 50.** Linear SVM classification accuracy for discriminating overdominance from NFDS across selection strengths ($2N_es$) using summary statistics based on frequency information. Each panel shows results for a single frequency-based statistic (panel titles), evaluated using four different feature sets derived from the same simulations: last sampling, autocorrelation, moments, and moments with autocorrelation. Summary statistics were calculated using $n$=200 haplotypes per sampling point. Points represent mean classification accuracy across the 10 cross-validation folds, with error bars indicating ± SE. The dashed line denotes random expectation (accuracy = 0.5).

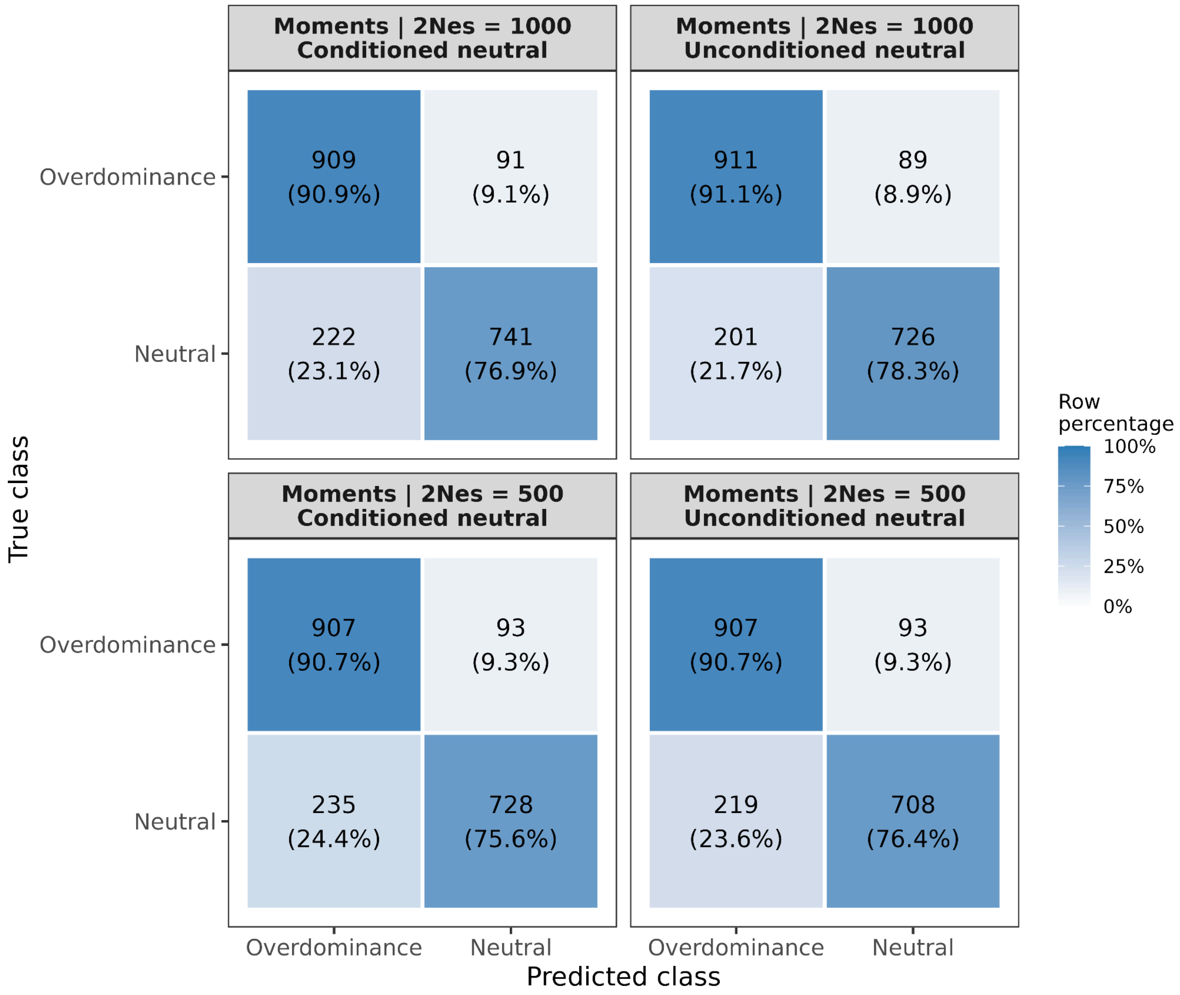


**Supplementary Figure 51.** RF confusion matrices for distinguishing overdominance from neutral evolution. Models were trained using the moments feature set constructed from all 23 summary statistics. Summary statistics were calculated using *n*=200 haplotypes per sampling point. Rows indicate the true class and columns the predicted class; cells show the number of observations and the corresponding percentage within each true class. Panels show results for selection strengths of $2N_es$ = 500 and $2N_es$ = 1000 under neutral simulations with and without segregation conditioning.

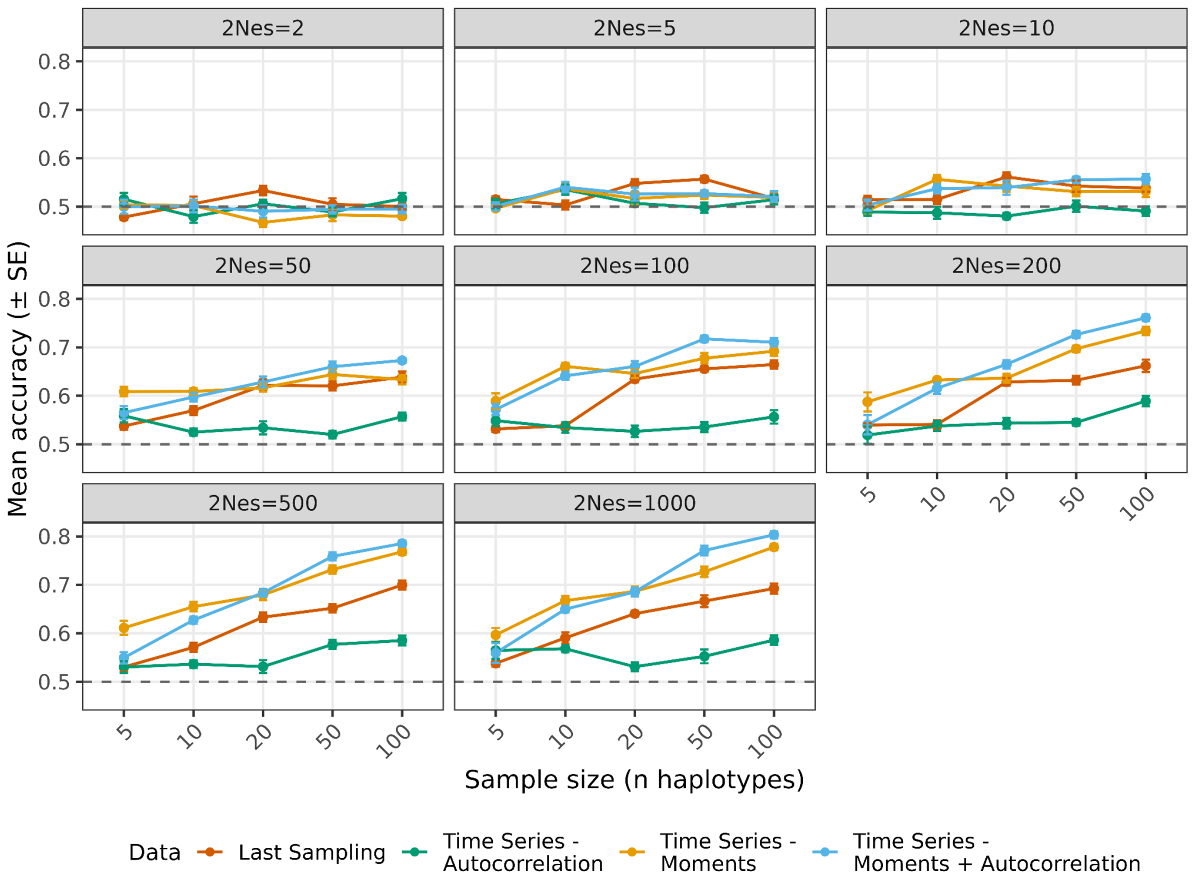


**Supplementary Figure 52.** RF classification accuracy for distinguishing balancing selection from neutral evolution, conditional on no fixation or loss, as a function of sample size (number of haplotypes). Models were trained using the four feature sets derived solely from the summary statistic Max PWD. The statistic is recalculated for randomly drawn samples of increasing sizes. Different panels show different selection strengths for overdominance and colors indicate different feature sets. Points represent the mean classification accuracy across 10 cross-validation folds, and error bars indicate ± SE. The dashed line denotes random expectation (accuracy = 0.5). Max PWD: Maximum per-site mean pairwise differences.
